# Critical Assessment of Protein Intrinsic Disorder Prediction

**DOI:** 10.1101/2020.08.11.245852

**Authors:** Marco Necci, Damiano Piovesan, CAID Predictors, DisProt Curators, Silvio C.E. Tosatto

## Abstract

Intrinsically disordered proteins defying the traditional protein structure-function paradigm represent a challenge to study experimentally. As a large part of our knowledge rests on computational predictions, it is crucial for their accuracy to be high. The Critical Assessment of protein Intrinsic Disorder prediction (CAID) experiment was established as a community-based blind test to determine the state of the art in predicting intrinsically disordered regions in proteins and the subset of disordered residues involved in binding other molecules. A total of 43 methods, 32 for disorder and 11 for binding regions, were evaluated on a dataset of 646 novel manually curated proteins from DisProt. The best methods use deep learning techniques and significantly outperform widely used earlier physicochemical methods across different types of targets. Disordered binding regions remain hard to predict correctly. Depending on the definition used, the top disorder predictor has an F_Max_ of 0.483 (*DisProt*) or 0.792 (*DisProt-PDB*). As the top binding predictor only attains an F_Max_ of 0.231, this suggests significant potential for improvement. Intriguingly, computing times among the top performing methods vary by up to four orders of magnitude.

## Introduction

Intrinsically disordered proteins and regions (IDPs/IDRs) that do not adopt a fixed three dimensional fold under physiological conditions are now well-established in structural biology^4^. Over the last two decades, there has been increasing evidence for the involvement of IDPs and IDRs in a variety of essential biological processes^5,6^ and molecular functions which complement those of globular domains^1,2^. As they are involved in many diseases, such as Alzheimer’s^7^, Parkinson’s^8^ and cancer^9^, they also represent promising novel targets for drug discovery^10,11^. Despite their importance, IDPs/IDRs are historically understudied due to the difficulties in measuring their dynamic behavior directly and because some of them are disordered only under particular environmental conditions, such as pH, PTMs, localization, and binding, i.e. their structural disorder is context dependent^3^. Experimental methods to detect intrinsic structural disorder (ID) include X-ray crystallography, nuclear magnetic resonance spectroscopy (NMR), small-angle X-ray scattering (SAXS), circular dichroism (CD) and Förster resonance energy transfer (FRET)^12–15^. Each technique provides a unique point of view on the phenomenon of ID and different types of experimental evidence give researchers insights over the functioning mechanisms of IDPs, such as flexibility, folding-upon-binding, conformational heterogeneity.

An accumulation of experimental evidence has corroborated the early notion that ID can be inferred from sequence features, first suggested by R J Williams^16^. Since then, dozens of ID prediction methods based on different principles and computing techniques have been published^17^, such as VSL2B^18^, DisEMBL^19^, DISOPRED^20^, IUPred^21^, Espritz^22^ and many others. Coordinates of intrinsically disordered regions (IDRs) and annotations related to their function both predicted and derived from experimental evidence are stored in a variety of dedicated databases, each focusing on particular aspects of the ID spectrum, namely DisProt^23^, MobiDB^24^, IDEAL^25^, DIBS^26^ and MFIB^27^. Since over the last few years, IDR coordinate annotations are also provided in some of the most important core data resources like InterPro^28^, UniProt^29^ and PDBe^30^.

The large variety of ID predictors available makes it difficult to compare them, confounding biologists wanting to make an informed choice to find the best performers. Similarly, binding predictions are widely used but an assessment of these predictors has never been performed and is highly needed in the field. Since many IDPs are interaction hubs^6^, this makes their binding regions particularly challenging to predict and a good benchmark. In this report, we describe the first edition of CAID, a bi-annual experiment inspired by the Critical Assessment of protein Structure Prediction (CASP) for the benchmarking of ID and binding predictors on a community curated dataset of 646 novel proteins obtained from DisProt^23^. CAID is the first experiment of its kind and is bound to set a new quality standard in the field.

## Results

Given a new protein sequence, the task of an IDR prediction software is to assign a score to each residue describing its tendency of being intrinsically disordered at any stage of the protein life. In CAID, we evaluate both the accuracy of prediction methods and technical aspects related to software implementation like their speed, which has a direct impact on carrying out large-scale analyses.

CAID was organized as follows (**Fig. 1A**). Participants submitted their implemented prediction software to the assessors and during the submission phase, the authors provided support to install and test execution on MobiDB servers. After the submission deadline, the assessors ran the packages and generated predictions for a set of proteins whose disorder annotations were not available at the training time.

**Figure 1.**
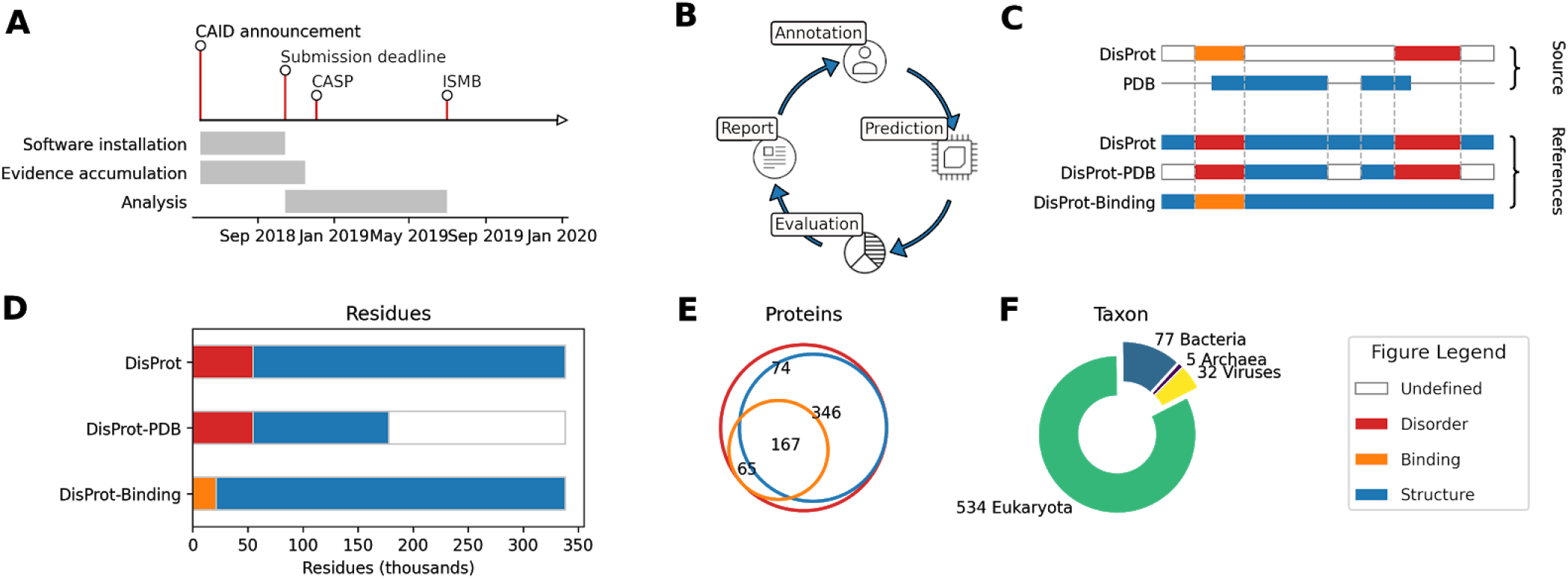
**(A)** CAID timeline: phases of CAID from June 2018 to today. **(B)** CAID process: Iterative process of the CAID experiment in 4 phases: (i) Annotation: any process that produces unpublished annotation of IDR coordinates, in this edition annotation refers to the DisProt round of annotation. (ii) Prediction: annotations are used to build references with which we test predictors (iii) Evaluation: Predictions are evaluated (iv) Report: A report of the evaluation is produced and published on peer-reviewed journals and to a web page that will allow to browse the evaluation of all CAID editions. **(C)** Residue classification strategy for the *DisProt* and *DisProt-PDB* references. **(D)** The number of residues for each class in different references. **(E)** The number of proteins for each set of annotations that they contain. **(F)** The number of proteins each taxon **(G)** Positive label (disorder/binding) fraction in the dataset (left), distribution of protein length (middle) and region length (right) in the different reference sets.

Structural properties of proteins can be studied by a number of different experimental techniques, some giving direct, some indirect evidence of disorder. Different techniques are biased in different ways, for example, IDRs inferred from missing residues in X-ray experiments are generally shorter because longer non-crystallizable IDRs are either excised when preparing the construct or are detrimental to the crystallization of the protein. On the contrary, circular dichroism can detect the absence of fixed structure in the full protein but does not provide any information about IDR coordinates. IDR annotations are more reliable when confirmed by multiple lines of independent and different experimental evidence.

In this first round of CAID, we selected the DisProt database as the reference for structural disorder because it provides a large number of manually curated disorder annotations at the protein level, with the majority of residues annotated with more than one experiment^23^. DisProt annotates disordered regions of at least 10 residues in length to evaluate IDRs likely to be associated with a biological function by excluding short loops, i.e. segments connecting secondary structure elements. DisProt also contains protein-protein interaction interfaces which fall into disordered regions. These binding regions were used as a separate dataset (DisProt-Binding).

In an ideal situation, all DisProt annotations would be complete, i.e. each protein in DisProt would be annotated with all the disordered (or binding) regions it contains under physiological conditions. If this were true, we could simply consider all residues to be structured (i.e. negatives) if they are not annotated as disordered (i.e. positives). Since not all IDRs are in DisProt yet, we created the DisProt-PDB dataset, in which negatives are restricted by PDB observed residues (**Fig. 1C**). This dataset is more conservative but can be considered more reliable as it excludes “uncertain” residues, which have neither structural nor disorder annotation. Compared to DisProt, DisProt-PDB is more similar to datasets used to train some of the disorder predictors (e.g. in^19,20,22^) and for CASP disorder challenges^31^.

The distribution of organisms reflects what is known from other studies^1,2^ with the majority of ID targets coming from eukaryotes with a good representation of viruses and bacteria, and much fewer from archaea (**Fig. 1F**). At the species level, annotations are strongly biased in favor of model organisms, with the majority of cases coming from human, mouse, rat, *Escherichia coli* and a few other common model organisms (see Supplementary Figure 6). Target proteins are not redundant at the sequence level and are different from known examples available in the previous DisProt release, with a mean sequence identity of 22.2% against the previous DisProt release and 17.1% within the dataset (see Supplementary Figure 3). CAID has two main categories: the prediction of intrinsic structural disorder and the predictions of binding sites found in IDRs.

### IDR prediction performance

The quality of IDR prediction can be evaluated in different ways depending on the scope of studying structural disorder. In some cases, it is relevant to know the fraction of disorder; in other cases, it is more important to know the exact position of the IDR in the sequence. Since disorder can be used as a proxy to estimate the complexity of an organism or to complement a sequence search, it is also important for a predictor to be fast enough to be applied at the genome-scale. For CAID, we report a simple metric, the maximum F1-score (*F_max_*; i.e. maximum harmonic mean between precision and recall across all thresholds), which takes into account predictions across the entire spectrum of sensitivity, from high to low^3^.

The performance of top methods, based on *F_max_* and calculated over all targets, are shown in **Figure 2** for the DisProt and DisProt-PDB datasets. The F1-score, which has the advantage of being insensitive to dataset imbalance (**Fig. 1D**), provides a ranking almost identical to the one obtained with the Matthew’s Correlation Coefficient (MCC). See Supplementary Figures 12, 13, 33, 34 for a full comparison and the dependence of F1-score and MCC on the predictor confidence score, along with the predictor default confidence threshold (Supplementary Figures 10, 11, 30, 31). All methods were compared with various baselines described in the “Online Methods”. In some applications, the objective was to predict which protein fragments of the protein are disordered, based on known examples in the PDB. However, this was a different problem than predicting functional IDRs, for instance, aiming to evaluate their biophysical properties. The naive baselines help us understand this difference and assess how effective the transfer-by-homology of structural information can be for IDR prediction (see Discussion). In the *PDB Observed* baseline, mimicking perfect knowledge, all residues not covered by any PDB structure are labelled as disordered. Alternatively, in the *Gene3D* baseline, residues are considered disordered if they do not match any Gene3D prediction for homologous domains. In the Shuffled-Dataset baseline, the reference is randomly shuffled at the dataset level, while Random is an actual random predictor which does not use any prior knowledge.

**Figure 2.**
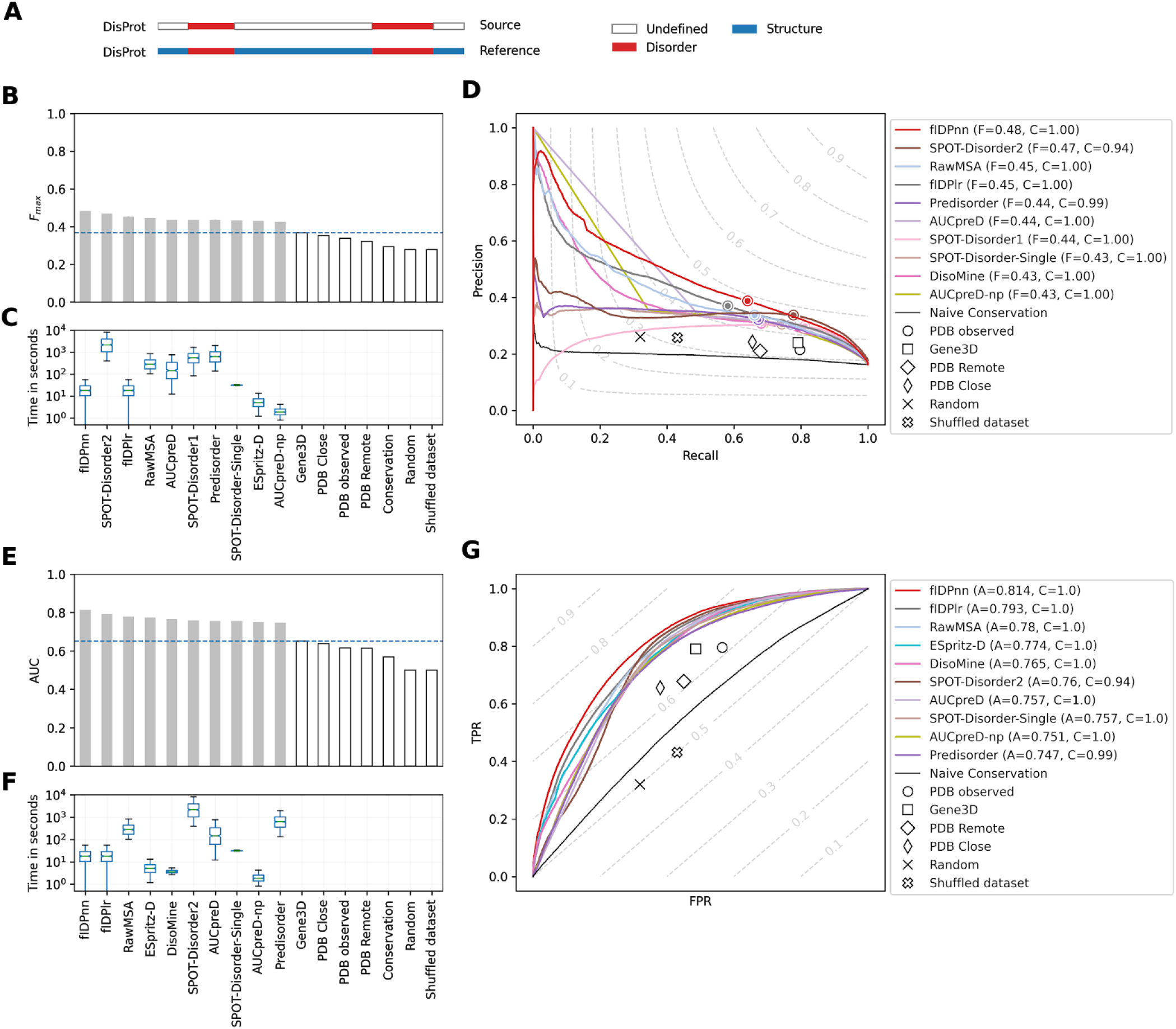
Prediction success and CPU times for the ten top-ranking disorder predictors in the *DisProt* dataset. Reference used (*DisProt*) in the analysis and how it is obtained (panel **A**). Performance of predictors expressed as maximum F1-Score across all thresholds (F_max_) (panel **B**) and AUC (panel **E**) for the top ten best ranking methods (light gray) and baselines (white) and the distribution of execution time per-target (panels **C, F**) using *DisProt* dataset. The horizontal line in panels B, E indicates the F_max_ and AUC of the best baseline, respectively. Precision-Recall (panel **D**) and ROC curves (panel **G**) of ten top-ranking methods and baselines using *DisProt* dataset, with level curves of the F1-Score and Balanced accuracy, respectively. Magenta dots on panels C, F indicate that the whole distribution of execution-times is lower than 1 second.

Both the *F_max_* (**Fig.s 2B, 2D**) and Area under the ROC curve (AUC) (**Fig.s 3E, 3G**) measures are substantially different when predictors are tested on the DisProt dataset, which contains “uncertain” residues, and when tested on the DisProt-PDB dataset. *PDB Observed* baseline, by definition, cannot predict negative residues outside PDB regions. Therefore, it mainly generates false positives (56.5%), which drop to zero when considering the DisProt-PDB dataset, where the “uncertain” residues are completely filtered out. IDRs overlapping with PDB regions, usually corresponding to residues that take part in folding-upon-binding events, instead generate false negatives. These are far less common (20.4%) and remain the same for the two datasets. The *Gene3D* baseline typically increases PDB coverage (negatives) exploiting the transfer-by-homology principle. As a consequence, the probability of false positives is lower (48.6%), and false negatives are only marginally more frequent (20.9%). For the DisProt dataset, *Gene3D* slightly outperforms *PDB Observed* both in terms of *F_max_* (**Fig. 2 panels B,D**) and AUC (**Fig. 3 panels E,G**). Instead, for the DisProt-PDB dataset, *PDB Observed* is significantly superior to all methods with only 6.3% of mispredicted residues, all false negatives.

**Figure 3.**
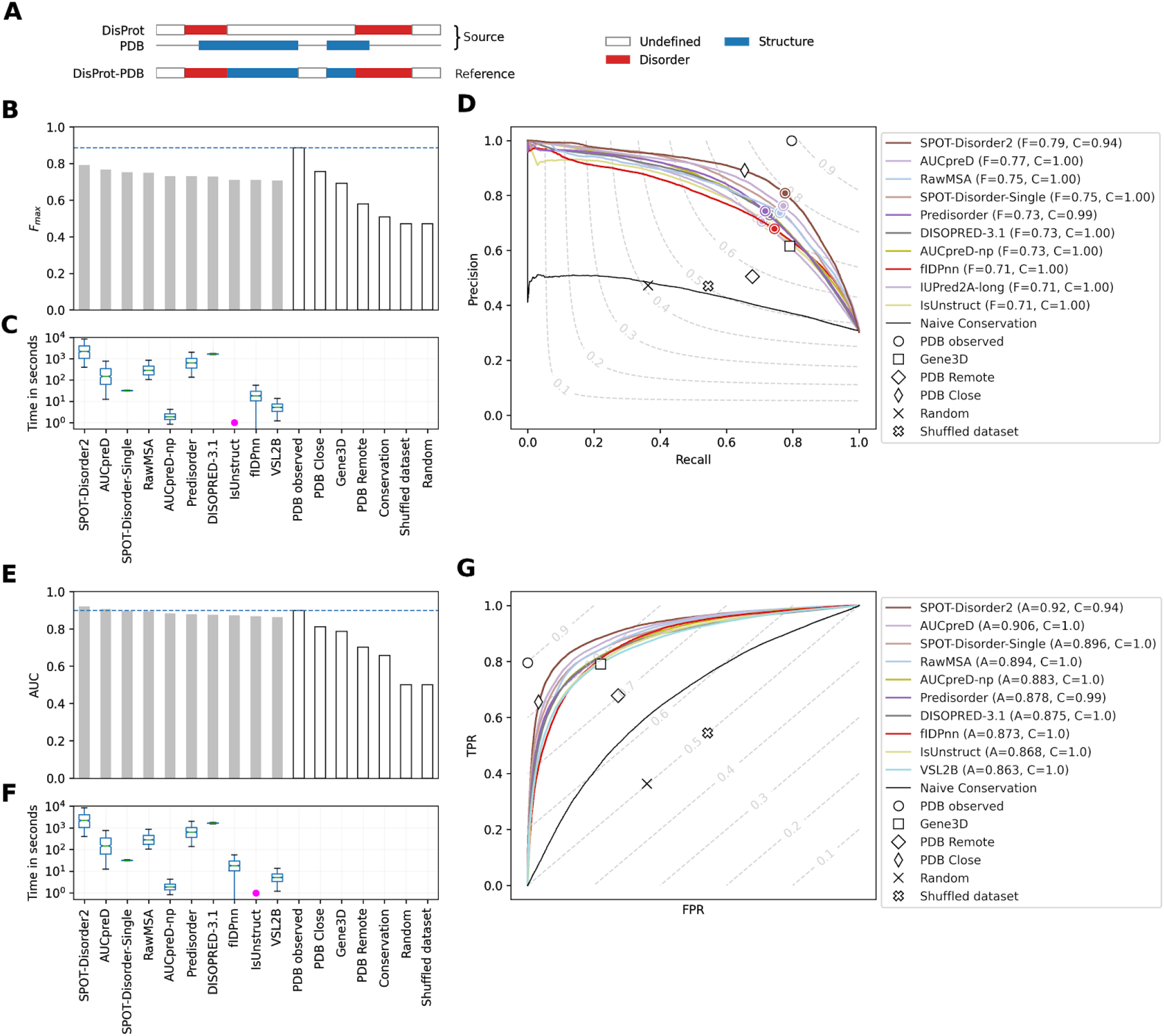
Prediction success and CPU times for the ten top-ranking disorder predictors in the *DisProt-PDB* dataset. Reference used (*DisProt-PDB*) in the analysis and how it is obtained (panel **A**). Performance of predictors expressed as maximum F1-Score across all thresholds (F_max_) (panel B) and AUC (panel **E**) for the top ten best ranking methods (light gray) and baselines (white) and the distribution of execution time per-target (panels **C, F**) using *DisProt-PDB* dataset. The horizontal line in panels B, E indicates the F_max_ and AUC of the best baseline, respectively. Precision-Recall (panel **D**) and ROC curves (panel **G**) of ten top-ranking methods and baselines using *DisProt-PDB* dataset, with level curves of the F1-Score and Balanced accuracy, respectively. Magenta dots on panels C, F indicate that the whole distribution of execution-times is lower than 1 second

Given the relevance of the host organism for determining environmental factors for IDPs such as temperature, we wondered whether predictor performance would be affected in different subsets. For this reason, performance has also been assessed separately for mammalian and prokaryotic proteins (see Supplementary Figures 19-28 for the *DisProt* and 40-49 for the *DisProt-PDB dataset*). The ranking changes only slightly and after the top two positions. Performance for mammalian sequences is ca. 0.05 and 0.03 lower in terms of *F_max_* and AUC for all methods, suggesting this to be a somewhat harder challenge.

Across the different performance measures, several methods can be consistently found in the top five: These are SPOT-Disorder2, fIDPnn, RawMSA and AUCpreD. While the ordering changes for different measures and reference sets, and the differences among them are not statistically significant (see Supplementary Figures 17-18, 23, 28, 38-39, 44 and 49), these methods can be broadly seen as performing consistently well. Looking at the Precision-Recall curves (**Fig. 2 panel D**), we notice that the top five methods (excluding fIDPnn/lr in the DisProt dataset and AUCpred-np in the DisProt-PDB dataset) leverage evolutionary information, introducing a database search as a preliminary step. The performance gain, which is on average 4.5% in terms of F_Max_, comes at the cost of slowing down the prediction by two to four orders of magnitude (**Fig. 2 panel C**, see also Supplementary Figures 4, 12-14 and 33-35).

### Fully disordered proteins

Within the challenge of disorder prediction, we separately considered the special category of fully disordered proteins (IDPs). These proteins are interesting as they are particularly challenging to investigate experimentally, e.g. they cannot be probed with X-ray crystallography, yet they are of great interest as they fulfil unique biological functions^2,32^. We therefore designed another challenge, in which the task is to tell apart fully from not-fully disordered proteins. In this challenge, predictors are asked to identify entire proteins considered fully disordered, i.e. the evaluation is performed on entire proteins instead of single residues. We consider a fully disordered protein, those with at least 95% of residues predicted or annotated as disordered. According to this definition, the number of fully disordered proteins in the DisProt dataset is 40 out of 646. Different threshold values did not significantly affect the ranking (see Supplementary Material Tables 6, 7 and 8). In **Table 1** all methods are sorted based on F1-Score. False positives are limited for many methods, although correct IDP predictions are generally made for less than half of the dataset. This suggests room for improvement, as fully disordered proteins should be easier to predict from sequence. Methods using secondary structure information may be penalized for IDP prediction, as annotations frequently rely on detection methods without residue level resolution (e.g. circular dichroism, see Supplementary Figure 7).

**Table 1:** Confusion matrix and metrics for the prediction of fully disordered proteins on *DisProt* dataset. TP (True Positives count), FP (False Positives count), FN (False Negatives count), TN (True Negatives count), MCC (Matthews correlation coefficient), F1-S (F1-Score), TNR (True Negative Rate, Specificity), TPR (True Positive Rate, Recall), PPV (Positive Predictive Value, Precision) and BAC (Balanced Accuracy) for the prediction of fully disordered proteins. Proteins with disorder prediction or disorder annotation covering at least 95% of the sequence are considered fully disordered. Predictors are sorted by their F1-Score. Baseline names are in bold.

| Fully disordered proteins (ID fraction $\geq 95\%$) | | | | | | | | | | |
| --- | --- | --- | --- | --- | --- | --- | --- | --- | --- | --- |
|  | TN | FP | FN | TP | MCC | F1-S | TNR | TPR | PPV | BAC |
| fIDPnn | 585 | 16 | 19 | 26 | 0.569 | 0.598 | 0.973 | 0.578 | 0.619 | 0.776 |
| RawMSA | 582 | 19 | 19 | 26 | 0.546 | 0.578 | 0.968 | 0.578 | 0.578 | 0.773 |
| VSL2B | 578 | 23 | 22 | 23 | 0.468 | 0.505 | 0.962 | 0.511 | 0.500 | 0.736 |
| fIDPIr | 566 | 35 | 18 | 27 | 0.468 | 0.505 | 0.942 | 0.600 | 0.435 | 0.771 |
| Predisorder | 589 | 12 | 26 | 19 | 0.479 | 0.500 | 0.980 | 0.422 | 0.613 | 0.701 |
| SPOT-Disorder1 | 572 | 29 | 23 | 22 | 0.416 | 0.458 | 0.952 | 0.489 | 0.431 | 0.720 |
| DisoMine | 551 | 50 | 17 | 28 | 0.421 | 0.455 | 0.917 | 0.622 | 0.359 | 0.770 |
| AUCpreD | 588 | 13 | 28 | 17 | 0.431 | 0.453 | 0.978 | 0.378 | 0.567 | 0.678 |
| SPOT-Disorder2 | 574 | 27 | 24 | 21 | 0.409 | 0.452 | 0.955 | 0.467 | 0.438 | 0.711 |
| SPOT-Disorder-Single | 594 | 7 | 30 | 15 | 0.452 | 0.448 | 0.988 | 0.333 | 0.682 | 0.661 |
| IsUnstruct | 588 | 13 | 29 | 16 | 0.411 | 0.432 | 0.978 | 0.356 | 0.552 | 0.667 |
| IUPred2A-long | 595 | 6 | 32 | 13 | 0.420 | 0.406 | 0.990 | 0.289 | 0.684 | 0.639 |
| **Gene3D** | 505 | 96 | 10 | 35 | 0.391 | 0.398 | 0.840 | 0.778 | 0.267 | 0.809 |
| ESpritz-N | 597 | 4 | 33 | 12 | 0.426 | 0.393 | 0.993 | 0.267 | 0.750 | 0.630 |
| ESpritz-D | 555 | 46 | 23 | 22 | 0.342 | 0.389 | 0.923 | 0.489 | 0.324 | 0.706 |
| PyHCA | 596 | 5 | 33 | 12 | 0.411 | 0.387 | 0.992 | 0.267 | 0.706 | 0.629 |
| JRONN | 595 | 6 | 33 | 12 | 0.397 | 0.381 | 0.990 | 0.267 | 0.667 | 0.628 |
| MobiDB-lite | 599 | 2 | 34 | 11 | 0.437 | 0.379 | 0.997 | 0.244 | 0.846 | 0.621 |
| DisPredict-2 | 586 | 15 | 32 | 13 | 0.330 | 0.356 | 0.975 | 0.289 | 0.464 | 0.632 |
| IUPred2A-short | 599 | 2 | 35 | 10 | 0.413 | 0.351 | 0.997 | 0.222 | 0.833 | 0.609 |
| S2D-2 | 572 | 29 | 30 | 15 | 0.288 | 0.337 | 0.952 | 0.333 | 0.341 | 0.643 |
| **PDB observed** | 468 | 133 | 13 | 32 | 0.286 | 0.305 | 0.779 | 0.711 | 0.194 | 0.745 |
| AUCpreD-np | 590 | 11 | 35 | 10 | 0.293 | 0.303 | 0.982 | 0.222 | 0.476 | 0.602 |
| ESpritz-X | 595 | 6 | 36 | 9 | 0.321 | 0.300 | 0.990 | 0.200 | 0.600 | 0.595 |
| FoldUnfold | 456 | 145 | 14 | 31 | 0.256 | 0.281 | 0.759 | 0.689 | 0.176 | 0.724 |
| DISOPRED-3.1 | 596 | 5 | 39 | 6 | 0.246 | 0.214 | 0.992 | 0.133 | 0.545 | 0.563 |
| DisEMBL-HL | 601 | 0 | 41 | 4 | 0.288 | 0.163 | 1.000 | 0.089 | 1.000 | 0.544 |
| **PDB Remote** | 590 | 11 | 42 | 3 | 0.085 | 0.102 | 0.982 | 0.067 | 0.214 | 0.524 |
| DisEMBL-465 | 601 | 0 | 43 | 2 | 0.204 | 0.085 | 1.000 | 0.044 | 1.000 | 0.522 |
| **PDB Close** | 589 | 12 | 43 | 2 | 0.043 | 0.068 | 0.980 | 0.044 | 0.143 | 0.512 |
| **Conservation** | 441 | 160 | 38 | 7 | -0.064 | 0.066 | 0.734 | 0.156 | 0.042 | 0.445 |
| DynaMine | 601 | 0 | 45 | 0 | 0.000 | 0.000 | 1.000 | 0.000 | 0.000 | 0.500 |
| GlobPlot | 601 | 0 | 45 | 0 | 0.000 | 0.000 | 1.000 | 0.000 | 0.000 | 0.500 |
| DFLpred | 601 | 0 | 45 | 0 | 0.000 | 0.000 | 1.000 | 0.000 | 0.000 | 0.500 |

### Prediction of disordered binding sites

As a second major challenge, CAID also evaluates the prediction of binding sites within IDRs, commonly referenced to as MoRFs, SLIMs or LIPs^24,33^, leveraging DisProt annotations of binding regions (see Supplementary Figure 51 for dataset composition and overlap to other databases). In DisProt, binding annotations retrieved from literature are fraught with more ambiguity than disorder ones. In addition, the experimental evidence for the exact position of a binding region is often not accurate, as binding is annotated as a feature of an IDR. Our reference includes all entries in the DisProt dataset even if they were not annotated with binding regions. This translates to a dataset where the majority of targets (414 out of 646) have no positives. In this challenge, we kept the *PDB Observed* and *Gene3D* baselines even if they are not designed to detect binding regions. Target binding regions in DisProt are found within IDRs. Therefore, the baselines are expected to attain high recall and low precision. All models perform poorly as do the naive baselines (**Fig.s 4B, 4D**). At *F_Max_*, their recall is higher than their precision as for the baselines (**Fig. 4C**). However, the top 5 methods, ANCHOR-2^21^, DisoRDPbind^34^, MoRFchibi (light and web)^35^, and OPAL^36^, perform better than the baselines (**Fig. 4B**), which trade-off much more precision due to an abundant over-prediction. The execution time of the top five methods have very different scales and are inversely proportional to their performance, with the best methods requiring less CPU time. Performance of predictors on mammalian and prokaryotic proteins for the *DisProt-Binding* dataset are only marginal (see Supplementary Figures 62-71).

**Figure 4.**
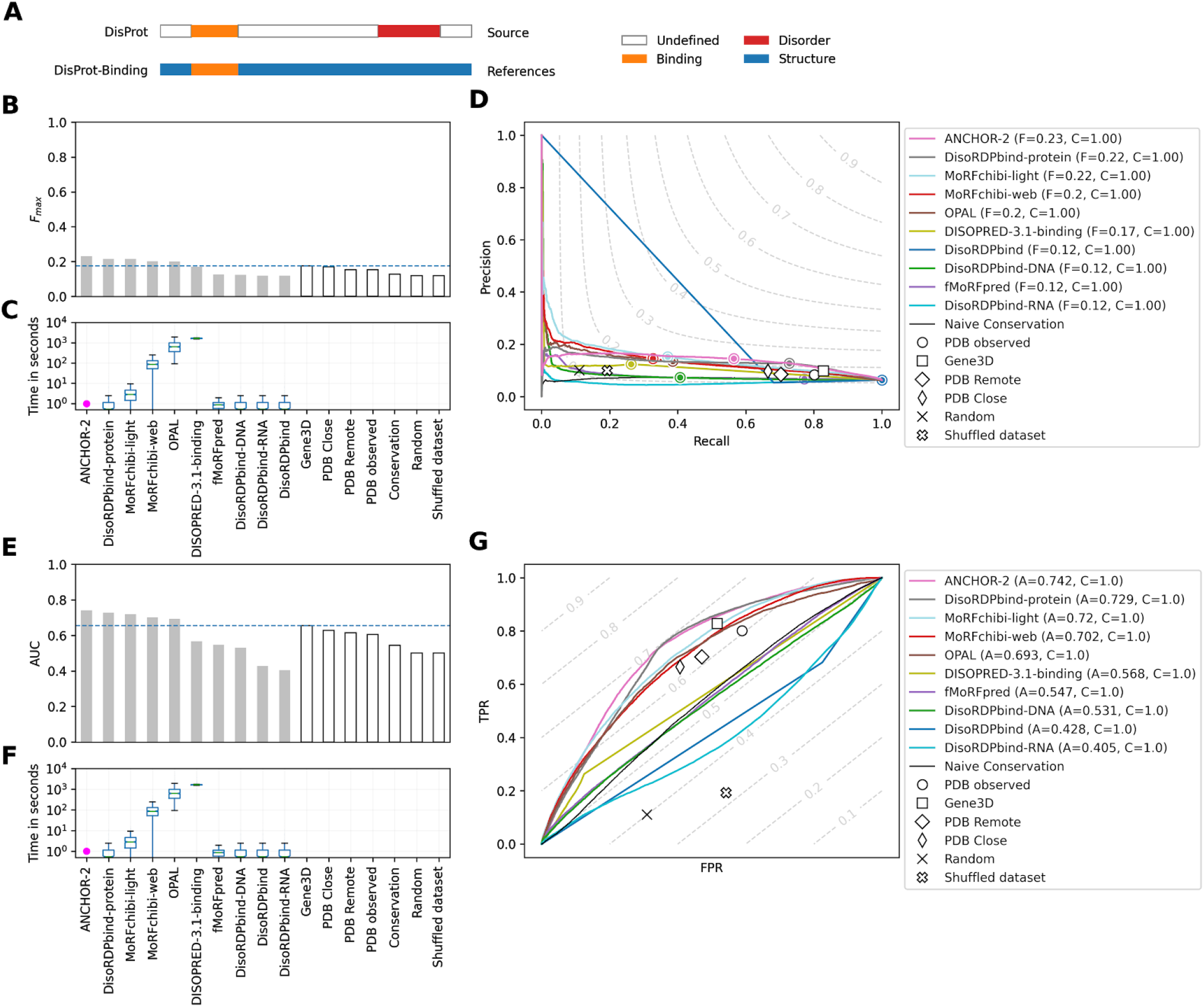
Prediction success and CPU times for the ten top-ranking binding predictors in the *DisProt-Binding* dataset. Reference used (*DisProt-Binding*) in the analysis and how it is obtained (panel **A**). Performance of predictors expressed as maximum F1-Score across all thresholds (F_max_) (panel **B**) and AUC (panel **E**) for the top ten best ranking methods (light gray) and baselines (white) and the distribution of execution time per-target (panels **C, F**) using *DisProt-Binding* dataset. The horizontal line in panels B, E indicates the F_max_ and AUC of the best baseline, respectively. Precision-Recall (panel **D**) and ROC curves (panel **G**) of ten top-ranking methods and baselines using *DisProt-Binding* dataset, with level curves of the F1-Score and Balanced accuracy, respectively. Magenta dots on panels C, F indicate that the whole distribution of execution-times is lower than 1 second

### Software implementation

We also evaluated for CAID technical aspects related to software implementation, i.e. speed and usability, which have a direct impact on the application of large-scale analyses. Speed in particular is highly variable, with methods of comparable performance varying by up to four orders of magnitude in execution time (see Supplementary Figure 4). In general, all methods incorporate a mix of different scripts and programming languages. Some software configuration scripts contain errors. In many cases data paths and file names are hardcoded inside the program, e.g. the path to the sequence database or the output file. Only a few programs allow specification of a temporary folder, which is important for parallel execution. It is possible to provide precalculated sequence searches only for a few methods. Several methods are implemented relying on dependencies, sometimes on specific software versions or CPUs with a modern instruction set. Some programs are particularly eager for RAM, crashing with longer input sequences, or do not have a timeout control and execute forever. Output formats differ, with some not adequately documented. Only a few software support multi-threading and only one was submitted as a Docker container. In summary, the software implementation for disorder predictors has considerable room for improvement in order to allow their use in practice.

## Discussion

The problem of predicting protein ID is challenging for several reasons. The first is in the definition of ID, which is a term that indicates that the sequence of a protein does not encode a stable structural state that is ordered. Defining ID as a property that a protein does not have (i.e. order) implies that many conformational states fit the definition, covering a continuum between fully disordered states and folded states with long dynamic regions^37,38^. The second problem, which follows from the first, is that we do not yet have a consensus reference experimental method, or set of experimental methods, to characterize ID (as we have X-ray crystallography to define ordered structures), so we lack a clear operational definition of ID. The third problem is that the disordered state is dependent on events or conditions at certain points in time along the life of a protein. Some proteins remain unfolded until they bind a partner^39^, others are disordered as long as they are in a specific cellular compartment and fold upon translocation^40^, and some enzymes undergo order-to-disorder-to-order transition as part of their catalytic cycle^41^.

Given these challenges, the CAID project represents a community effort to develop and implement evaluation strategies to assess: (1) clear definitions of ID and (2) the performance of methods to predict ID. Concerning (1), in its first round, CAID leverages the community-driven DisProt database^23^, a repository of manually curated experimental evidence of disorder, to blind test disorder predictors. In DisProt, curators store the coordinates of IDRs when there is experimental evidence in peer-reviewed articles of highly mobile residue stretches longer than 10 residues. We anticipate that future rounds may include reference data coming from ever-improving consensus operational definitions, as for example those coming from NMR measurements, which are particularly powerful in characterising experimentally protein disorder. For example, one could define disordered regions as those that exhibit high conformational variability under physiological conditions using multiple orthogonal measures. Concerning (2), this kind of challenge was tried from 5th to 10th editions of CASP but was abandoned due to the lack of good reference data. One of our long-term goals with CAID is to help guide the choice of candidate IDPs for experimentalists to test in their experiments.

One of the main properties of IDPs is their ability to form many low-affinity and high-specificity interactions^42^. It is, however, challenging to predict the interacting residues of an IDPs from its sequence. Presently, multiple high-throughput experiments for the detection of interactions capable of resolving the interacting regions exist^43^. However, binding sites obtained from high-throughput experiments (e.g. CoIP, Y2H) and reported in literature often lack this grade of resolution. Furthermore, while some attempts have been made to mitigate this problem^44^, a high rate of false positives plagues all experimental methods to identify binding: proteins interacting in experimental conditions do not necessarily interact in the cell under physiological physico-chemical conditions or simply due to spatio-temporal segregation^45^. DisProt annotates binding partners and interaction regions of IDPs, which we use in CAID to attempt the first assessment of binding predictors.

One of the major challenges we encountered in CAID is the identification of residues that are not disordered or do not bind, in other words, the definition of negatives. Knowledge about negative results is a long-standing problem in biology^46^ and is especially relevant for our assessment. If the annotation of IDRs in a protein is not complete, how do we know which regions of the protein are structured? This is even more relevant for binding regions, since at the moment we are far from mapping all binding partners of a protein with residue-resolution under different cellular settings. To overcome this problem intrinsic to how we detect and store data, we tested the performance of ID predictors in two scenarios. In the first scenario, we assumed that all annotations were complete, thus considering all the residues outside of annotated regions as structured. In the second scenario, we used resolved residues from PDBs to annotate structure and filtered out all residues that were neither covered by disorder nor structure annotation. Binding site predictors were tested on a dataset where all residues outside of binding regions are considered not-binding.

Despite these challenges, CAID revealed progress in the detection of ID from sequence and highlighted that there is still scope for improving both disorder- and binding-site predictors. One of the primary goals of CAID was to determine whether automated algorithms perform better than naive assumptions on the structural state based on indirect sources, such as sequence conservation or three-dimensional structure. As far as ID is concerned, the performance of predictors in comparison to naive baselines largely depends on the assumption made on non-disordered residues. On the *DisProt-PDB* dataset, where disorder is inferred from DisProt annotation and order inferred from the presence of a PDB structure and all other residues are filtered out, naive baselines outperform predictors. However, when only DisProt annotations are considered (*DisProt* dataset) tables are turned, and predictors, while obtaining lower overall scores, outperform naive baselines (**Fig.s 2 & 3**). When “uncertain” residues are kept in the analysis (*DisProt* dataset), the number of false positives increases and, as a result, precision plunges, lowering the F1-score. This means that predictors detect ID in the “uncertain” residues, indicating that DisProt annotation is incomplete, predictors over-predict or both. Naive baselines are outperformed by predictors since they predict all “uncertain” residues as disordered, which are all counted as false positives. This suggests that predictors have reached a state of maturity and can be trusted with relative confidence when no experimental evidence is available. It also confirms that when experimental evidence is present, it is more reliable than predictions.

An interesting particular case of disorder prediction is how predictors behave with DisProt targets whose annotations cover all (or almost all) residues, i.e. fully disordered proteins (Table 1). This case is compelling because usually predictors are not trained on these examples. Predictors vastly outperform naive baselines in these cases due to their large over-prediction. The count of false positives puts baselines at a disadvantage, compensating for their low count of false negatives. PDB Observed classifies a protein as fully disordered whenever no structure is available for that protein. However, the absence of a protein from PDB may be simply due to the lack of studies on that protein. Gene3D performs better since Gene3D models generalize from existing structures but still tends to over-predict disorder (or under-predict order). At the opposite side of the spectrum, methods that are too conservative in their disorder classification perform worse than expected on fully disordered proteins, for example, MobiDB-lite, which is conservative by design. Results on the *DisProt* dataset suggest several methods are consistently among the top performers, although the exact ranking is subject to some variation. fIDPnn and SPOT-Disorder2 perform consistently well, with RawMSA and AUCpreD following closely. The execution times for these four methods vary by up to three orders of magnitude, suggesting that there is room for optimizing the software. Of note, both fIDPnn and RawMSA were unpublished at the time of the CAID experiment.

While top-performing methods are able to achieve a certain balance between under- and over-prediction, it is interesting to notice how they are not able to identify all fully disordered targets: not even methods that trade-off specificity to increase their detection of relevant cases are able to attain full sensitivity. This confirms that predictors are not trained on this particular class of proteins and suggests that they have room for improvement in this direction.

The challenge of binding site identification is the first attempt at assessing binding predictors. As discussed above, this is intrinsically difficult due to the complex nature of this phenomenon and how it is detected and stored. While we are aware of these difficulties, we still think that an assessment is useful for researchers who either use or develop binding predictors. Furthermore, while it is arguable that this evaluation has limitations, its publication helps highlight such constraints and thus exposing this problem to the rest of the scientific community. We compared predictors to the same baselines used for the disorder challenge but, while their design remains unchanged, their underlying naive assumption changes slightly. The PDB observed baseline assumes that whatever is not covered by a structural annotation in PDB, is not only disordered but also involved in one or more interactions. When considering all targets in the CAID dataset, including those which are not annotated as binders, predictors slightly outperform the baselines, but have limited performance overall. Figure 4 shows a disagreement with the *DisProt-Binding* reference in both the positive and the negative classification, highlighting the potential for improvement of binding predictors. We have to consider that the dataset used is strongly unbalanced. Although a prominent function of IDPs is to mediate protein-protein interactions, most of the targets (414 of 646) do not contain an identified binding region and targets that do include binding regions often have them spanning the whole disordered region in which they are found. This strong bias is due to how DisProt was annotated in the past, with the label “binding” being associated with an entire IDR. In the latest version of DisProt this annotation style has been dropped in favor of a more detailed one, resulting in future editions of CAID being less biased towards long binding regions. The improved definition of the boundaries of disordered binding regions could favour methods that were trained specifically to recognize shorter binding regions. In all, this indicates that there is large growth potential in both the models and the reference building for this challenge.

In conclusion, the CAID experiment has provided the first fully blind assessment of ID predictors in almost a decade and the first-ever of ID binding regions. The results are encouraging, showing that the methods are mature enough to be useful but significant room for improvement remains. As the quality of ID data improves, we expect predictors to become more accurate and reliable.

## Author contributions

MN and DP collected the predictions, produced the data, carried out the assessment and wrote the initial manuscript. SCET designed the experiment, guided the overall project and edited the manuscript. The CAID Predictors contributed prediction methods for assessment. All authors contributed to the discussions and writing of the manuscript.

## Acknowledgements

The development of the predictors was supported in part by National Science Foundation (grant 1617369), Natural Sciences and Engineering Research Council of Canada (grant 298328), Tianjin Municipal Science and Technology Commission (grant 13ZCZDGX01099), National Natural Science Foundation of China (grants 31970649 and 11701296), Natural Science Foundation of Tianjin (grant 18JCYBJC24900), Japan Agency for Medical Research and Development (grant 16cm0106320h0001), Australian Research Council (ARC) [DP180102060], Research Foundation Flanders (FWO) - project nr. G.0328.16N, Agence Nationale de la Recherche (ANR-14-CE10-0021, ANR-17-CE12-0016). The state task “Bioinformatics and proteomics studies of proteins and their complexes” N<u>o</u> 0115-2019-004 for OG and ML. ZD acknowledges funding from ELTE Thematic Excellence Programme (ED-18-1-2019-0030) supported by the Hungarian Ministry for Innovation and Technology and the “Lendület” grant from the Hungarian Academy of Sciences (LP2014-18). PT acknowledges grants K124670 and K131702 from the Hungarian Scientific Research Fund (OTKA). This project has received funding from the European Union’s Horizon 2020 research and innovation programme under the Marie Skłodowska-Curie grant agreement No 778247, the Italian Ministry of University and Research PRIN 2017 grant 2017483NH8 and ELIXIR, the European infrastructure for biological data.

## CAID Predictors

Tamjidul Hoque; Computer Science, University of New Orleans, USA

Ian Walsh; Bioprocessing Technology Institute, Agency for Science, Technology and Research (A*STAR), Singapore

Sumaiya Iqbal; Broad Institute of MIT and Harvard, USA

Michele Vendruscolo, Pietro Sormanni; Department of Chemistry, University of Cambridge, UK

Chen Wang; Columbia University, USA

Daniele Raimondi; ESAT-STADIUS, KU Leuven, Belgium

Ronesh Sharma; Fiji National University, Suva, Fiji

Yaoqi Zhou, Thomas Litfin; Institute for Glycomics and School of Information and Communication Technology, Griffith University, Australia

Oxana V. Galzitskaya, Michail Yu. Lobanov; Institute of Protein Research; Russian Academy of Sciences, Russia

Wim Vranken; Interuniversity Institute of Bioinformatics in Brussels, Vrije Universiteit Brussel, Belgium

Björn Wallner, Claudio Mirabello; Linköping University

Nawar Malhis; Michael Smith Laboratories, University of British Columbia, Canada

Zsuzsanna Dosztányi, Gábor Erdős, Bálint Mészáros; Department of Biochemistry, Eötvös Loránd University, Hungary

Jianzhao Gao, Kui Wang, Gang Hu, Zhonghua Wu; Nankai University, China

Alok Sharma; RIKEN Center for Integrative Medical Sciences, Japan; Griffith University, Australia

Jack Hanson, Kuldip Paliwal; School of Engineering and Built Environment, Griffith University, Australia

Isabelle Callebaut, Tristan Bitard-Feildel; Sorbonne Université, Muséum National d’Histoire Naturelle, UMR CNRS 7590, Institut de Minéralogie, de Physique des Matériaux et de Cosmochimie, Paris, France

Gabriele Orlando; VIB-KU Leuven, 3000 Leuven, Belgium

Zhenling Peng; Tianjin University, China

Jinbo Xu, Sheng Wang; Toyota Technological Institute at Chicago, USA

David T. Jones, Domenico Cozzetto; University College London, UK

Fanchi Meng, Jing Yan; University of Alberta, Canada

Joerg Gsponer; University of British Columbia

Jianlin Cheng, Tianqi Wu; University of Missouri, Columbia

Lukasz Kurgan; Virginia Commonwealth University, USA

## DisProt Curators

Vasilis J Promponas, Stella Tamana; Department of Biological Sciences, University of Cyprus, Cyprus

Cristina Marino-Buslje, Elizabeth Martínez-Pérez; Fundación Instituto Leloir, Argentina

Anastasia Chasapi, Christos Ouzounis; Chemical Process & Energy Resources Institute, Centre for Research & Technology Hellas, Greece

A. Keith Dunker; Center for Computational Biology and Bioinformatics, Indiana University School of Medicine, USA

Andrey V Kajava, Jeremy Y Leclercq; Centre de Recherche en Biologie cellulaire de Montpellier (CRBM), UMR 5237 CNRS, Université Montpellier, France

Burcu Aykac-Fas, Matteo Lambrughi, Emiliano Maiani, Elena Papaleo; Danish Cancer Society Research Center, Denmark

Lucia Beatriz Chemes, Lucía Álvarez, Nicolás S González Foutel; Consejo Nacional de Investigaciones Científicas y Técnicas. Instituto de Investigaciones Biotecnológicas IIBIO, Universidad Nacional de San Martín, Argentina

Valentin Iglesias, Jordi Pujols, Salvador Ventura; Departament de Bioquimica i Biologia Molecular and Institut de Biotecnologia i Biomedicina, Universitat Autònoma de Barcelona, Spain

Nicolás Palopoli, Guillermo Ignacio Benítez, Gustavo Parisi; Departamento de Ciencia y Tecnología, Universidad Nacional de Quilmes - CONICET, Argentina

Claudio Bassot, Arne Elofsson, Sudha Govindarajan, John Lamb, Marco Salvatore; Department of Biochemistry and Biophysics and Science for Life Laboratory, Stockholm University, Sweden

András Hatos, Alexander Miguel Monzon, Martina Bevilacqua, Ivan Mičetić, Giovanni Minervini, Lisanna Paladin, Federica Quaglia; Department of Biomedical Sciences, University of Padova, Italy

Emanuela Leonardi; Department of Woman and Child Health, University of Padova, Italy

Norman Davey; Division of Cancer Biology, The Institute of Cancer Research, UK

Tamas Horvath, Orsolya Panna Kovacs, Nikoletta Murvai, Rita Pancsa, Eva Schad, Beata Szabo, Agnes Tantos; Institute of Enzymology, Research Centre for Natural Sciences, Hungarian Academy of Sciences, Hungary

Sandra Macedo-Ribeiro, Jose A Manso, Pedro José Barbosa Pereira; Instituto de Biologia Molecular e Celular (IBMC) and Instituto de Investigação e Inovação em Saúde (i3S), Universidade do Porto, Portugal

Radoslav Davidović, Nevena Veljkovic; Institute of Nuclear Sciences Vinca, University of Belgrade, Serbia

Borbála Hajdu-Soltész, Mátyás Pajkos, Tamás Szaniszló; Department of Biochemistry, Eötvös Loránd University, Hungary

Mainak Guharoy, Tamas Lazar, Mauricio Macossay-Castillo, Peter Tompa; Structural Biology Brussels, Vrije Universiteit Brussel (VUB), Belgium

## Methods

All software programs were executed using a homogeneous cluster of nodes running Ubuntu 16.04 on Intel 8 core processors with 16 GB of RAM and a mechanical harddisk. In the text we refer to proteins as targets, to disordered residues as positive labels and structured/ordered residues as negative labels.

### Reference sets

In CAID different reference sets were built, differing in the subset of DisProt used to define positive labels and in the definition of negatives labels.

For the disorder challenge, we generated two reference sets called *DisProt* and *DisProt-PDB*. Both references are composed of a set of 646 targets, annotated between June 2018 and November 2018 (DisProt release 2018_11). Positive labels in both reference sets are those residues annotated as disordered in the DisProt database. In the *DisProt* reference set, all labels that are not positive are assigned as negatives. In the *DisProt-PDB* set, PDB structures mapping on the protein sequence define negative labels. All residues that are not covered by either DisProt annotation or PDB structures, are masked and excluded from the analysis. It should be noted that a fraction of resolved structures in the PDB have been annotated as disordered^47,48^. While in this edition we decided to consider any resolved residue from crystallography, NMR or electron microscopy experiments (excluding those overlapping with DisProt annotation) as structured, we plan to apply a filtering on the next editions of CAID. This problem will become less and less relevant as DisProt annotations will become more complete, since disorder always overwrites structure.

For the binding challenge we generated a reference set that we called *DisProt-Binding*. Positive labels are those residues annotated as binding in the DisProt database, whereas all labels that are not positive are assigned as negatives. Notice that 232 targets have at least one annotation of binding in the DisProt database. *DisProt-Binding* is composed of all 646 targets considered in the analysis, hence the majority of targets (i.e. 646 − 232 = 414) do not contain positive labels.

### Predictions

Most predictors output a series of score and state pairs per residue of the input sequence. Scores are floating point numbers, while states are binary labels predicting if a residue is in a disordered or structured state. If scores are missing, states will be used as scores. If states are missing, they are generated by applying a threshold to scores. When a threshold is not available, it defaults to 0.5. Default threshold is always inferred from states. This ensures correct default threshold estimates for any distribution of scores. Prediction scores are rounded to the third decimal figure. This sets the number of possible thresholds to 1000. Bootstrapping samples the whole dataset with replacements 1000 times. Re-sampling is done at the label (residue) level. Confidence intervals are calculated on Student’s T distribution at alpha set to 0.05.

### Baselines

A number of baseline predictors have been built in order to be compared with actual predictors. Two are based on randomizing the dataset (*Shuffled dataset, Random*) and one is based on an estimate of residue conservation through evolution (*Conservation*). The last four consider the opposite of structure as disorder (*PDB Observed, PDB Close, PDB Remote* and *Gene3D*).

*Shuffled dataset* is a reshuffling of the *DisProt* dataset, i.e. random permutation of labels across the entire dataset. This preserves the proportion of positive labels across the dataset but not necessarily for each single target. The *Random* baseline is a random classifier in which the prediction score of each label is assigned randomly. It is built by randomly drawing floating point numbers out of a uniform distribution [0,…,1] and applying a threshold of 0.5.

The *Conservation* baseline uses the naive consideration that IDPs on average are less conserved than globular proteins. It is calculated from the distance between the residue frequencies of homologous sequences for each target against the residue frequencies of the BLOSUM62 substitution matrix. Homologous sequences are retrieved by running 3 iterations of PSI-BLAST^49^ against UniRef90. The distance is calculated from the Jensen-Shannon divergence^50^ of the two frequencies. This returns values in the [0,…,1] interval where any position with a score above 0.4 is considered positive (i.e. disordered).

Several naive baselines are based on the assumption that whatever is not annotated as structure in the PDB is disordered. *PDB Observed* has the structure annotation defined by PDB structures as mapped on UniProt sequences by Mobi 2.0^51^ (October 2019). Whenever we are unable to map perfectly the PDB sequence on the UniProt sequence, unmapped residues are considered not observed and excluded from the analysis. This applies to His-tags, mutated sequences and missing residues (in both X-ray and NMR structures), PDB Close and PDB Remote have the structure annotation defined by observed residues in PDBs with similar sequence. The similarity is calculated as the identity percentage given by a 3 iteration PSI-BLAST^49^ of DisProt targets against PDB seqres. PDB Close considers PDB structures with at least 30% of sequence identity (i.e. close homologs), while PDB remote considers only PDB structures with sequence identity between 20% and 30% (i.e. remote homologs). *Gene3D* has structure annotations defined by Gene3D^52^ (version 4.2.0) predictions, calculated with InterProScan^28^ (version 5.38-76.0).

### Target and Dataset metrics

Metrics were calculated following two strategies: dataset and target. In the dataset strategy, all targets (proteins) reference classifications and prediction classifications are concatenated in two single arrays. Confusion matrix and subsequent evaluation metrics are calculated once comparing these arrays. In the target strategy confusion matrix and subsequent evaluation metrics are calculated for each target (protein) and the mean value of the evaluation metrics is taken. The former strategy is equivalent to summing together the confusion matrices for each target and computing evaluation metrics on the resulting confusion matrix, while the latter strategy is equivalent to calculating the evaluation metrics on the average of the confusion matrices of the targets.

### Notes on evaluation metrics calculation

Throughout the manuscript, F_max_ and AUC are the main assessment criteria used. F_max_ is the maximum point in the precision/recall curve. AUC is the area under the ROC curve. Additional metrics are used for comparison and they all follow standard definitions as described in the Supplementary Table 3. Fbeta (0.5, 1, 2) and MCC are set to 0 if the denominator is 0. Since the MCC denominator is a multiplication of the number of positive classifications, negative classifications, positive labels in the reference and negative labels in the reference, if any of these classes amounts to 0, we set MCC to 0. This means that for fully disordered proteins and for proteins predicted to be fully disordered or fully ordered, MCC is 0.

This situation is very likely in target-strategy with *DisProt-PDB* dataset and it explains why the MCC for target-strategy is much lower than that for dataset-strategy (see Supplementary Figure 34). This effect can also be seen in the heatmap of targets MCC where a large number of targets have MCC = 0.

### Assessors policy

Prediction methods published by the assessors did not take part in the challenges. Their methods may be included for reference only.

### Data Availability

Raw DisProt annotations can be downloaded in the download section of the DisProt website. Dataset downloaded from DisProt were:

- disprot-2018-11-disorder.fasta obtained from: https://disprot.org/api/search?release=2018_11&show_ambiguous=false&show_obsolete=false&format=fasta&namespace=structural_state&get_consensus=true
- disprot-2016-10-disorder.fasta obtained from: https://disprot.org/api/search?release=2016_10&show_ambiguous=false&show_obsolete=false&format=fasta&namespace=structural_state&get_consensus=true
- disprot-2018-11-interaction.fasta obtained from: https://disprot.org/api/search?release=2018_11&show_ambiguous=false&show_obsolete=false&format=fasta&namespace=interaction_partner&get_consensus=true
- disprot-2018-11.json obtained from: https://disprot.org/api/search?release=current&show_ambiguous=false&show_obsolete=false&format=json

Datasets were processed to produce references, which are available at: http://idpcentral.org/caid/data/1/ The process and code to produce references is available in the CAID repository with scripts available in the CAID repository: https://github.com/BioComputingUP/CAID

Predictions formatted in the CAID format are available at: http://idpcentral.org/caid/data/1/.

All data used in the analysis is also available in the Code Ocean capsule available at: https://doi.org/10.24433/CO.3610625.v1.

### Software Description and Availability

Results of the CAID assessment can be fully reproduced downloading the code and following the instructions in the CAID repository: https://github.com/BioComputingUP/CAID

The CAID software is a python3 package that produces all outputs necessary for a CAID edition, including baselines, references, metrics and plots, starting from predictions, references and sequence annotations from complementary sources. (see Data Availability section to know how to obtain this data). The CAID repository depends on published python3 libraries and on the vectorized_cls_metrics library, available at https://github.com/marnec/vectorized_cls_metrics.

The code is also available at and its reproducibility ensured by the Code Ocean capsule available at https://doi.org/10.24433/CO.3610625.v1.

