## supplementary material for "Critical Assessment of Protein Intrinsic Disorder Prediction"

### Supplementary Information

This document is structured as follows:


| Section | Description | Page |
|---|---|---|
| **Tables** | | 2 |
| Prediction methods | List of disorder and binding predictors | 2 |
| Evaluation metrics | List of evaluation metrics used throughout the analysis | 9 |
| DisProt dataset | | 10 |
| Evaluation Results | Performance of disorder predictors on the *DisProt* dataset | 10 |
| Evaluation Results (Full ID) | Performance of disorder predictors on the soft challenge of telling apart fully disordered proteins | 12 |
| DisProt-PDB dataset | | 15 |
| Evaluation Results | Performance of disorder predictors on the *DisProt-PDB* dataset | 15 |
| Disprot-Binding dataset | | 17 |
| Evaluation Results | Performance of binding predictors on the *DisProt-Binding* dataset | 17 |
| **Figures** | | 19 |
| Dataset | | 19 |
| Annotations | Statistics on the disorder annotations in the references | 19 |
| Redundancy | Statistics on the sequence similarity of reference targets versus other targets in DisProt | 20 |
| Predictor CPU time | Statistics on the execution time of predictors | 20 |
| Species representation | Origin species of reference targets | 22 |
| Fully ID | Disorder detection methods annotated in targets of the fully disordered proteins subset | 23 |
| Disorder | | 23 |
| DisProt dataset | Performance of disorder predictors on the *DisProt* dataset | 23 |
| Mammals | Performance of disorder predictors on the subset of mammalian proteins in the *DisProt* dataset | 32 |
| Prokaryotes | Performance of disorder predictors on the subset of | 37 |

| | | |
|---|---|---|
| | prokaryotic proteins in the *DisProt* dataset | |
| DisProt-PDB dataset | Performance of disorder predictors on the *DisProt-PDB* dataset | 41 |
| Mammals | Performance of disorder predictors on the subset of mammalian proteins in the *DisProt-PDB* dataset | 51 |
| Prokaryotes | Performance of disorder predictors on the subset of prokaryotic proteins in the *DisProt-PDB* dataset | 56 |
| Binding | | 61 |
| DisProt-Binding dataset | Performance of binding predictors on the *DisProt-Binding* dataset | 62 |
| Mammals | Performance of binding order predictors on the subset of mammalian proteins in the *DisProt-Binding* dataset | 68 |
| Prokaryotes | Performance of binding predictors on the subset of prokaryotic proteins in the *DisProt-Binding* dataset | 73 |


### Tables

#### Prediction methods

| Method | Principal Investigator | Keywords | Publication (PubMed ID) |
|---|---|---|---|
| AUCpreD | Jinbo Xu | sequence labeling, deep conditional neural fields, conditional random fields, deep convolutional neural networks, AUC-maximization | 27587688 |
| AUCpreD_np | Jinbo Xu | sequence labeling, deep conditional neural fields, conditional random fields, deep convolutional neural networks, AUC-maximization | 27587688 |
| DFLpred | Lukasz Kurgan | disordered linkers, logistic regression, sliding window | 27307636 |
| DisEMBL-465 | Rob B. Russell | Coils, missing coordinates, target selection | 14604535 |
| DisEMBL-HL | Rob B. Russell | Coils, missing coordinates, loops | 14604535 |
| DisoMine | Wim Vranken | Protein disorder prediction, recurrent neural network, biophysical features | n/a |
| DISOPRED-3.1 | David T Jones | Meta-predictor, neural network, long IDRs | 25391399 |
| DisPredict2 | Tamjidul Hoque | PSEE, contact energy, burial | 27588752 |
| DynaMine | Wim Vranken | Protein backbone dynamics prediction, linear regression, biophysical information | 24225580 |
| ESpritz-D | Silvio Tosatto | neural network, fast, DisProt | 22190692 |
| ESpritz-N | Silvio Tosatto | neural network, fast, NMR | 22190692 |
| ESpritz-X | Silvio Tosatto | neural network, fast, xray | 22190692 |

| | | | |
|---|---|---|---|
| fIDPlr | Lukasz Kurgan | meta-predictor, disorder function, binding, sliding window | n/a |
| fIPDnn | Lukasz Kurgan | meta-predictor, disorder function, binding | n/a |
| FoldUnfold | Oxana V. Galzitskaya | intrinsically flexibility, size of window, threshold | 17021161 |
| GlobPlot | Toby J. Gibson | Globularity, propensity, secondary structure, missing electron densities | 12824398 |
| IsUnstruct | Oxana V. Galzitskaya | residual potentials, boundary energy, disordered patterns, two-state model | 21572175 |
| IUpred-long | Zsuzsanna Dosztányi | energy estimation, statistical potential, fast | 15955779 |
| IUpred-short | Zsuzsanna Dosztányi | energy estimation, statistical potential, fast | 15955779 |
| IUpred2A-long | Zsuzsanna Dosztányi | energy estimation, statistical potential, fast | 29860432 |
| IUpred2A-short | Zsuzsanna Dosztányi | energy estimation, statistical potential, fast | 29860432 |
| JRONN | Robert Esnouf | Regional order neural network | 15947016 |
| MobiDB-lite | Silvio Tosatto | Meta-predictor, fast, InterPro | 28453683 |
| Predisorder | Jianlin Cheng | recurrent neural networks, disorder prediction | 20025768 |
| pyHCA | Isabelle Callebaut | Hydrophobic cluster, residue physico chemical properties, sequence topology, SVC | n/a |
| rawMSA | Björn Wallner | deep network, bidirectional neural network, embeddings, multiple sequence alignment | 31415569 |
| s2D-1 | Michele Vendruscolo | Secondary structure, intrinsic disorder, NMR spectroscopy, sequence-based predictions | 25534081 |
| s2D-2 | Michele Vendruscolo | Secondary structure, intrinsic disorder, NMR spectroscopy, sequence-based predictions | n/a |
| SPOT-Disorder1 | Yaoqi Zhou | bidirectional recurrent neural network, LSTM | 28011771 |
| SPOT-Disorder-Single | Yaoqi Zhou | single-sequence, deep neural network | 28011771 |
| SPOT-Disorder2 | Yaoqi Zhou | IncReSeNet | 32173600 |
| VSL2B | Zoran Obradovic | SVM, DisProt, xray | 16618368 |

**Supplementary Table 1.** Participating methods for **disorder region** prediction grouped according to the Principal Investigator.

| **Method** | **Principal Investigator** | **Keywords** | **Publication (PubMed ID)** |
|---|---|---|---|
| ANCHOR | Zsuzsanna Dosztányi | energy estimation, statistical potential, fast, disorder-to-order transition, energy gain | 19717576 |
| ANCHOR-2 | Zsuzsanna Dosztányi | energy estimation, statistical potential, fast, disorder-to-order transition, energy gain | 29860432 |
| DISOPRED-3.1 binding | David T. Jones | Meta-predictor, machine-learning, long IDRs, SVM | 25391399 |
| DisoRDPbind_all | Lukasz Kurgan | disorder function, nucleic acids, DNA-binding, RNA-binding, protein-binding | 26109352 |
| DisoRDPbind_dna | Lukasz Kurgan | disorder function, nucleic acids, DNA-binding, | 26109352 |

| | | | |
|---|---|---|---|
| | | RNA-binding, protein-binding | |
| DisoRDPbind_prot | Lukasz Kurgan | disorder function, nucleic acids, DNA-binding, RNA-binding, protein-binding | 26109352 |
| DisoRDPbind_rna | Lukasz Kurgan | disorder function, nucleic acids, DNA-binding, RNA-binding, protein-binding | 26109352 |
| fMoRFpred | Lukasz Kurgan | MoRF, induced folding, molecular recognition features, sliding window | 26651072 |
| MorfChibi-light | Joerg Gsponer | Molecular recognition features (MoRFs), Support Vector Machines, Hierarchical learning | 27174932 |
| MorfChibi-web | Joerg Gsponer | Molecular recognition features (MoRFs), Support Vector Machines, Hierarchical learning, Conservation | 27174932 |
| OPAL | Alok Sharma | Molecular Recognition Features, Intrinsically Disordered Proteins, Support Vector Machine, BigramMoRF and StructMoRF | 29360926 |

**Supplementary Table 2.** Participating methods for **disorder binding site** prediction grouped according to the Principal Investigator.

| **Method (Principal Investigator)** | **Description** |
|---|---|
| **ANCHOR (Zsuzsanna Dosztányi)** | ANCHOR is a fast method that predicts disordered regions that can undergo a disorder-to-order transition upon binding to globular protein partners. To predict such regions ANCHOR aims to capture the general disorder tendency of these regions and their ability to gain energy by interacting with a more structured environment using the estimated energy approach. Most of the parameters are based on known protein structures and only five parameters were optimized on known disordered binding regions, lending the methods its robustness. |
| **ANCHOR2 (Zsuzsanna Dosztányi)** | ANCHOR2 retains the original idea behind ANCHOR and employs a simple biophysics-based model to predict disordered binding regions in a similar way. This approach relies on a larger collection of known disordered binding regions to improve on the modeling of the structured environment of these regions. |
| **AUCpreD (Jinbo Xu)** | The AUCpreD method developed by Xu group formulates IDR prediction as a sequence labeling problem and employs a machine learning technique called Deep Convolutional Neural Fields (DeepCNF) to solve it. DeepCNF is an integration of deep convolutional neural networks (DCNN) and conditional random fields (CRF). DeepCNF can model not only complex sequence-structure relationships in a hierarchical manner, but also the correlation of order and disorder labels among adjacent residues. AUCpredD can predict IDRs from not only sequence profiles (i.e., evolutionary information), but also primary sequence. Predicting from a primary sequence is not as accurate as predicting from a sequence profile, but much faster. The distribution of ordered and disordered residues is highly imbalanced (about 15:1), which makes it challenging to accurately predict IDRs using traditional machine learning approaches. To deal with this, AUCpredD is trained by maximizing area under the ROC curve (AUC), which is an unbiased measure for class-imbalanced data. Four publicly available datasets are used to train, validate and evaluate AUCpreD. In particular, the UniProt90 dataset released before May 01, 2010 is |

| | |
|---|---|
| | used to train and validate the model parameters. The CASP9, CASP10 and CAMEO test proteins are used to evaluate prediction accuracy. Sequence identity 25% is employed to remove redundancy between training and test data. In total, there are 13,800 training and validation proteins and a 10-fold cross-validation is performed to train 10 different models, which are then combined to form the final prediction model. |
| **AUCpreD_no-profile (Jinbo Xu)** | The AUCpreD method developed by Xu group formulates IDR prediction as a sequence labeling problem and employs a machine learning technique called Deep Convolutional Neural Fields (DeepCNF) to solve it. DeepCNF is an integration of deep convolutional neural networks (DCNN) and conditional random fields (CRF). DeepCNF can model not only complex sequence-structure relationships in a hierarchical manner, but also the correlation of order and disorder labels among adjacent residues. AUCpredD can predict IDRs from not only sequence profiles (i.e., evolutionary information), but also primary sequence. Predicting from a primary sequence is not as accurate as predicting from a sequence profile, but much faster. The distribution of ordered and disordered residues is highly imbalanced (about 15:1), which makes it challenging to accurately predict IDRs using traditional machine learning approaches. To deal with this, AUCpredD is trained by maximizing area under the ROC curve (AUC), which is an unbiased measure for class-imbalanced data. Four publicly available datasets are used to train, validate and evaluate AUCpreD. In particular, the UniProt90 dataset released before May 01, 2010 is used to train and validate the model parameters. The CASP9, CASP10 and CAMEO test proteins are used to evaluate prediction accuracy. Sequence identity 25% is employed to remove redundancy between training and test data. In total, there are 13,800 training and validation proteins and a 10-fold cross-validation is performed to train 10 different models, which are then combined to form the final prediction model. |
| **DFLpred (Lukasz Kurgan)** | Logistic regression based on custom-designed features extracted from putative disorder and propensity for helix and coil conformations. Fast and requires protein sequence as the only input, thus applicable to prediction at the whole genome scale. |
| **DisEMBL-465 (Rob B. Russel)** | DisEMBL is a computational tool for prediction of disordered/unstructured regions within a protein sequence. As no clear definition of disorder exists, we have developed parameters based on several alternative definitions, and introduced a new one based on the concept of ``hot loops'', i.e. coils with high temperature factors. DisEMBL is useful for target selection and the design of constructs as needed for many biochemical studies, particularly structural biology and structural genomics projects. |
| **DisEMBL-HL (Rob B. Russel)** | DisEMBL is a computational tool for prediction of disordered/unstructured regions within a protein sequence. As no clear definition of disorder exists, we have developed parameters based on several alternative definitions, and introduced a new one based on the concept of ``hot loops'', i.e. coils with high temperature factors. DisEMBL is useful for target selection and the design of constructs as needed for many biochemical studies, particularly structural biology and structural genomics projects. |
| **DisoMine (Wim Vranken)** | DisoMine predicts protein disorder with recurrent neural networks from simple predictions of protein dynamics, secondary structure and early folding. The tool is fast and requires only a single sequence, making it applicable for large-scale screening, including poorly studied and orphan proteins. |
| **DISOPRED-3.1 (David T. Jones)** | DisoPred3 first identifies disordered residues through a consensus of the output generated by DISOPRED2 and two additional machine-learning based modules trained on large IDRs, and then annotates them as protein binding through an additional SVM classifier |
| **DISOPRED-3.1_binding (David T. Jones)** | DISOPRED3.1 binding finds binding regions in disordered regions predicted by DISOPRED3. It’s based on an SVM classifier. Using a sliding window of size 15, we derived three independent SVM classifiers from the training data that are based on (i) single sequences alone; (ii) PSSM values obtained after three search iterations of PSI-BLAST against UniRef90; (iii) the same PSSM scores, followed by the length of input |

| | |
|---|---|
| | region. |
| **DisoRDPbind_all (Lukasz Kurgan)** | Logistic regression based on custom-designed features extracted from putative disorder, putative secondary structure, sequence complexity profile, and selected physicochemical properties of residues. Regression predictions are combined with results of the alignment into a dataset of functionally annotated disordered regions. Fast and requires protein sequence as the only input, thus applicable to prediction at the whole genome scale. |
| **DisoRDPbind_dna (Lukasz Kurgan)** | Logistic regression based on custom-designed features extracted from putative disorder, putative secondary structure, sequence complexity profile, and selected physicochemical properties of residues. Regression predictions are combined with results of the alignment into a dataset of functionally annotated disordered regions. Fast and requires protein sequence as the only input, thus applicable to prediction at the whole genome scale. |
| **DisoRDPbind_prot (Lukasz Kurgan)** | Logistic regression based on custom-designed features extracted from putative disorder, putative secondary structure, sequence complexity profile, and selected physicochemical properties of residues. Regression predictions are combined with results of the alignment into a dataset of functionally annotated disordered regions. Fast and requires protein sequence as the only input, thus applicable to prediction at the whole genome scale. |
| **DisoRDPbind_rna (Lukasz Kurgan)** | Logistic regression based on custom-designed features extracted from putative disorder, putative secondary structure, sequence complexity profile, and selected physicochemical properties of residues. Regression predictions are combined with results of the alignment into a dataset of functionally annotated disordered regions. Fast and requires protein sequence as the only input, thus applicable to prediction at the whole genome scale. |
| **DisPredict2 (Md Tamjidul Hoque)** | DisPredic2 uses position specific estimated energy, named PSEE, for each residue of a protein based on sequence information alone. The quantification of PSEE includes the interaction effect of the target residue within a neighborhood in terms of pairwise contact energies between different amino acid types. Neighborhood size is estimated in terms of the number of residues on either side of the target residue with which it can form favorable contacts. Furthermore, it utilizes the predicted relative exposure (or burial) of a residue to approximate the local three-dimensional conformational position and stability of the residue. |
| **DynaMine (Wim Vranken)** | DynaMine is a fast, high-quality predictor of protein backbone dynamics from single protein sequences. DynaMine is trained on information derived from experimental NMR chemical shift data for proteins in solution, and can identify disordered regions within proteins without depending on prior disorder knowledge or three-dimensional structural information. |
| **ESpritz-D (Silvio Tosatto)** | ESpritz combines a sophisticated BRNN architecture with enhanced definitions of disorder flavors. This version is based on DisProt2. The BRNN improves performance in general compared to previous iterations of this predictor (Spritz3 and CSpritz4) and especially on this training-set. |
| **ESpritz-N (Silvio Tosatto)** | ESpritz combines a sophisticated BRNN architecture with enhanced definitions of disorder flavors. This is based on NMR mobility calculated on NMR conformers from PDB. The BRNN improves performance in general compared to previous iterations of this predictor (Spritz3 and CSpritz4) and substantially on this training-set. |
| **ESpritz-X (Silvio Tosatto)** | ESpritz combines a sophisticated BRNN architecture with enhanced definitions of disorder flavors. This version is based on X-ray structures from PDB. The BRNN improves performance in general compared to previous iterations of this predictor (Spritz3 and CSpritz4) and slightly on this training-set. |
| **fIDPlr (Lukasz Kurgan)** | Logistic regression model trained with a comprehensive set of custom-designed features extracted from predicted disorder and disorder function predicted with DFLpred, DisoRDPbind and fMoRFpred and pre-processed using wrapper-based feature selection. Fast and requires protein sequence as the only input, thus applicable to prediction at the whole genome scale. |

| | |
|---|---|
| **flPDnn (Lukasz Kurgan)** | Dense and deep neural network trained with a comprehensive feature set extracted from predicted disorder and disorder function predicted with DFLpred, DisoRDPbind and fMoRFpred. Fast and requires protein sequence as the only input, thus applicable to prediction at the whole genome scale. |
| **fMoRFpred (Lukasz Kurgan)** | Support Vector Machine (SVM) based on custom-designed features extracted from putative intrinsic disorder, putative secondary structure, estimated B-factor and physicochemical characteristics including structural stability and unfolding energy. Fast and requires protein sequence as the only input, thus applicable to prediction at the whole genome scale. |
| **FoldUnfold (Oxana V. Galzitskaya)** | A new parameter, namely, the average packing density of the residues, was introduced to detect disordered regions in the protein sequence. We showed that regions with a low expected packing density will be responsible for the appearance of disordered regions. |
| **GlobPlot (Toby J Gibson)** | GlobPlot identifies regions of globularity and disorder within protein sequences based on a running sum of the propensity for amino acids to be in an ordered or disordered state. The GlobPlot package currently contains seven different propensity sets, as for example tendency to form secondary structure and missing electron densities in X-Ray experiments. |
| **IsUnstruct (Oxana V. Galzitskaya)** | The Ising model is used. The energy of transfer to the unfolded state is attributed to each residue, and the energy of the boundary between the folded and unfolded states is also taken into account. These energies were optimally selected based on PDB. |
| **IUPred-long (Zsuzsanna Dosztányi)** | IUPred-long predicts protein disorder based on an energy estimation approach utilizing statistical potentials. The parameters of the method are derived from a collection of known structures only and disordered regions are recognized based on their unfavorable estimated energies. IUPred is a fast and robust method that carries out predictions for single protein sequences without using evolutionary information. |
| **IUPred-short (Zsuzsanna Dosztányi)** | IUPred-short is based on the same approach as IUPred-long but choices of window sizes are slightly tailored towards predicting missing residues from PDB structures. |
| **IUPred2A-long (Zsuzsanna Dosztányi)** | IUPred2A-long is an implementation of IUPred-long in PYTHON with minor bug fixes. |
| **IUPred2A-short (Zsuzsanna Dosztányi)** | IUPred2A-short is an implementation of IUPred-short in PYTHON with minor bug fixes. |
| **JRONN (Robert Esnouf)** | Regional order neural network (RONN) software as an application of our recently developed 'bio-basis function neural network' pattern recognition algorithm to the detection of natively disordered regions in proteins. The decision about the likelihood of disorder is based on alignments to an ensemble of sequences of known folding state. |
| **MobiDB-lite (Silvio Tosatto)** | MobiDB-lite is a meta-predictor that combines the results of 8 highly orthogonal disorder predictors in a consensus. A post processing phase smooths a strict majority consensus. Finally, predicted regions shorter than 20 residues are filtered out. |
| **MorfChibi-light (Joerg Gsponer)** | MoRFchibi_light scores are computed using Hierarchical Learning [PMID: 26517836, 30952844]. Specifically, the scores of two modules (MoRFchibi and ESpritz) are combined hierarchically using Bayes rule to generate the MoRFchibi_web score. The MoRFchibi module is assembled hierarchically from two support vector machines that utilize RBF and Sigmoid kernels and are designed to identify MoRFs based on the contrast of physicochemical properties of MoRFs and their flanking regions. ESpritz1 is used with the (D) option parameter to predict long disordered protein regions. The hierarchical structure of the predictor generates more balanced scores such that MoRF sequences used in the training will not get too high scores that overshadow or obfuscate the ones of novel MoRF sequences. |

| | |
|---|---|
| **MorfChibi-web (Joerg Gsponer)** | MoRFchibi_web scores are computed using Hierarchical Learning [PMID: 26517836, 30952844]. Specifically, the scores of three modules (MoRFchibi, ESpritz, and ICS) are combined hierarchically using Bayes rule to generate the MoRFchibi_web score. The MoRFchibi module is assembled hierarchically from two support vector machines that utilize RBF and Sigmoid kernels and are designed to identify MoRFs based on the contrast of physicochemical properties of MoRFs and their flanking regions. ESpritz1 is used with the (D) option parameter to predict long disordered protein regions. ICS predicts MoRF regions based on conservation information using PSI-BLAST PSSM matrices. The hierarchical structure of the predictor generates more balanced scores such that MoRF sequences used in the training will not get too high scores that overshadow or obfuscate the ones of novel MoRF sequences. |
| **OPAL (Alok Sharma)** | OPAL is an ensemble of two predictors, MoRFchibi and PROMIS. PROMIS is trained using the structural information of the disordered protein sequences and MoRFchibi uses the physicochemical properties of the disordered protein sequences. StructMoRF and BigramMoRF framework is used to extract features from the MoRF residues and the neighboring amino acids upstream and downstream of the MoRF region. Successive feature selection scheme in the forward direction is performed and a support vector machine (SVM) classifier is used for prediction. OPAL provided a significant performance improvement over the benched marked predictors. The datasets used are obtained from Disfani et al.,(2012) and Malhis et al., (2015). TRAIN set is used to train the model; TEST set is used for evaluation and EXP53 set for validating the performance. |
| **PreDisorder (Jianlin Cheng)** | The bidirectional recurrent neural networks are used to predict disordered residues. The predicted disorder probabilities are scaled to make the ratio of predicted disordered residues is similar to that in the training data. |
| **pyHCA (Isabelle Callebaut)** | The pyHCA disorder predictor uses a topology-based approach considering hydrophobic and hydrophilic runs within HCA-based hydrophobic clusters. The approach only uses the information contained in a single protein sequence, i.e. without homology information such as that provided by MSAs. Physico-chemical features were extracted from the hydrophobic clusters and used to train a support vector classifier for the residue disordered states. Training was setup using a 5 fold cross-validation methodology on a splitted Disprot7 database (train/validation/test, 0.8/0.1/0.1).The set of features was selected based on prediction comparisons on the validation set. This approach is particularly well suited for small datasets or proteins without homologs as no large datasets are required for training. |
| **rawMSA (Björn Wallner)** | rawMSA is a suite of methods for the prediction of structural features of proteins. Here, the input is not a set of pre-determined features (such as evolutionary profiles or predicted secondary structure) as is common in classical ML-based methods. Instead, the whole MSA is used as a textual input to the neural network so that the evolutionary information is not compressed in any way and the feature extraction can be automatically performed by the deep network. The mapping between amino acid letters to floating point vectors is done in the first layer of the deep network with an embedding layer, as is done in Natural Language Processing (NLP) techniques. The input is not split into windows, and all predictions are done at the same time for all input amino acids so that long disordered / ordered regions are not split up at prediction stage. |
| **s2D (Michele Vendruscolo)** | s2D predicts secondary-structure populations from amino acid sequences, which simultaneously characterizes structure and disorder in a unified statistical mechanics framework. This method is based on advances made in the analysis of NMR chemical shifts that provide quantitative information about the probability distributions of secondary-structure elements in disordered states. s2D predicts secondary-structure populations with an average error of about 14%. |

| | |
|---|---|
| **s2D-2 (Michele Vendruscolo)** | s2D-2 is a re-trained and improved version of the s2D predictor. |
| **SPOT-Disorder1 (Yaoqi Zhou)** | SPOT-Disorder implements deep bidirectional LSTM recurrent neural networks in the problem of protein intrinsic disorder prediction. Its results improve on a similar method using a traditional, window-based neural network (SPINE-D) in all datasets tested without separate training on short and long disordered regions. |
| **SPOT-Disorder-Single (Yaoqi Zhou)** | SPOT-Disorder single is based on the same architecture of SPOT-Disorder a deep bidirectional LSTM recurrent neural networks. However applies a single threshold to prediction scores. |
| **SPOT-Disorder2 (Yaoqi Zhou)** | SPOT-Disorder2 improves on SPOT-Disorder by the use of an ensemble of IncReSeNet, LSTM, and FC network topologies, rather than a single LSTM topology in the previous version. Further advancements were: multiple inception-style pathways and signal Squeeze-and-Excitation, an updated feature set from our previous work to include the latest state-of-the-art predictions for protein secondary structure from SPOT-1D. |
| **VSL2B (Zoran Obradovic)** | PONDR® VSL2 predictor is a combination of neural network predictors for both short and long disordered regions. A length limit of 30 residues divides short and long disordered regions. Each individual predictor is trained by the dataset containing sequences of that specific length. The final prediction is a weighted average determined by a second layer predictor. PONDR® VSL2 applies not only to the sequence profile, but also the result of sequence alignments from PSI-BLAST and secondary structure prediction from PHD and PSIPRED. This predictor is so far the most accurate predictor in the PONDR® family. |

**Supplementary Table 3.** Name, Principal Investigator authors and brief description of all participating methods for **disorder and binding site** prediction.

### Evaluation metrics

| Label | Meaning | Formula |
|---|---|---|
| bac | Balanced accuracy | $(\frac{TP}{TP+FN} + \frac{TN}{TN+FP})/2$ |
| f1s | F1-score | $F\beta = (1 + \beta^2 \frac{ppv \ \ tpr}{(\beta^2 \cdot ppv)+tpr})$ where $\beta = 1$ |
| fpr | False Positive Rate / fall-out | $\frac{FP}{FP+TN}$ |
| mcc | Matthew's Correlation Coefficient | $\frac{TP\times TN - FP\times FN}{\sqrt{(TP+FP)(TP+FN)(TN+FP)(TN+FN)}}$ |
| ppv | Positive Predictive Value / Precision | $\frac{TP}{TP+FP}$ |
| tpr | True Positive Rate / Recall / Sensitivity | $\frac{TP}{TP+FN}$ |

| | | |
|---|---|---|
| tnr | True Negative Rate / Specificity / Selectivity | $\frac{TN}{TN+FP}$ |
| auc | Area under the ROC curve (trapezoidal rule) | $\int_a^b f(x)\,dx \approx \sum_{k=1}^{N} \frac{f(x_{k-1})+f(x_k)}{2} \Delta x_k$ |
| aps | Average Precision Score | $\sum_n (rec_n - rec_{n-1}) ppv_n$ |

**Supplementary Table 4** List of evaluation metrics used in the analysis, their acronym and how they are calculated.

Where:

- TP: true positives
- FP: false positives
- TN: true negatives
- FN: false negatives
- β: 0.5, 1, 2 for F0.5, F1 and F2 scores respectively
- ROC: Receiver Operating Characteristic
- PR: Precision Recall
- $P_n$: Precision at threshold n
- $R_n$: Recall at threshold n

### DisProt dataset

#### Evaluation Results

| | BAC | F1-S | FPR | MCC | PPV | TPR | TNR | COV |
|---|---|---|---|---|---|---|---|---|
| SPOT-Disorder2 | 0.712 | 0.486 | 0.325 | 0.308 | 0.447 | 0.734 | 0.675 | 610 |
| SPOT-Disorder1 | 0.706 | 0.475 | 0.368 | 0.295 | 0.424 | 0.761 | 0.632 | 644 |
| RawMSA | 0.692 | 0.449 | 0.281 | 0.290 | 0.439 | 0.655 | 0.719 | 646 |
| AUCpreD | 0.704 | 0.466 | 0.370 | 0.283 | 0.409 | 0.762 | 0.630 | 644 |
| DISOPRED-3.1 | 0.674 | 0.427 | 0.319 | 0.267 | 0.421 | 0.665 | 0.681 | 646 |
| Predisorder | 0.671 | 0.429 | 0.260 | 0.263 | 0.425 | 0.601 | 0.740 | 642 |
| IUPred2A-short | 0.674 | 0.424 | 0.280 | 0.256 | 0.407 | 0.625 | 0.720 | 646 |
| IUPred-short | 0.675 | 0.424 | 0.285 | 0.256 | 0.405 | 0.632 | 0.715 | 645 |
| AUCpreD-np | 0.681 | 0.441 | 0.328 | 0.254 | 0.401 | 0.679 | 0.672 | 646 |
| SPOT-Disorder-Single | 0.676 | 0.440 | 0.349 | 0.251 | 0.401 | 0.686 | 0.651 | 646 |
| MobiDB-lite | 0.668 | 0.423 | 0.289 | 0.247 | 0.400 | 0.619 | 0.711 | 645 |
| fIDPnn | 0.668 | 0.440 | 0.308 | 0.247 | 0.416 | 0.627 | 0.692 | 645 |
| IsUnstruct | 0.667 | 0.425 | 0.304 | 0.244 | 0.401 | 0.632 | 0.696 | 646 |
| IUPred-long | 0.654 | 0.395 | 0.263 | 0.243 | 0.432 | 0.569 | 0.737 | 645 |

| | | | | | | | | |
|---|---|---|---|---|---|---|---|---|
| ESpritz-X | 0.669 | 0.427 | 0.327 | 0.241 | 0.388 | 0.658 | 0.673 | 645 |
| VSL2B | 0.663 | 0.421 | 0.320 | 0.240 | 0.395 | 0.639 | 0.680 | 644 |
| IUPred2A-long | 0.654 | 0.396 | 0.268 | 0.240 | 0.424 | 0.573 | 0.732 | 646 |
| JRONN | 0.657 | 0.404 | 0.282 | 0.238 | 0.402 | 0.598 | 0.718 | 645 |
| ESpritz-N | 0.647 | 0.389 | 0.237 | 0.236 | 0.413 | 0.539 | 0.763 | 645 |
| fIDPlr | 0.647 | 0.409 | 0.297 | 0.220 | 0.420 | 0.575 | 0.703 | 645 |
| DynaMine | 0.654 | 0.400 | 0.325 | 0.220 | 0.373 | 0.630 | 0.675 | 645 |
| DisoMine | 0.643 | 0.420 | 0.410 | 0.206 | 0.388 | 0.673 | 0.590 | 646 |
| **PDB Close** | 0.598 | 0.383 | 0.365 | 0.199 | 0.404 | 0.561 | 0.635 | 604 |
| PyHCA | 0.640 | 0.390 | 0.345 | 0.198 | 0.370 | 0.618 | 0.655 | 646 |
| S2D-1 | 0.610 | 0.350 | 0.280 | 0.190 | 0.377 | 0.514 | 0.720 | 644 |
| **Gene3D** | 0.630 | 0.405 | 0.474 | 0.188 | 0.380 | 0.709 | 0.526 | 652 |
| DisEMBL-465 | 0.608 | 0.357 | 0.218 | 0.180 | 0.383 | 0.446 | 0.782 | 644 |
| S2D-2 | 0.618 | 0.365 | 0.398 | 0.173 | 0.336 | 0.634 | 0.602 | 644 |
| **PDB Remote** | 0.600 | 0.337 | 0.403 | 0.172 | 0.343 | 0.607 | 0.597 | 530 |
| FoldUnfold | 0.620 | 0.386 | 0.382 | 0.169 | 0.383 | 0.607 | 0.618 | 621 |
| ESpritz-D | 0.632 | 0.400 | 0.435 | 0.167 | 0.359 | 0.672 | 0.565 | 645 |
| **PDB observed** | 0.609 | 0.428 | 0.507 | 0.164 | 0.408 | 0.697 | 0.493 | 652 |
| GlobPlot | 0.570 | 0.300 | 0.230 | 0.136 | 0.353 | 0.397 | 0.770 | 645 |
| DisEMBL-HL | 0.563 | 0.269 | 0.124 | 0.135 | 0.371 | 0.284 | 0.876 | 644 |
| **Conservation** | 0.558 | 0.285 | 0.380 | 0.116 | 0.332 | 0.516 | 0.620 | 652 |
| DisPredict-2 | 0.557 | 0.310 | 0.437 | 0.060 | 0.300 | 0.530 | 0.563 | 646 |
| DFLpred | 0.473 | 0.033 | 0.009 | 0.020 | 0.286 | 0.022 | 0.991 | 646 |

**Supplementary Table 4**. Performance of predictors and baselines for *DisProt* dataset. Metrics are averaged over targets (proteins), sorted by MCC and predictor thresholds are optimized on MCC. Baselines are shown in bold. COV is coverage, i.e. number of predicted target proteins.

| | BAC | F1-S | FPR | MCC | PPV | TPR | TNR | COV |
|---|---|---|---|---|---|---|---|---|
| fIDPnn | 0.720 | 0.483 | 0.189 | 0.370 | 0.392 | 0.629 | 0.811 | 645 |
| SPOT-Disorder2 | 0.725 | 0.469 | 0.343 | 0.349 | 0.333 | 0.794 | 0.657 | 610 |
| fIDPlr | 0.693 | 0.452 | 0.184 | 0.330 | 0.374 | 0.570 | 0.816 | 645 |
| SPOT-Disorder1 | 0.723 | 0.434 | 0.386 | 0.330 | 0.294 | 0.832 | 0.614 | 644 |
| RawMSA | 0.714 | 0.445 | 0.297 | 0.328 | 0.321 | 0.725 | 0.703 | 646 |
| AUCpreD | 0.712 | 0.433 | 0.376 | 0.318 | 0.297 | 0.801 | 0.624 | 644 |
| SPOT-Disorder-Single | 0.710 | 0.432 | 0.341 | 0.315 | 0.302 | 0.760 | 0.659 | 646 |
| ESpritz-D | 0.703 | 0.428 | 0.325 | 0.307 | 0.303 | 0.731 | 0.675 | 645 |
| AUCpreD-np | 0.699 | 0.424 | 0.327 | 0.301 | 0.300 | 0.725 | 0.673 | 646 |
| Predisorder | 0.691 | 0.435 | 0.280 | 0.301 | 0.324 | 0.661 | 0.720 | 642 |

| | | | | | | | | |
|---|---|---|---|---|---|---|---|---|
| DisoMine | 0.698 | 0.424 | 0.326 | 0.299 | 0.300 | 0.721 | 0.674 | 646 |
| IUPred2A-short | 0.688 | 0.420 | 0.297 | 0.290 | 0.305 | 0.674 | 0.703 | 646 |
| MobiDB-lite | 0.688 | 0.420 | 0.296 | 0.289 | 0.305 | 0.673 | 0.704 | 645 |
| IsUnstruct | 0.689 | 0.418 | 0.311 | 0.288 | 0.300 | 0.688 | 0.689 | 646 |
| ESpritz-X | 0.689 | 0.418 | 0.309 | 0.288 | 0.301 | 0.686 | 0.691 | 645 |
| IUPred-short | 0.688 | 0.418 | 0.304 | 0.288 | 0.302 | 0.679 | 0.696 | 645 |
| IUPred-long | 0.686 | 0.418 | 0.294 | 0.287 | 0.305 | 0.666 | 0.706 | 645 |
| IUPred2A-long | 0.685 | 0.416 | 0.299 | 0.285 | 0.302 | 0.670 | 0.701 | 646 |
| VSL2B | 0.684 | 0.408 | 0.341 | 0.277 | 0.286 | 0.709 | 0.659 | 644 |
| JRONN | 0.672 | 0.401 | 0.318 | 0.263 | 0.287 | 0.663 | 0.682 | 645 |
| ESpritz-N | 0.664 | 0.400 | 0.271 | 0.259 | 0.300 | 0.599 | 0.729 | 645 |
| DISOPRED-3.1 | 0.674 | 0.393 | 0.401 | 0.258 | 0.266 | 0.749 | 0.599 | 646 |
| PyHCA | 0.660 | 0.385 | 0.346 | 0.241 | 0.271 | 0.666 | 0.654 | 646 |
| DynaMine | 0.660 | 0.384 | 0.362 | 0.240 | 0.267 | 0.682 | 0.638 | 645 |
| **Gene3D** | 0.653 | 0.368 | 0.486 | 0.226 | 0.240 | 0.791 | 0.514 | 652 |
| DisEMBL-465 | 0.627 | 0.363 | 0.215 | 0.214 | 0.296 | 0.468 | 0.785 | 644 |
| FoldUnfold | 0.642 | 0.365 | 0.382 | 0.211 | 0.251 | 0.666 | 0.618 | 621 |
| S2D-1 | 0.633 | 0.361 | 0.329 | 0.203 | 0.259 | 0.595 | 0.671 | 644 |
| **PDB Close** | 0.637 | 0.353 | 0.380 | 0.202 | 0.242 | 0.655 | 0.620 | 604 |
| S2D-2 | 0.624 | 0.347 | 0.439 | 0.183 | 0.232 | 0.687 | 0.561 | 644 |
| **PDB observed** | 0.616 | 0.339 | 0.565 | 0.174 | 0.215 | 0.796 | 0.435 | 652 |
| DisEMBL-HL | 0.577 | 0.286 | 0.099 | 0.172 | 0.330 | 0.253 | 0.901 | 644 |
| **PDB Remote** | 0.614 | 0.321 | 0.450 | 0.163 | 0.210 | 0.678 | 0.550 | 530 |
| DisPredict-2 | 0.599 | 0.326 | 0.326 | 0.152 | 0.237 | 0.523 | 0.674 | 646 |
| GlobPlot | 0.587 | 0.312 | 0.253 | 0.143 | 0.246 | 0.427 | 0.747 | 645 |
| **Conservation** | 0.552 | 0.288 | 0.483 | 0.077 | 0.191 | 0.587 | 0.517 | 652 |
| DFLpred | 0.503 | 0.025 | 0.008 | 0.022 | 0.249 | 0.013 | 0.992 | 646 |

**Supplementary Table 5**. Performance of predictors and baselines for *DisProt* dataset. Metrics are calculated over the whole dataset, sorted by MCC and predictors threshold are optimized on MCC. Baselines are shown in bold. COV is coverage, i.e. number of predicted target proteins.

#### Evaluation Results (Full ID)

| Fully disordered proteins (ID fraction ≥80%) | | | | | | | | | | |
|---|---|---|---|---|---|---|---|---|---|---|
| | **TN** | **FP** | **FN** | **TP** | **MCC** | **F1-S** | **TNR** | **TPR** | **PPV** | **BAC** |
| fIDPnn | 569 | 21 | 24 | 32 | 0.549 | 0.587 | 0.964 | 0.571 | 0.604 | 0.768 |
| fIDPlr | 544 | 46 | 17 | 39 | 0.515 | 0.553 | 0.922 | 0.696 | 0.459 | 0.809 |
| RawMSA | 553 | 37 | 21 | 35 | 0.503 | 0.547 | 0.937 | 0.625 | 0.486 | 0.781 |
| AUCpreD | 568 | 22 | 28 | 28 | 0.487 | 0.528 | 0.963 | 0.500 | 0.560 | 0.731 |

| | | | | | | | | | | |
|---|---|---|---|---|---|---|---|---|---|---|
| SPOT-Disorder2 | 539 | 51 | 19 | 37 | 0.471 | 0.514 | 0.914 | 0.661 | 0.420 | 0.787 |
| PyHCA | 562 | 28 | 27 | 29 | 0.467 | 0.513 | 0.953 | 0.518 | 0.509 | 0.735 |
| IUPred2A-long | 564 | 26 | 28 | 28 | 0.464 | 0.509 | 0.956 | 0.500 | 0.519 | 0.728 |
| ESpritz-N | 565 | 25 | 29 | 27 | 0.455 | 0.500 | 0.958 | 0.482 | 0.519 | 0.720 |
| SPOT-Disorder-Single | 577 | 13 | 34 | 22 | 0.461 | 0.484 | 0.978 | 0.393 | 0.629 | 0.685 |
| JRONN | 558 | 32 | 28 | 28 | 0.432 | 0.483 | 0.946 | 0.500 | 0.467 | 0.723 |
| Predisorder | 528 | 62 | 19 | 37 | 0.434 | 0.477 | 0.895 | 0.661 | 0.374 | 0.778 |
| IUPred2A-short | 585 | 5 | 37 | 19 | 0.492 | 0.475 | 0.992 | 0.339 | 0.792 | 0.665 |
| DisoMine | 516 | 74 | 17 | 39 | 0.423 | 0.462 | 0.875 | 0.696 | 0.345 | 0.786 |
| VSL2B | 516 | 74 | 18 | 38 | 0.411 | 0.452 | 0.875 | 0.679 | 0.339 | 0.777 |
| AUCpreD-np | 577 | 13 | 36 | 20 | 0.428 | 0.449 | 0.978 | 0.357 | 0.606 | 0.668 |
| MobiDB-lite | 586 | 4 | 39 | 17 | 0.471 | 0.442 | 0.993 | 0.304 | 0.810 | 0.648 |
| ESpritz-D | 533 | 57 | 24 | 32 | 0.388 | 0.441 | 0.903 | 0.571 | 0.360 | 0.737 |
| IsUnstruct | 548 | 42 | 29 | 27 | 0.374 | 0.432 | 0.929 | 0.482 | 0.391 | 0.705 |
| ESpritz-X | 576 | 14 | 37 | 19 | 0.403 | 0.427 | 0.976 | 0.339 | 0.576 | 0.658 |
| DisPredict-2 | 538 | 52 | 29 | 27 | 0.338 | 0.400 | 0.912 | 0.482 | 0.342 | 0.697 |
| **Gene3D** | 464 | 126 | 11 | 45 | 0.376 | 0.396 | 0.786 | 0.804 | 0.263 | 0.795 |
| DisEMBL-HL | 580 | 10 | 43 | 13 | 0.327 | 0.329 | 0.983 | 0.232 | 0.565 | 0.608 |
| FoldUnfold | 451 | 139 | 19 | 37 | 0.269 | 0.319 | 0.764 | 0.661 | 0.210 | 0.713 |
| **PDB observed** | 407 | 183 | 13 | 43 | 0.270 | 0.305 | 0.690 | 0.768 | 0.190 | 0.729 |
| S2D-2 | 454 | 136 | 23 | 33 | 0.230 | 0.293 | 0.769 | 0.589 | 0.195 | 0.679 |
| DISOPRED-3.1 | 568 | 22 | 44 | 12 | 0.223 | 0.267 | 0.963 | 0.214 | 0.353 | 0.588 |
| **Conservation** | 175 | 415 | 18 | 38 | -0.015 | 0.149 | 0.297 | 0.679 | 0.084 | 0.488 |
| DisEMBL-465 | 590 | 0 | 52 | 4 | 0.256 | 0.133 | 1.000 | 0.071 | 1.000 | 0.536 |
| **PDB Close** | 540 | 50 | 49 | 7 | 0.040 | 0.124 | 0.915 | 0.125 | 0.123 | 0.520 |
| **PDB Remote** | 542 | 48 | 51 | 5 | 0.008 | 0.092 | 0.919 | 0.089 | 0.094 | 0.504 |
| GlobPlot | 589 | 1 | 55 | 1 | 0.082 | 0.034 | 0.998 | 0.018 | 0.500 | 0.508 |
| DFLpred | 590 | 0 | 56 | 0 | 0.000 | 0.000 | 1.000 | 0.000 | 0.000 | 0.500 |
| DynaMine | 590 | 0 | 56 | 0 | 0.000 | 0.000 | 1.000 | 0.000 | 0.000 | 0.500 |

**Supplementary Table 6.** Performance of predictors and baselines in the task of identifying fully disordered proteins sorted by F1-Score. Where a fully disordered protein is defined as protein with 80% or more residues annotated / predicted as disordered. Baselines are shown in bold.

| Fully disordered proteins (ID fraction ≥90%) | | | | | | | | | | |
|---|---|---|---|---|---|---|---|---|---|---|
| | **TN** | **FP** | **FN** | **TP** | **MCC** | **F1-S** | **TNR** | **TPR** | **PPV** | **BAC** |
| fIDPnn | 580 | 17 | 19 | 30 | 0.595 | 0.625 | 0.972 | 0.612 | 0.638 | 0.792 |
| RawMSA | 572 | 25 | 18 | 31 | 0.556 | 0.590 | 0.958 | 0.633 | 0.554 | 0.795 |
| IUPred2A-long | 590 | 7 | 28 | 21 | 0.542 | 0.545 | 0.988 | 0.429 | 0.750 | 0.708 |

| | | | | | | | | | | |
|---|---|---|---|---|---|---|---|---|---|---|
| fIDPlr | 557 | 40 | 18 | 31 | 0.479 | 0.517 | 0.933 | 0.633 | 0.437 | 0.783 |
| PyHCA | 591 | 6 | 30 | 19 | 0.518 | 0.514 | 0.990 | 0.388 | 0.760 | 0.689 |
| JRONN | 588 | 9 | 29 | 20 | 0.503 | 0.513 | 0.985 | 0.408 | 0.690 | 0.697 |
| Predisorder | 576 | 21 | 25 | 24 | 0.473 | 0.511 | 0.965 | 0.490 | 0.533 | 0.727 |
| SPOT-Disorder-Single | 589 | 8 | 30 | 19 | 0.495 | 0.500 | 0.987 | 0.388 | 0.704 | 0.687 |
| AUCpreD | 585 | 12 | 29 | 20 | 0.474 | 0.494 | 0.980 | 0.408 | 0.625 | 0.694 |
| ESpritz-N | 588 | 9 | 30 | 19 | 0.485 | 0.494 | 0.985 | 0.388 | 0.679 | 0.686 |
| VSL2B | 556 | 41 | 20 | 29 | 0.446 | 0.487 | 0.931 | 0.592 | 0.414 | 0.762 |
| DisoMine | 540 | 57 | 17 | 32 | 0.428 | 0.464 | 0.905 | 0.653 | 0.360 | 0.779 |
| IsUnstruct | 579 | 18 | 29 | 20 | 0.425 | 0.460 | 0.970 | 0.408 | 0.526 | 0.689 |
| SPOT-Disorder2 | 562 | 35 | 24 | 25 | 0.412 | 0.459 | 0.941 | 0.510 | 0.417 | 0.726 |
| **Gene3D** | 499 | 98 | 10 | 39 | 0.409 | 0.419 | 0.836 | 0.796 | 0.285 | 0.816 |
| ESpritz-D | 546 | 51 | 24 | 25 | 0.349 | 0.400 | 0.915 | 0.510 | 0.329 | 0.712 |
| MobiDB-lite | 594 | 3 | 36 | 13 | 0.443 | 0.400 | 0.995 | 0.265 | 0.812 | 0.630 |
| DisPredict-2 | 573 | 24 | 31 | 18 | 0.351 | 0.396 | 0.960 | 0.367 | 0.429 | 0.664 |
| ESpritz-X | 589 | 8 | 35 | 14 | 0.398 | 0.394 | 0.987 | 0.286 | 0.636 | 0.636 |
| AUCpreD-np | 584 | 13 | 34 | 15 | 0.370 | 0.390 | 0.978 | 0.306 | 0.536 | 0.642 |
| IUPred2A-short | 593 | 4 | 37 | 12 | 0.406 | 0.369 | 0.993 | 0.245 | 0.750 | 0.619 |
| S2D-2 | 538 | 59 | 28 | 21 | 0.265 | 0.326 | 0.901 | 0.429 | 0.262 | 0.665 |
| **PDB observed** | 449 | 148 | 13 | 36 | 0.286 | 0.309 | 0.752 | 0.735 | 0.196 | 0.743 |
| DisEMBL-HL | 596 | 1 | 40 | 9 | 0.390 | 0.305 | 0.998 | 0.184 | 0.900 | 0.591 |
| FoldUnfold | 455 | 142 | 15 | 34 | 0.271 | 0.302 | 0.762 | 0.694 | 0.193 | 0.728 |
| DISOPRED-3.1 | 589 | 8 | 41 | 8 | 0.255 | 0.246 | 0.987 | 0.163 | 0.500 | 0.575 |
| DisEMBL-465 | 597 | 0 | 46 | 3 | 0.238 | 0.115 | 1.000 | 0.061 | 1.000 | 0.531 |
| **Conservation** | 314 | 283 | 29 | 20 | -0.035 | 0.114 | 0.526 | 0.408 | 0.066 | 0.467 |
| **PDB Remote** | 576 | 21 | 45 | 4 | 0.064 | 0.108 | 0.965 | 0.082 | 0.160 | 0.523 |
| **PDB Close** | 576 | 21 | 46 | 3 | 0.036 | 0.082 | 0.965 | 0.061 | 0.125 | 0.513 |
| DynaMine | 597 | 0 | 49 | 0 | 0.000 | 0.000 | 1.000 | 0.000 | 0.000 | 0.500 |
| GlobPlot | 597 | 0 | 49 | 0 | 0.000 | 0.000 | 1.000 | 0.000 | 0.000 | 0.500 |
| DFLpred | 597 | 0 | 49 | 0 | 0.000 | 0.000 | 1.000 | 0.000 | 0.000 | 0.500 |

**Supplementary Table 7.** Performance of predictors and baselines in the task of identifying fully disordered proteins sorted by F1-Score. Where a fully disordered protein is defined as protein with 90% or more residues annotated / predicted as disordered. Baselines are shown in bold.

| Fully disordered proteins (ID fraction ≥99%) | | | | | | | | | | |
|---|---|---|---|---|---|---|---|---|---|---|
| | **TN** | **FP** | **FN** | **TP** | **MCC** | **F1-S** | **TNR** | **TPR** | **PPV** | **BAC** |
| RawMSA | 588 | 13 | 20 | 25 | 0.578 | 0.602 | 0.978 | 0.556 | 0.658 | 0.767 |
| fIDPnn | 587 | 14 | 22 | 23 | 0.534 | 0.561 | 0.977 | 0.511 | 0.622 | 0.744 |

| | | | | | | | | | | |
|---|---|---|---|---|---|---|---|---|---|---|
| VSL2B | 583 | 18 | 22 | 23 | 0.502 | 0.535 | 0.970 | 0.511 | 0.561 | 0.741 |
| fIDPlr | 572 | 29 | 20 | 25 | 0.467 | 0.505 | 0.952 | 0.556 | 0.463 | 0.754 |
| DisoMine | 557 | 44 | 18 | 27 | 0.429 | 0.466 | 0.927 | 0.600 | 0.380 | 0.763 |
| Predisorder | 596 | 5 | 30 | 15 | 0.478 | 0.462 | 0.992 | 0.333 | 0.750 | 0.663 |
| IsUnstruct | 593 | 8 | 30 | 15 | 0.440 | 0.441 | 0.987 | 0.333 | 0.652 | 0.660 |
| AUCpreD | 589 | 12 | 29 | 16 | 0.420 | 0.438 | 0.980 | 0.356 | 0.571 | 0.668 |
| SPOT-Disorder2 | 576 | 25 | 26 | 19 | 0.385 | 0.427 | 0.958 | 0.422 | 0.432 | 0.690 |
| IUPred2A-long | 598 | 3 | 32 | 13 | 0.465 | 0.426 | 0.995 | 0.289 | 0.812 | 0.642 |
| SPOT-Disorder-Single | 594 | 7 | 31 | 14 | 0.430 | 0.424 | 0.988 | 0.311 | 0.667 | 0.650 |
| ESpritz-N | 599 | 2 | 33 | 12 | 0.460 | 0.407 | 0.997 | 0.267 | 0.857 | 0.632 |
| **Gene3D** | 505 | 96 | 10 | 35 | 0.391 | 0.398 | 0.840 | 0.778 | 0.267 | 0.809 |
| ESpritz-D | 558 | 43 | 24 | 21 | 0.337 | 0.385 | 0.928 | 0.467 | 0.328 | 0.698 |
| DisPredict-2 | 594 | 7 | 33 | 12 | 0.384 | 0.375 | 0.988 | 0.267 | 0.632 | 0.628 |
| MobiDB-lite | 599 | 2 | 35 | 10 | 0.413 | 0.351 | 0.997 | 0.222 | 0.833 | 0.609 |
| **PDB observed** | 493 | 108 | 13 | 32 | 0.328 | 0.346 | 0.820 | 0.711 | 0.229 | 0.766 |
| JRONN | 597 | 4 | 35 | 10 | 0.377 | 0.339 | 0.993 | 0.222 | 0.714 | 0.608 |
| ESpritz-X | 597 | 4 | 36 | 9 | 0.351 | 0.310 | 0.993 | 0.200 | 0.692 | 0.597 |
| AUCpreD-np | 591 | 10 | 35 | 10 | 0.302 | 0.308 | 0.983 | 0.222 | 0.500 | 0.603 |
| FoldUnfold | 456 | 145 | 14 | 31 | 0.256 | 0.281 | 0.759 | 0.689 | 0.176 | 0.724 |
| IUPred2A-short | 600 | 1 | 38 | 7 | 0.354 | 0.264 | 0.998 | 0.156 | 0.875 | 0.577 |
| S2D-2 | 597 | 4 | 38 | 7 | 0.293 | 0.250 | 0.993 | 0.156 | 0.636 | 0.574 |
| PyHCA | 601 | 0 | 39 | 6 | 0.354 | 0.235 | 1.000 | 0.133 | 1.000 | 0.567 |
| DISOPRED-3.1 | 599 | 2 | 41 | 4 | 0.227 | 0.157 | 0.997 | 0.089 | 0.667 | 0.543 |
| DisEMBL-HL | 601 | 0 | 42 | 3 | 0.250 | 0.125 | 1.000 | 0.067 | 1.000 | 0.533 |
| DisEMBL-465 | 601 | 0 | 43 | 2 | 0.204 | 0.085 | 1.000 | 0.044 | 1.000 | 0.522 |
| **PDB Close** | 598 | 3 | 43 | 2 | 0.115 | 0.080 | 0.995 | 0.044 | 0.400 | 0.520 |
| **PDB Remote** | 592 | 9 | 43 | 2 | 0.058 | 0.071 | 0.985 | 0.044 | 0.182 | 0.515 |
| **Conservation** | 590 | 11 | 43 | 2 | 0.047 | 0.069 | 0.982 | 0.044 | 0.154 | 0.513 |
| DynaMine | 601 | 0 | 45 | 0 | 0.000 | 0.000 | 1.000 | 0.000 | 0.000 | 0.500 |
| GlobPlot | 601 | 0 | 45 | 0 | 0.000 | 0.000 | 1.000 | 0.000 | 0.000 | 0.500 |
| DFLpred | 601 | 0 | 45 | 0 | 0.000 | 0.000 | 1.000 | 0.000 | 0.000 | 0.500 |

**Supplementary Table 8.** Performance of predictors and baselines in the task of identifying fully disordered proteins sorted by F1-Score. Where a fully disordered protein is defined as protein with 99% or more residues annotated / predicted as disordered. Baselines are shown in bold.

#### DisProt-PDB dataset

Evaluation Results

| | BAC | F1-S | FPR | MCC | PPV | TPR | TNR | COV |
|---|---|---|---|---|---|---|---|---|
| **PDB observed** | 0.817 | 0.751 | 0.260 | 0.426 | 0.847 | 0.844 | 0.740 | 652 |
| SPOT-Disorder2 | 0.753 | 0.643 | 0.106 | 0.356 | 0.745 | 0.651 | 0.894 | 610 |
| **PDB Close** | 0.716 | 0.624 | 0.291 | 0.341 | 0.780 | 0.712 | 0.709 | 604 |
| AUCpreD | 0.738 | 0.623 | 0.122 | 0.334 | 0.719 | 0.646 | 0.878 | 644 |
| SPOT-Disorder1 | 0.761 | 0.649 | 0.162 | 0.330 | 0.718 | 0.690 | 0.838 | 644 |
| DISOPRED-3.1 | 0.703 | 0.581 | 0.116 | 0.306 | 0.712 | 0.588 | 0.884 | 646 |
| Predisorder | 0.697 | 0.579 | 0.116 | 0.303 | 0.705 | 0.582 | 0.884 | 642 |
| RawMSA | 0.736 | 0.612 | 0.182 | 0.299 | 0.690 | 0.668 | 0.818 | 646 |
| AUCpreD-np | 0.722 | 0.601 | 0.155 | 0.297 | 0.679 | 0.637 | 0.845 | 646 |
| SPOT-Disorder-Single | 0.730 | 0.608 | 0.172 | 0.290 | 0.679 | 0.651 | 0.828 | 646 |
| MobiDB-lite | 0.673 | 0.550 | 0.120 | 0.274 | 0.692 | 0.542 | 0.880 | 645 |
| IsUnstruct | 0.691 | 0.574 | 0.157 | 0.272 | 0.674 | 0.591 | 0.843 | 646 |
| ESpritz-X | 0.712 | 0.586 | 0.195 | 0.272 | 0.646 | 0.646 | 0.805 | 645 |
| IUPred2A-short | 0.693 | 0.567 | 0.151 | 0.270 | 0.666 | 0.591 | 0.849 | 646 |
| IUPred-short | 0.700 | 0.574 | 0.162 | 0.270 | 0.660 | 0.607 | 0.838 | 645 |
| ESpritz-N | 0.661 | 0.524 | 0.103 | 0.268 | 0.678 | 0.517 | 0.897 | 645 |
| VSL2B | 0.674 | 0.549 | 0.144 | 0.264 | 0.676 | 0.556 | 0.856 | 644 |
| JRONN | 0.657 | 0.528 | 0.117 | 0.258 | 0.677 | 0.518 | 0.883 | 645 |
| fIDPnn | 0.713 | 0.591 | 0.253 | 0.252 | 0.654 | 0.673 | 0.747 | 645 |
| DynaMine | 0.657 | 0.527 | 0.139 | 0.245 | 0.643 | 0.533 | 0.861 | 645 |
| IUPred-long | 0.679 | 0.540 | 0.164 | 0.244 | 0.675 | 0.564 | 0.836 | 645 |
| **Gene3D** | 0.740 | 0.625 | 0.380 | 0.243 | 0.657 | 0.778 | 0.620 | 652 |
| IUPred2A-long | 0.669 | 0.529 | 0.151 | 0.242 | 0.677 | 0.544 | 0.849 | 646 |
| fIDPlr | 0.689 | 0.562 | 0.277 | 0.230 | 0.657 | 0.649 | 0.723 | 645 |
| PyHCA | 0.642 | 0.500 | 0.154 | 0.226 | 0.646 | 0.518 | 0.846 | 646 |
| S2D-1 | 0.603 | 0.456 | 0.124 | 0.218 | 0.642 | 0.447 | 0.876 | 644 |
| DisEMBL-465 | 0.610 | 0.485 | 0.151 | 0.209 | 0.620 | 0.476 | 0.849 | 644 |
| DisoMine | 0.693 | 0.558 | 0.323 | 0.205 | 0.622 | 0.675 | 0.677 | 646 |
| S2D-2 | 0.649 | 0.511 | 0.278 | 0.190 | 0.569 | 0.606 | 0.722 | 644 |
| **PDB Remote** | 0.662 | 0.518 | 0.345 | 0.184 | 0.595 | 0.646 | 0.655 | 530 |
| FoldUnfold | 0.665 | 0.553 | 0.330 | 0.176 | 0.618 | 0.637 | 0.670 | 621 |
| GlobPlot | 0.549 | 0.394 | 0.127 | 0.160 | 0.605 | 0.365 | 0.873 | 645 |
| ESpritz-D | 0.670 | 0.534 | 0.374 | 0.152 | 0.581 | 0.666 | 0.626 | 645 |
| **Conservation** | 0.557 | 0.396 | 0.261 | 0.135 | 0.565 | 0.464 | 0.739 | 652 |
| DisEMBL-HL | 0.661 | 0.531 | 0.523 | 0.132 | 0.514 | 0.763 | 0.477 | 644 |
| DisPredict-2 | 0.580 | 0.435 | 0.401 | 0.061 | 0.519 | 0.543 | 0.599 | 646 |
| DFLpred | 0.368 | 0.046 | 0.009 | 0.024 | 0.515 | 0.029 | 0.991 | 646 |

**Supplementary Table 9.** Performance of predictors and baselines for *DisProt-PDB* dataset. Metrics are averaged over targets (proteins) and sorted by MCC. Predictors thresholds are optimized on MCC. Baselines are shown in bold. COV is coverage, i.e. number of predicted target proteins.

| | BAC | F1-S | FPR | MCC | PPV | TPR | TNR | COV |
|---|---|---|---|---|---|---|---|---|
| **PDB observed** | 0.898 | 0.886 | 0.000 | 0.854 | 1.000 | 0.796 | 1.000 | 652 |
| SPOT-Disorder2 | 0.836 | 0.784 | 0.055 | 0.706 | 0.851 | 0.727 | 0.945 | 610 |
| SPOT-Disorder1 | 0.846 | 0.788 | 0.090 | 0.696 | 0.795 | 0.782 | 0.910 | 644 |
| **PDB Close** | 0.811 | 0.755 | 0.033 | 0.689 | 0.891 | 0.655 | 0.967 | 604 |
| AUCpreD | 0.816 | 0.756 | 0.070 | 0.662 | 0.820 | 0.701 | 0.930 | 644 |
| SPOT-Disorder-Single | 0.817 | 0.751 | 0.095 | 0.646 | 0.775 | 0.729 | 0.905 | 646 |
| RawMSA | 0.815 | 0.745 | 0.106 | 0.635 | 0.755 | 0.736 | 0.894 | 646 |
| Predisorder | 0.788 | 0.717 | 0.067 | 0.619 | 0.813 | 0.642 | 0.933 | 642 |
| AUCpreD-np | 0.797 | 0.725 | 0.092 | 0.615 | 0.769 | 0.686 | 0.908 | 646 |
| DISOPRED-3.1 | 0.796 | 0.724 | 0.092 | 0.613 | 0.768 | 0.684 | 0.908 | 646 |
| IUPred-long | 0.783 | 0.704 | 0.096 | 0.588 | 0.754 | 0.661 | 0.904 | 645 |
| IsUnstruct | 0.779 | 0.700 | 0.091 | 0.585 | 0.760 | 0.648 | 0.909 | 646 |
| IUPred2A-long | 0.776 | 0.697 | 0.087 | 0.584 | 0.766 | 0.640 | 0.913 | 646 |
| MobiDB-lite | 0.764 | 0.683 | 0.063 | 0.583 | 0.806 | 0.592 | 0.937 | 645 |
| VSL2B | 0.774 | 0.695 | 0.087 | 0.581 | 0.765 | 0.636 | 0.913 | 644 |
| fIDPnn | 0.782 | 0.701 | 0.113 | 0.576 | 0.727 | 0.676 | 0.887 | 645 |
| IUPred2A-short | 0.773 | 0.691 | 0.094 | 0.574 | 0.752 | 0.640 | 0.906 | 646 |
| IUPred-short | 0.775 | 0.693 | 0.104 | 0.571 | 0.738 | 0.654 | 0.896 | 645 |
| ESpritz-X | 0.778 | 0.695 | 0.119 | 0.566 | 0.717 | 0.675 | 0.881 | 645 |
| ESpritz-N | 0.751 | 0.662 | 0.073 | 0.554 | 0.779 | 0.575 | 0.927 | 645 |
| DisoMine | 0.780 | 0.693 | 0.160 | 0.550 | 0.668 | 0.721 | 0.840 | 646 |
| JRONN | 0.751 | 0.661 | 0.081 | 0.546 | 0.762 | 0.583 | 0.919 | 645 |
| ESpritz-D | 0.778 | 0.690 | 0.166 | 0.544 | 0.660 | 0.723 | 0.834 | 645 |
| **Gene3D** | 0.785 | 0.692 | 0.220 | 0.539 | 0.615 | 0.791 | 0.780 | 652 |
| fIDPlr | 0.761 | 0.671 | 0.119 | 0.537 | 0.705 | 0.641 | 0.881 | 645 |
| DynaMine | 0.739 | 0.641 | 0.110 | 0.505 | 0.704 | 0.588 | 0.890 | 645 |
| PyHCA | 0.731 | 0.629 | 0.107 | 0.494 | 0.704 | 0.569 | 0.893 | 646 |
| S2D-1 | 0.724 | 0.617 | 0.089 | 0.494 | 0.728 | 0.536 | 0.911 | 644 |
| FoldUnfold | 0.736 | 0.636 | 0.193 | 0.462 | 0.608 | 0.666 | 0.807 | 621 |
| DisEMBL-465 | 0.694 | 0.570 | 0.110 | 0.426 | 0.667 | 0.498 | 0.890 | 644 |
| S2D-2 | 0.703 | 0.591 | 0.253 | 0.386 | 0.536 | 0.658 | 0.747 | 644 |
| **PDB Remote** | 0.703 | 0.579 | 0.273 | 0.377 | 0.505 | 0.678 | 0.727 | 530 |
| GlobPlot | 0.641 | 0.480 | 0.111 | 0.328 | 0.613 | 0.394 | 0.889 | 645 |
| DisEMBL-HL | 0.641 | 0.535 | 0.470 | 0.262 | 0.415 | 0.752 | 0.530 | 644 |

| | | | | | | | | |
|---|---|---|---|---|---|---|---|---|
| DisPredict-2 | 0.625 | 0.491 | 0.285 | 0.240 | 0.455 | 0.534 | 0.715 | 646 |
| **Conservation** | 0.618 | 0.485 | 0.296 | 0.227 | 0.445 | 0.533 | 0.704 | 652 |
| DFLpred | 0.504 | 0.027 | 0.005 | 0.043 | 0.530 | 0.014 | 0.995 | 646 |

**Supplementary Table 10**. Performance of predictors and baselines for *DisProt-PDB* dataset. Metrics are calculated over the whole dataset and sorted by MCC. Predictors thresholds are optimized on MCC. Baselines are shown in bold. COV is coverage, i.e. number of predicted target proteins.

### Disprot-Binding dataset

#### Evaluation Results

| | BAC | F1-S | FPR | MCC | PPV | TPR | TNR | COV |
|---|---|---|---|---|---|---|---|---|
| DisoRDPbind-protein | 0.652 | 0.137 | 0.357 | 0.062 | 0.139 | 0.229 | 0.643 | 646 |
| ANCHOR-2 | 0.677 | 0.130 | 0.319 | 0.055 | 0.138 | 0.213 | 0.681 | 646 |
| **Gene3D** | 0.529 | 0.143 | 0.523 | 0.053 | 0.125 | 0.261 | 0.477 | 652 |
| MoRFchibi-light | 0.629 | 0.124 | 0.339 | 0.041 | 0.125 | 0.178 | 0.661 | 644 |
| MoRFchibi-web | 0.588 | 0.131 | 0.407 | 0.039 | 0.122 | 0.205 | 0.593 | 644 |
| OPAL | 0.482 | 0.141 | 0.583 | 0.039 | 0.116 | 0.272 | 0.417 | 644 |
| DISOPRED-3.1-binding | 0.725 | 0.095 | 0.172 | 0.036 | 0.130 | 0.105 | 0.828 | 646 |
| ANCHOR | 0.571 | 0.127 | 0.462 | 0.026 | 0.112 | 0.231 | 0.538 | 645 |
| fMoRFpred | 0.790 | 0.031 | 0.033 | 0.014 | 0.128 | 0.020 | 0.967 | 646 |
| DisoRDPbind-DNA | 0.804 | 0.005 | 0.005 | 0.004 | 0.122 | 0.003 | 0.995 | 646 |
| DisoRDPbind-RNA | 0.799 | 0.007 | 0.011 | 0.002 | 0.098 | 0.005 | 0.989 | 646 |
| DisoRDPbind | 0.194 | 0.131 | 1.000 | 0.000 | 0.100 | 0.358 | 0.000 | 646 |
| **PDB observed** | 0.490 | 0.128 | 0.569 | -0.011 | 0.115 | 0.224 | 0.431 | 652 |

**Supplementary Table 11**. Performance of predictors and baselines for *DisProt-Binding* dataset. Metrics are averaged over targets (proteins) and sorted by MCC. Predictors thresholds are optimized on MCC. Baselines are shown in bold. COV is coverage, i.e. number of predicted target proteins.

| | BAC | F1-S | FPR | MCC | PPV | TPR | TNR | COV |
|---|---|---|---|---|---|---|---|---|
| ANCHOR-2 | 0.694 | 0.220 | 0.320 | 0.199 | 0.130 | 0.708 | 0.680 | 646 |
| DisoRDPbind-protein | 0.697 | 0.214 | 0.353 | 0.198 | 0.125 | 0.746 | 0.647 | 646 |
| MoRFchibi-light | 0.636 | 0.212 | 0.200 | 0.161 | 0.137 | 0.472 | 0.800 | 644 |
| **Gene3D** | 0.656 | 0.175 | 0.516 | 0.153 | 0.098 | 0.828 | 0.484 | 652 |
| OPAL | 0.652 | 0.186 | 0.374 | 0.151 | 0.108 | 0.678 | 0.626 | 644 |
| ANCHOR | 0.651 | 0.178 | 0.451 | 0.148 | 0.101 | 0.754 | 0.549 | 645 |
| MoRFchibi-web | 0.631 | 0.194 | 0.257 | 0.143 | 0.119 | 0.519 | 0.743 | 644 |
| **PDB observed** | 0.606 | 0.152 | 0.589 | 0.106 | 0.084 | 0.801 | 0.411 | 652 |

| | | | | | | | | |
|---|---|---|---|---|---|---|---|---|
| DISOPRED-3.1-binding | 0.569 | 0.169 | 0.125 | 0.099 | 0.124 | 0.263 | 0.875 | 646 |
| fMoRFpred | 0.515 | 0.072 | 0.017 | 0.054 | 0.157 | 0.047 | 0.983 | 646 |
| DisoRDPbind-DNA | 0.502 | 0.008 | 0.000 | 0.052 | 0.724 | 0.004 | 1.000 | 646 |
| DisoRDPbind-RNA | 0.501 | 0.010 | 0.002 | 0.014 | 0.136 | 0.005 | 0.998 | 646 |
| DisoRDPbind | 0.500 | 0.119 | 1.000 | 0.000 | 0.063 | 1.000 | 0.000 | 646 |

**Supplementary Table 12**. Performance of predictors and baselines for *DisProt-Binding* dataset. Metrics are calculated over the whole dataset and sorted by MCC. Predictors thresholds are optimized on MCC. Baselines are shown in bold. COV is coverage, i.e. number of predicted target proteins.

### Figures

#### Dataset

##### Annotations

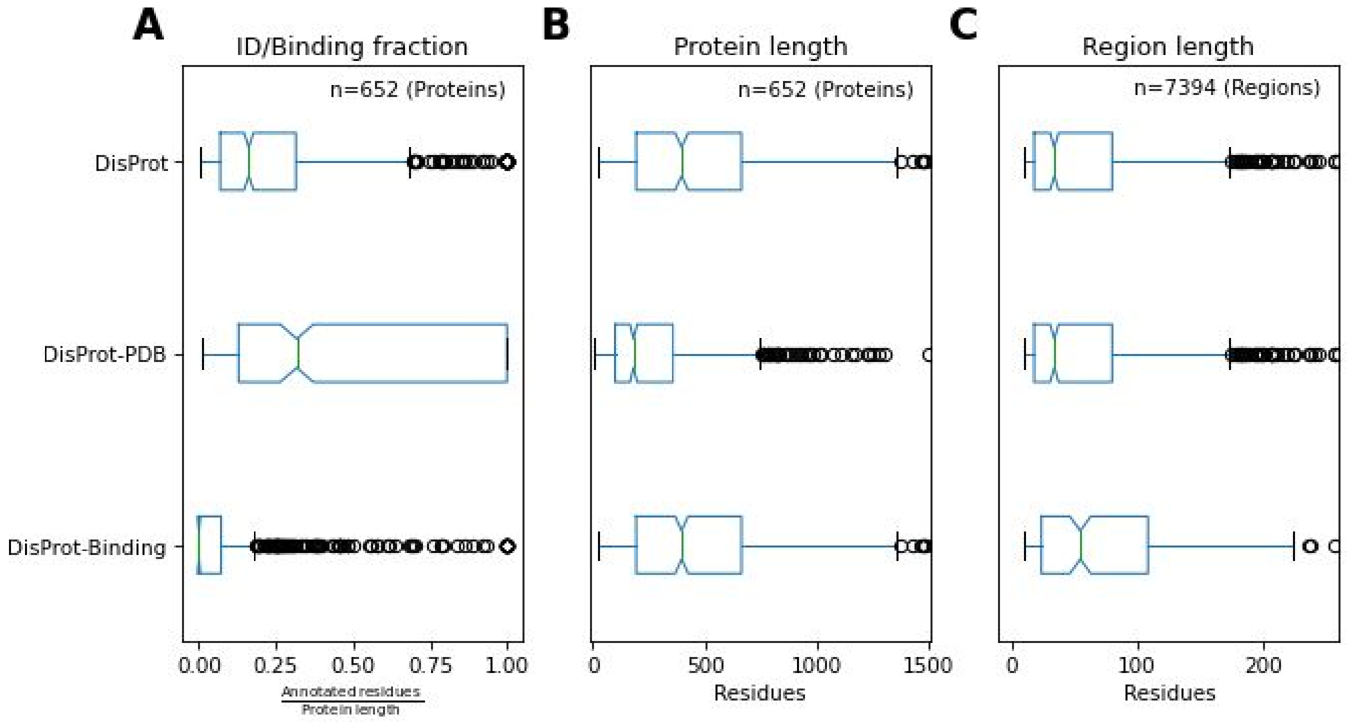


**Supplementary Figure 1** Distribution of the fraction of disordered / binding residues in each protein (panel A), of the protein lengths (panel B) and region lengths (panel C) in the three datasets (*DisProt*, *DisProt-PDB* and *DisProt-Binding*).

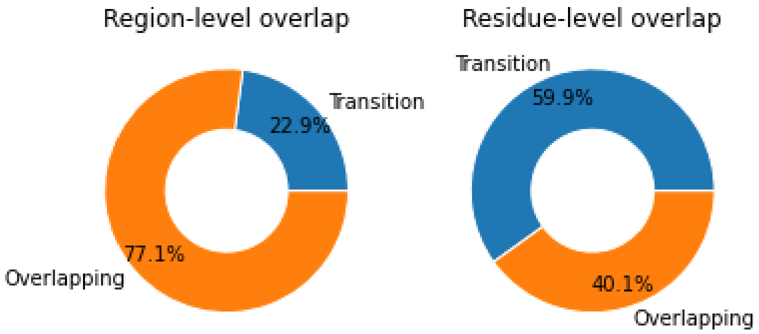


**Supplementary Figure 2** Overlap of DisProt structural transitions with PDB. Fraction of Regions/Residues labelled with a structural transition term from the Disorder Ontology that do (orange) or do not (blue) overlap with PDB resolved regions/residues.

#### Redundancy

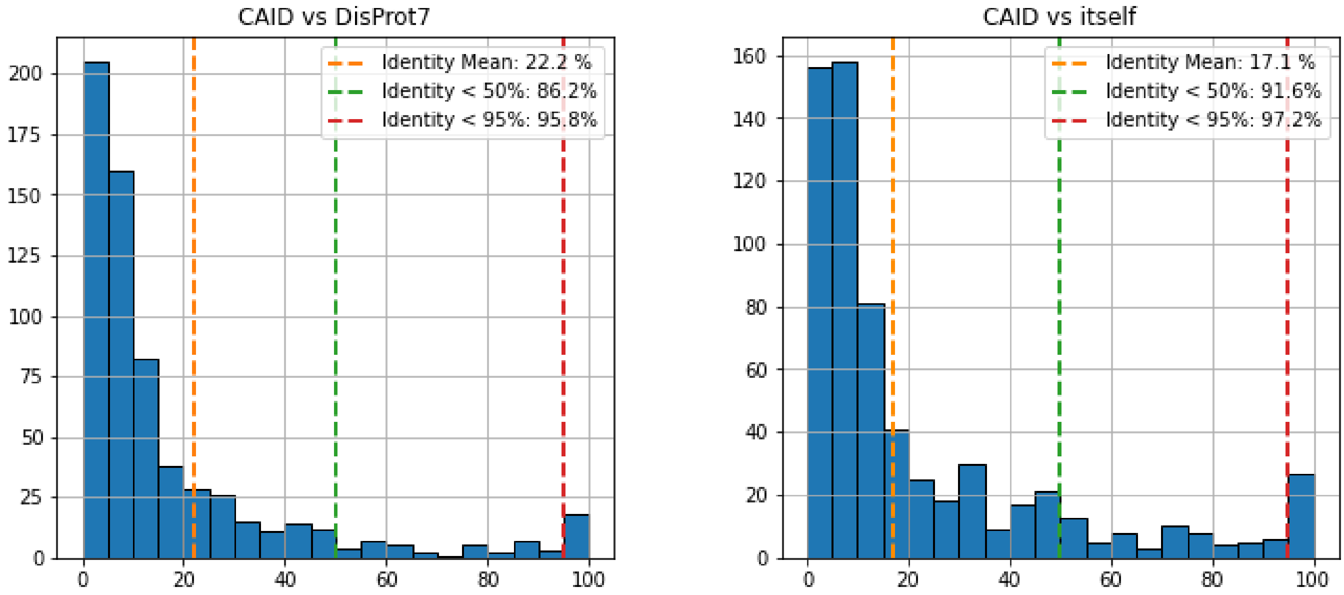


**Supplementary Figure 3** Distribution of highest sequence identity percentage for each target in the CAID dataset when compared with itself (right) and with DisProt 7.0 (left).

#### Predictor CPU time

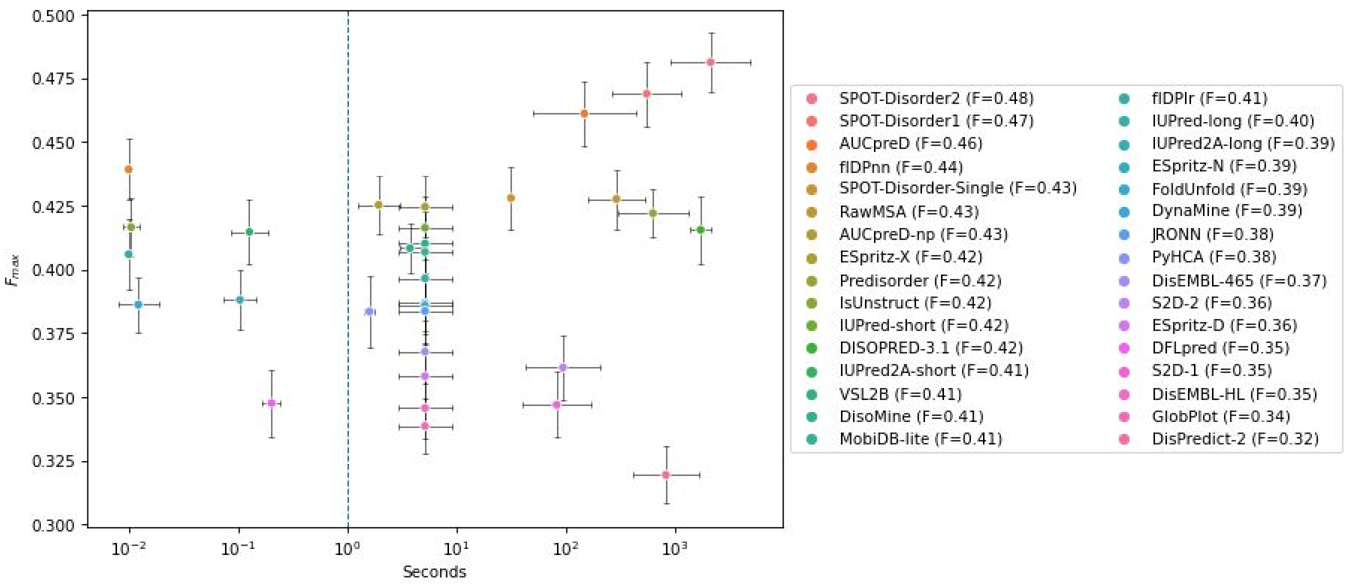


**Supplementary Figure 4**. CPU time to performance for disorder predictors. Scatterplot of CPU time in ms in logarithmic scale (x axis) and performance expressed as $F_{Max}$ (y axis) calculated on the *DisProt* dataset.

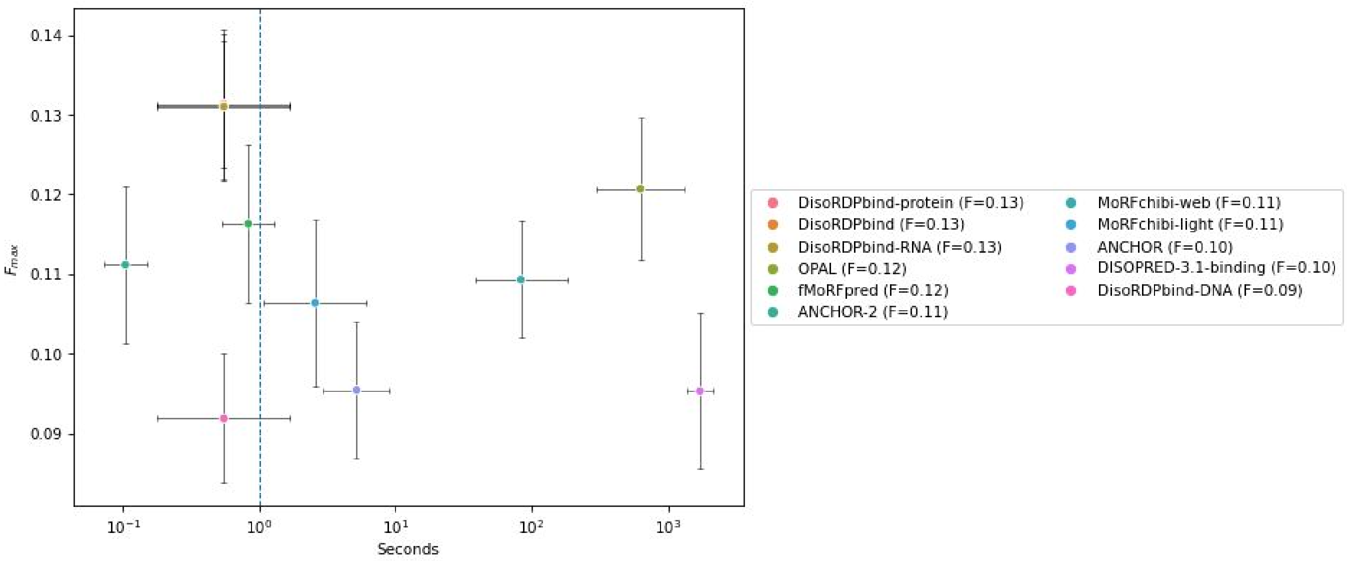


**Supplementary Figure 5**. CPU time to performance for binding predictors. Scatterplot of CPU time in ms in logarithmic scale (x axis) and performance expressed as $F_{Max}$ (yaxis) calculated on the *DisProt-Binding* dataset.

#### Species representation

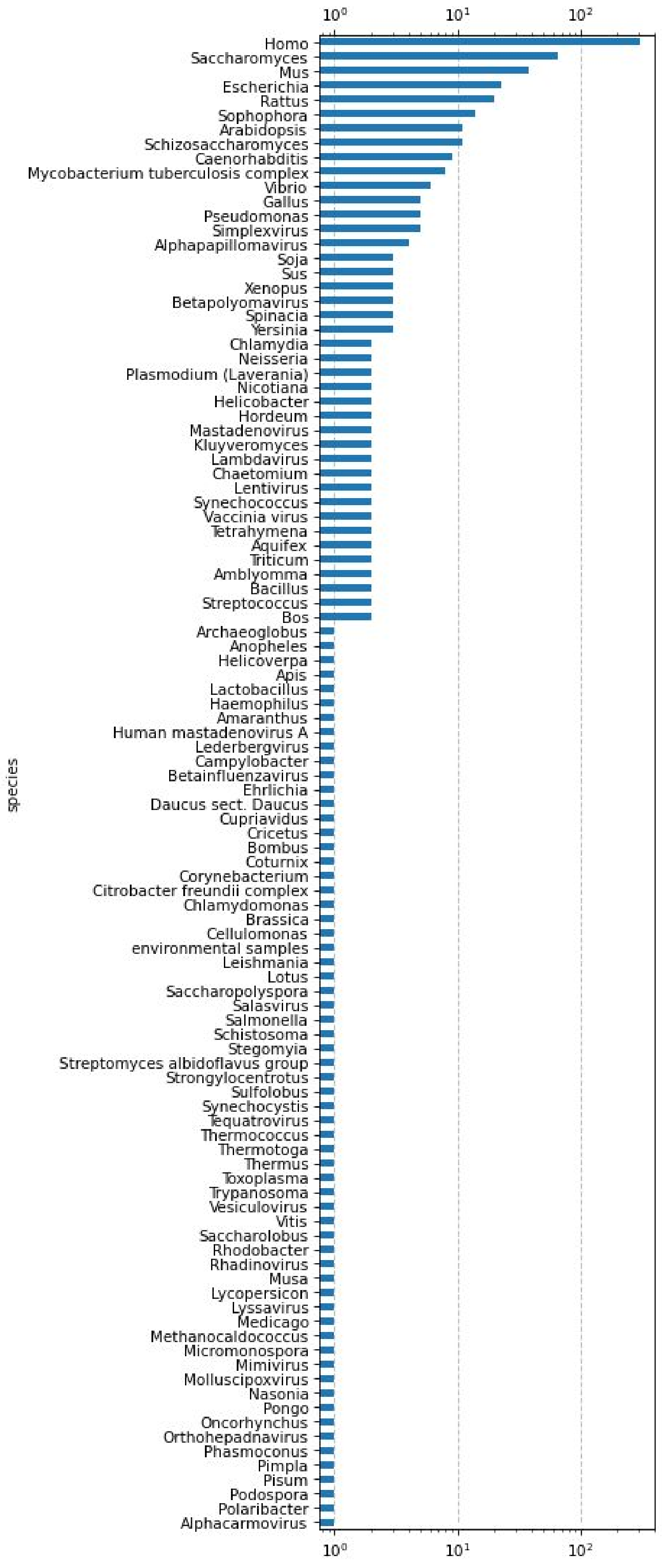


**Supplementary Figure 6**. Number of entries for each species in the CAID dataset.

#### Fully ID

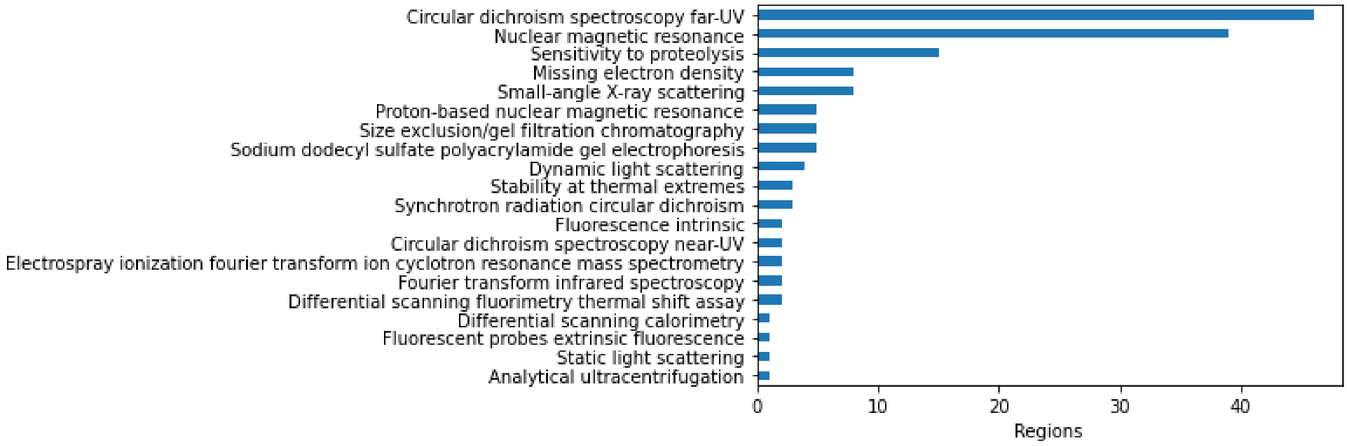


**Supplementary Figure 7.** Number of regions detected per detection method in fully disordered proteins (calculated as those proteins with a fraction of disorder greater than 95%).

### Disorder

#### DisProt dataset

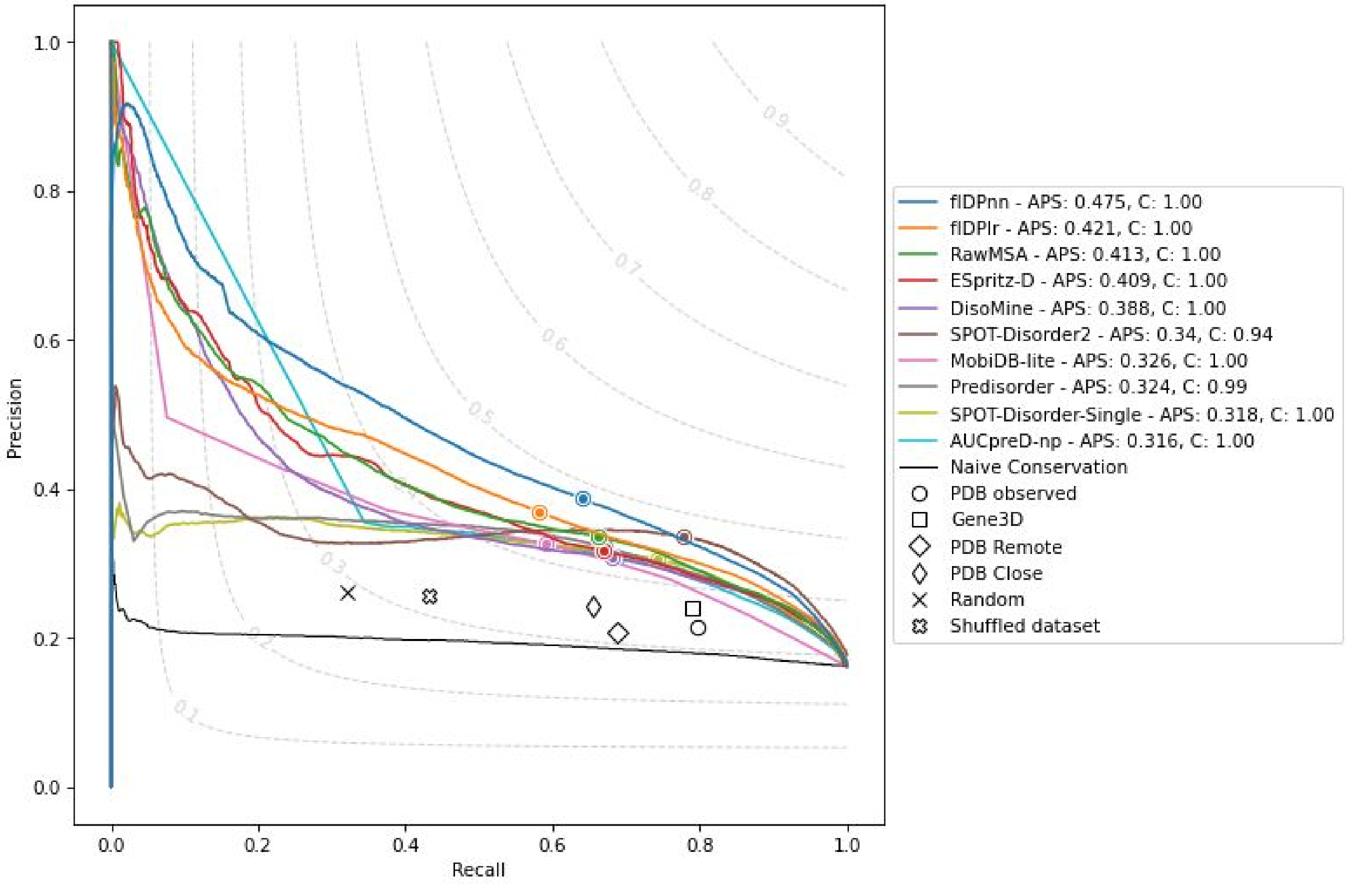


**Supplementary Figure 8**: Precision (y-axis) recall (x-axis) curves of the 10 best ranking methods. Ranking is based on their APS (average precision score) in the *DisProt* dataset.

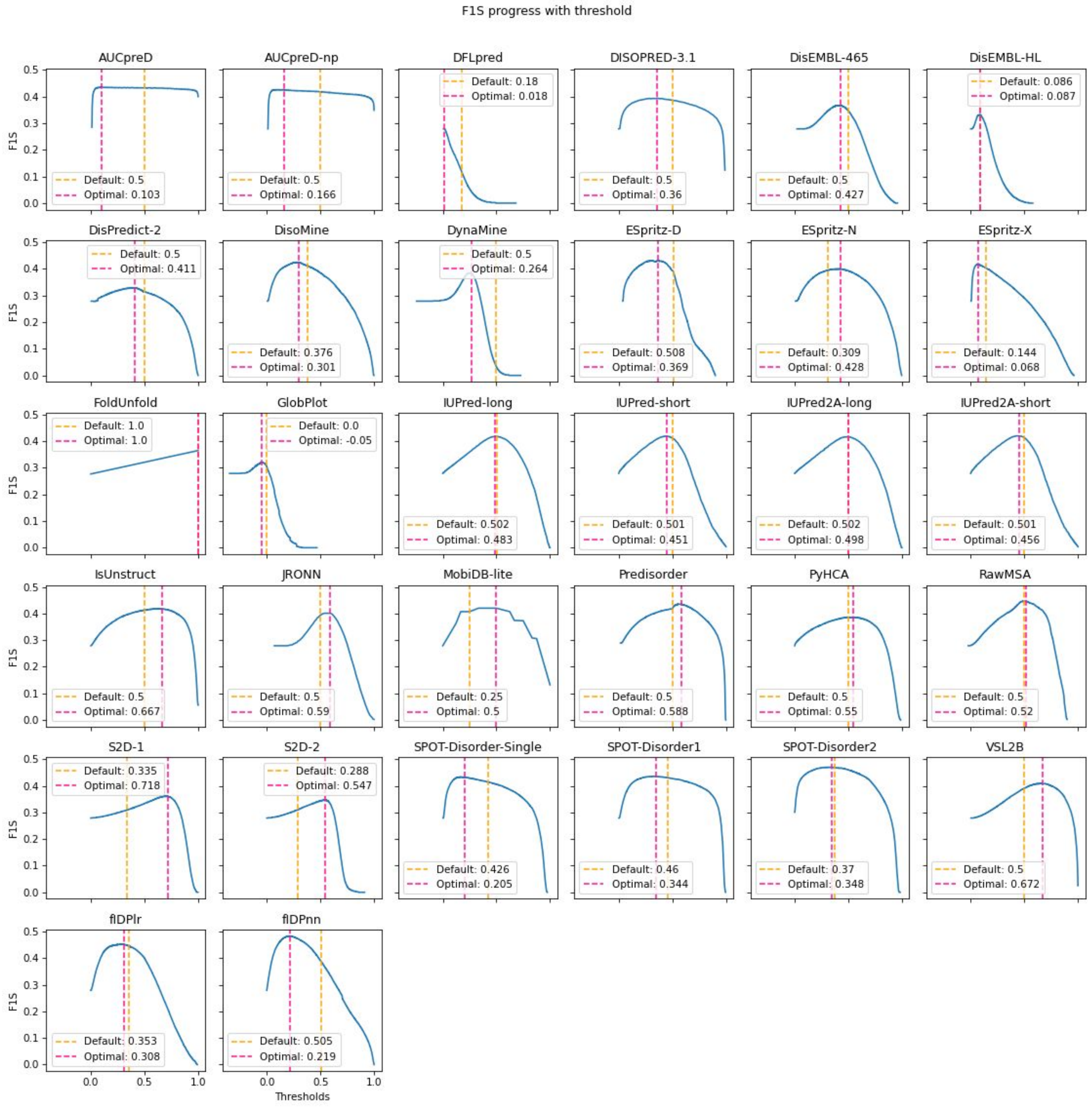


**Supplementary Figure 9**. F1-score progress (y-axis) with increasing threshold value (x-axis) for each predictor in the *DisProt* dataset.

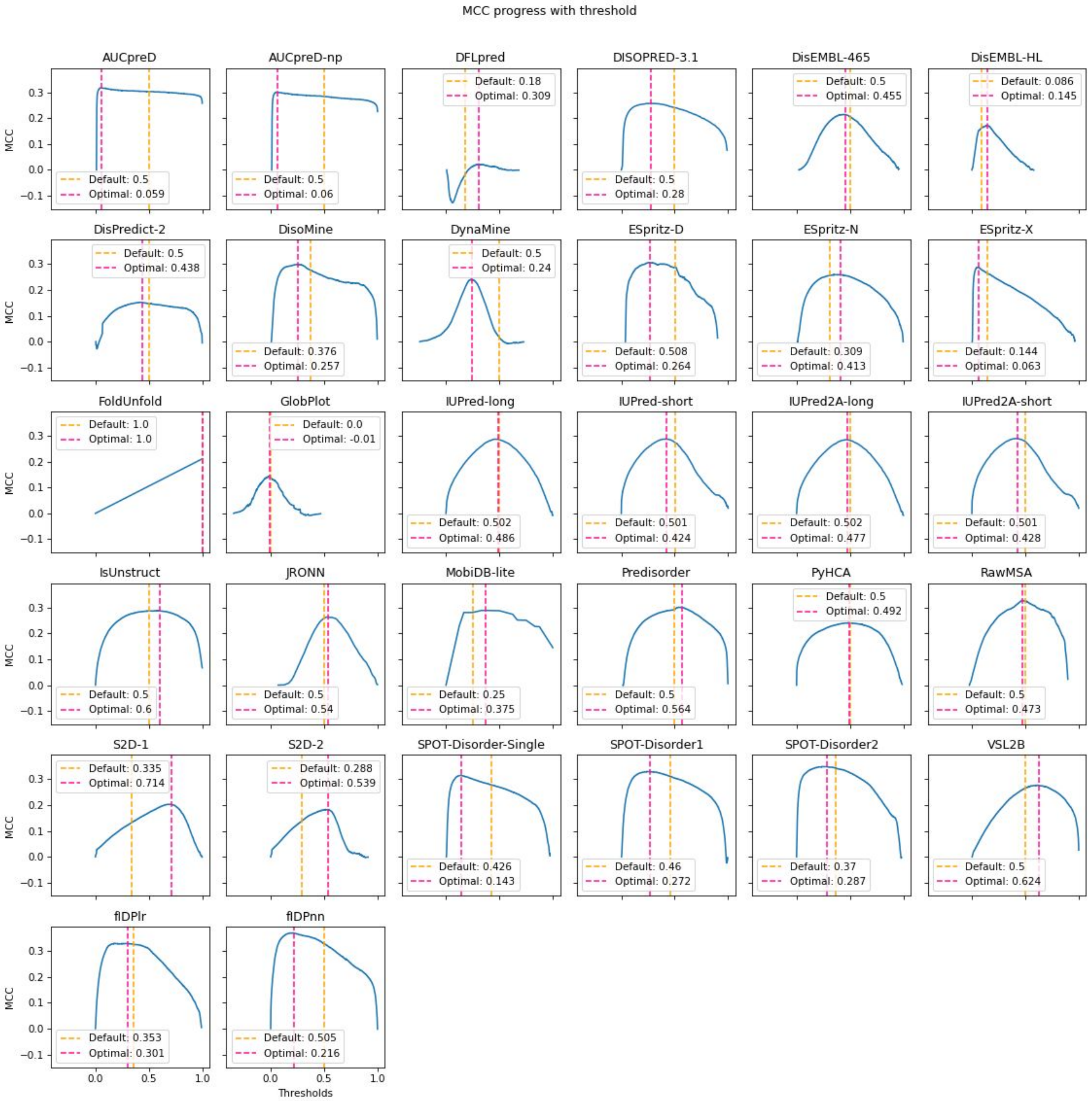


**Supplementary Figure 10**. MCC progress (y-axis) with increasing threshold value (x-axis) for each predictor in the *DisProt* dataset.

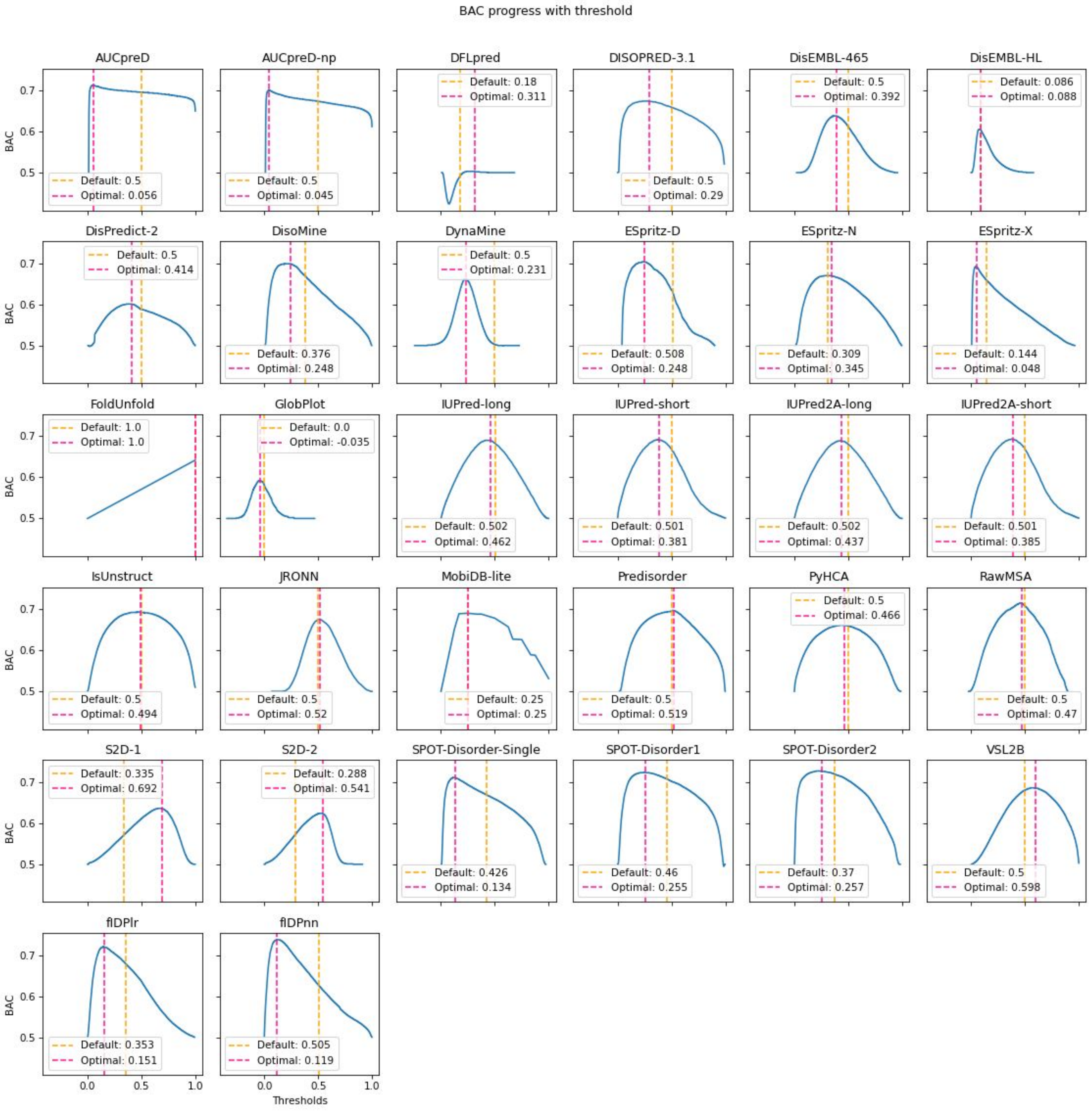


**Supplementary Figure 11**. Balanced accuracy progress (y-axis) with increasing threshold value (x-axis) for each predictor in the *DisProt* dataset.

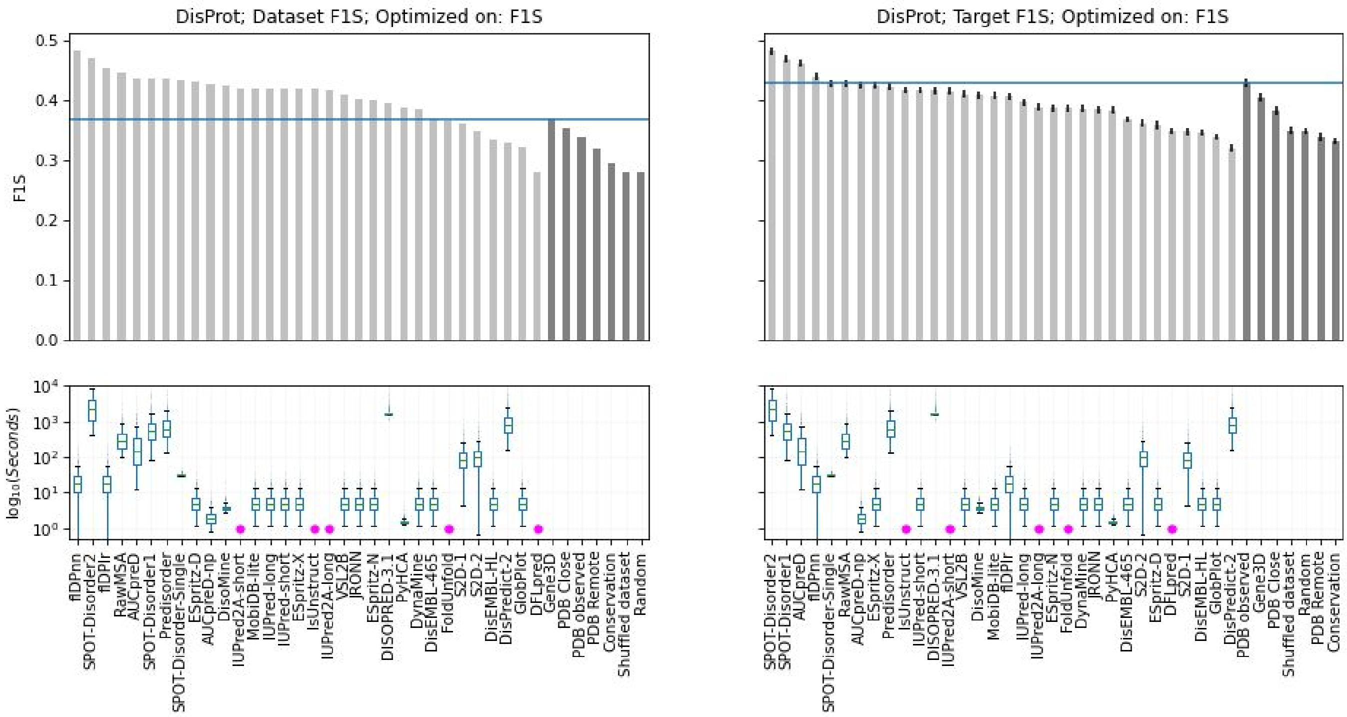


**Supplementary Figure 12**: $F_{Max}$ calculated on the whole dataset with confidence intervals as error bars (left) and averaged over proteins with Standard-Error as error bars (right). Calculated on *DisProt* dataset. Magenta dots indicate that the whole distribution of execution-times is lower than 1 second.

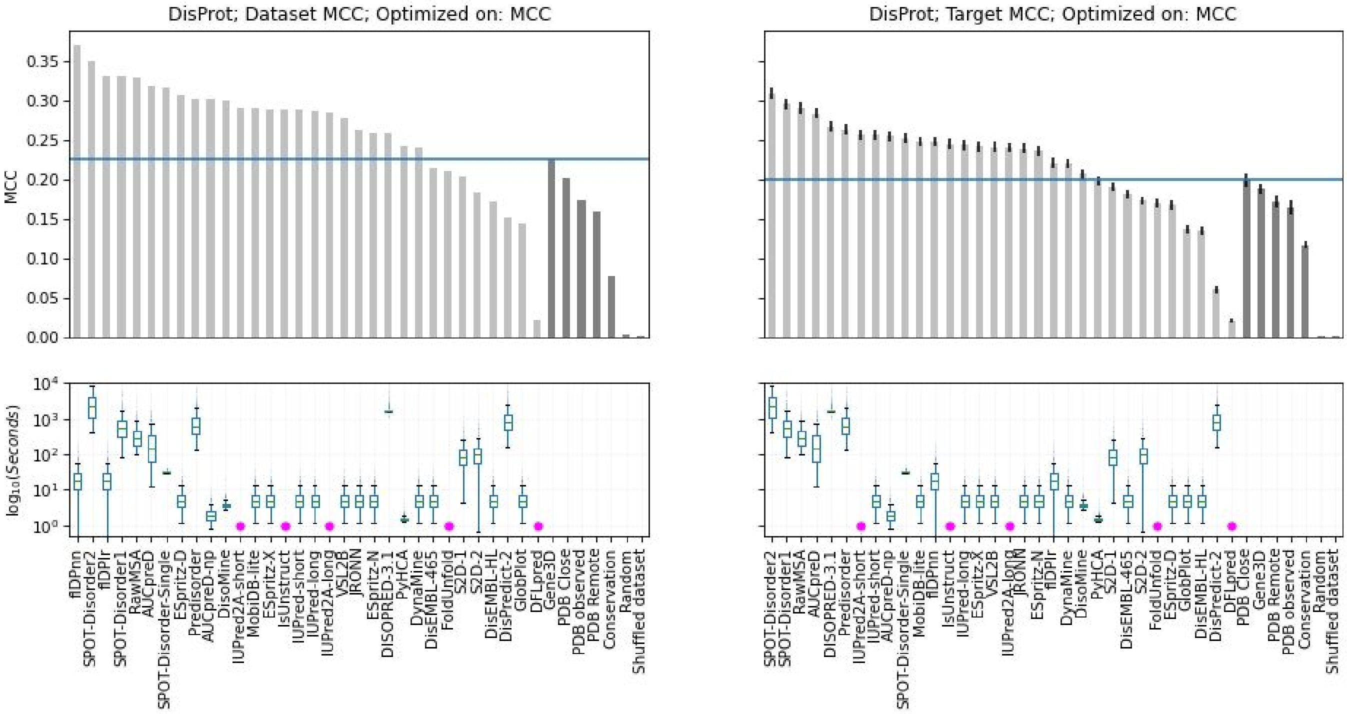


**Supplementary Figure 13**: MCC calculated on the whole dataset with confidence intervals as error bars (left) and averaged over proteins with Standard-Error as error bars (right). Calculated on *DisProt* dataset. Predictors threshold is optimized on MCC. Magenta dots indicate that the whole distribution of execution-times is lower than 1 second.

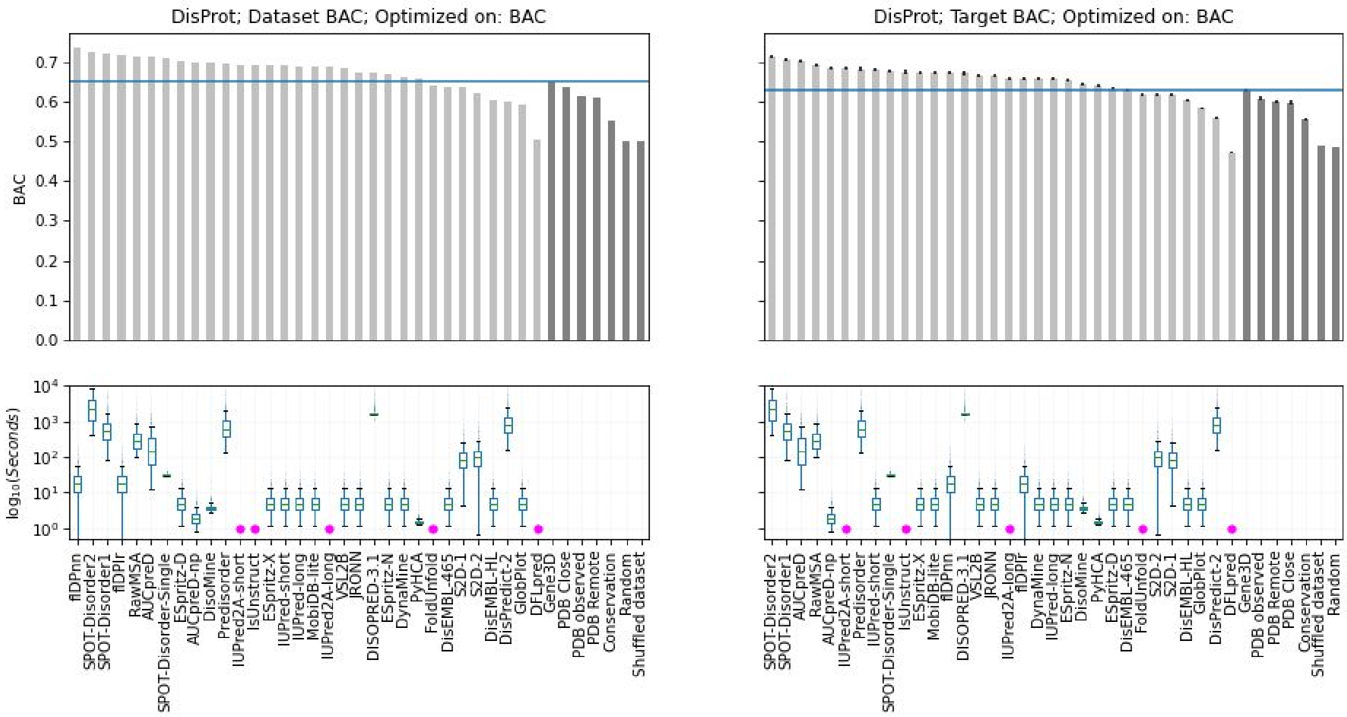


**Supplementary Figure 14**: Balanced accuracy calculated on the whole dataset with confidence intervals as error bars (left) and averaged over proteins with Standard-Error as error bar (right). Calculated *DisProt* dataset. Predictors threshold is optimized on Balanced accuracy. Magenta dots indicate that the whole distribution of execution-times is lower than 1 second.

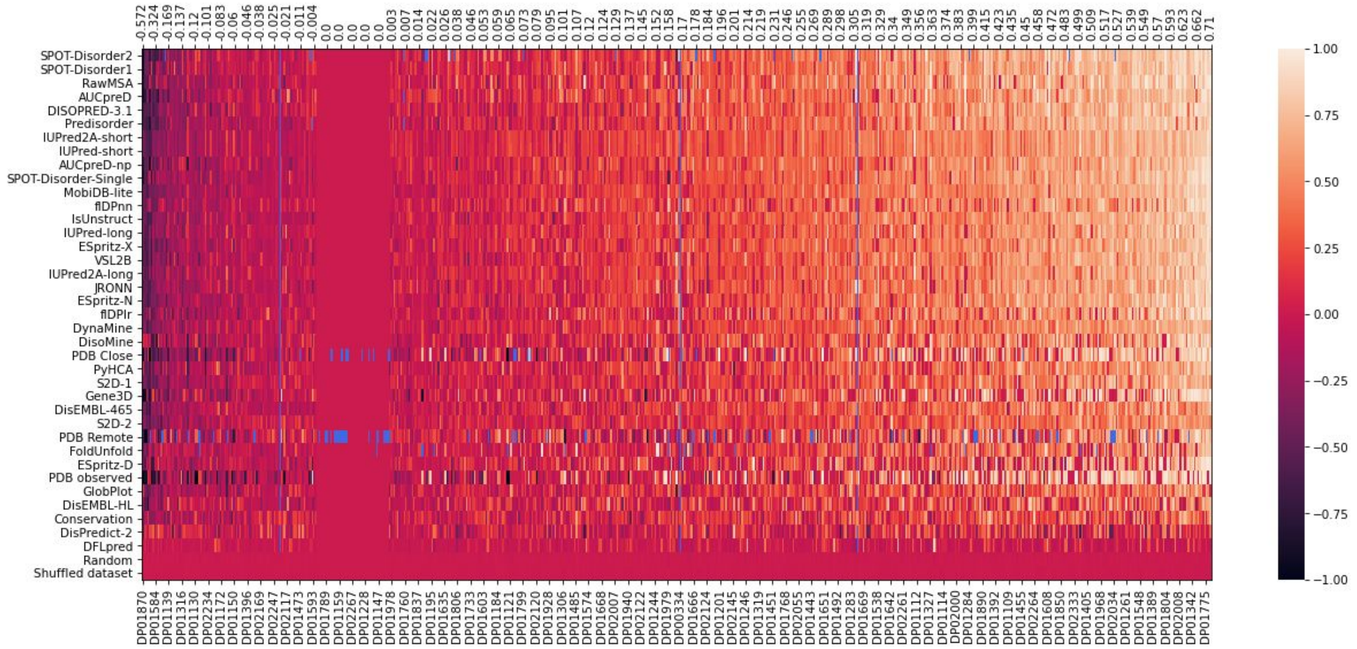


**Supplementary Figure 15**: MCC of each target (x-axis, bottom labels, not all labels are visible) from each predictor (y-axis). Targets are sorted by average MCC (x-axis, top labels). Calculated on *DisProt* dataset. Predictors threshold is optimized on MCC. Missing values are in blue.

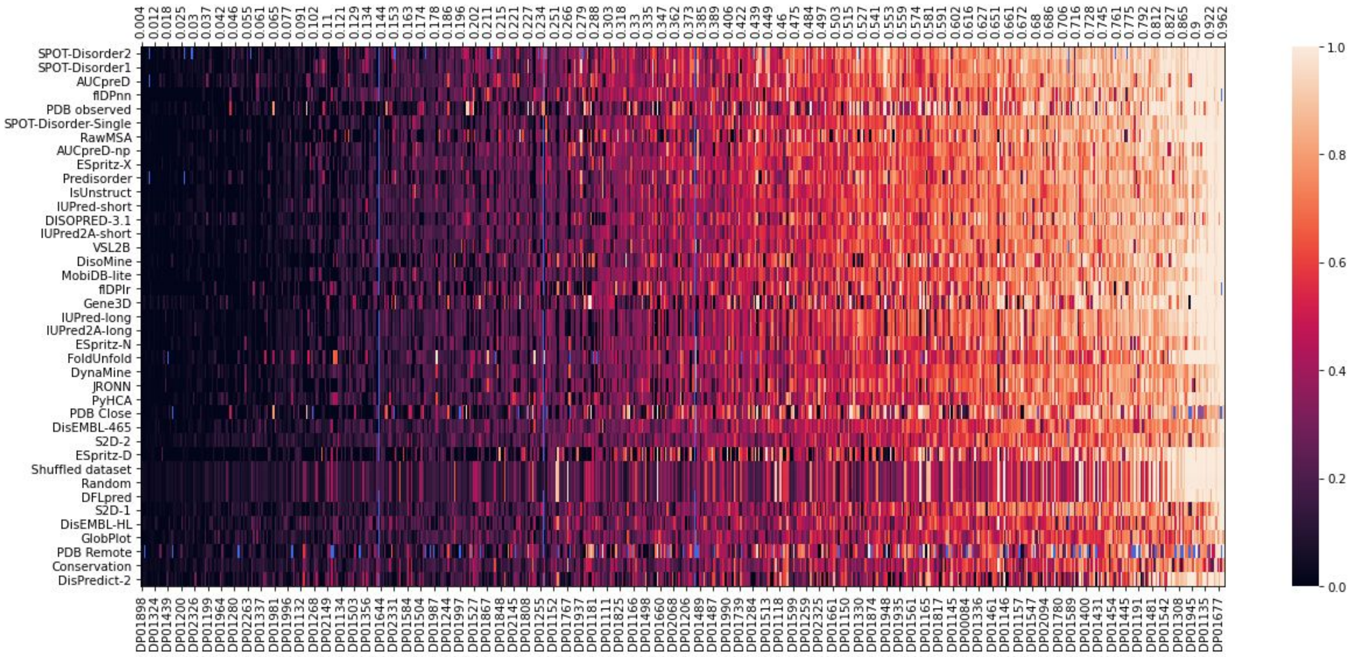


**Supplementary Figure 16**: $F_{Max}$ of each target (x-axis, bottom labels, not all labels are visible) from each predictor (y-axis). Targets are sorted by average $F_{Max}$ (x-axis, top labels). Calculated on all *DisProt* dataset. Missing values are in blue.

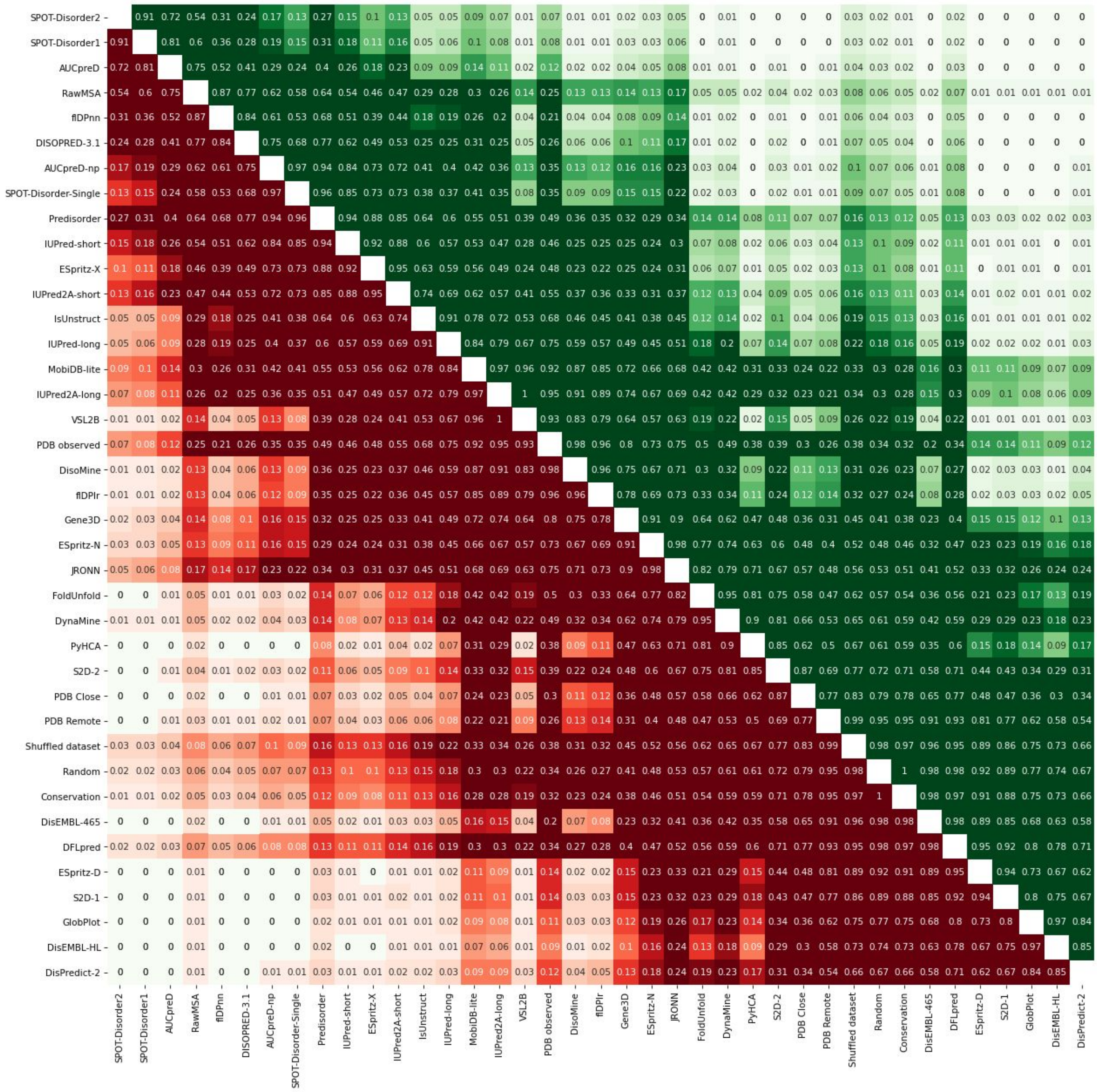

**Supplementary Figure 17**. Overall average ranking of all predictors and baselines in the *DisProt* dataset. Heatmap of the T-test p-value associated to the statistical significance of the difference between ranking distribution of predictors. A ranking distribution for a predictor is the position of that predictor in its ranking for the following metrics: 'bac', 'f1s', 'fpr', 'mcc', 'ppv', 'tpr', 'tnr'. Metrics used are: bac, f1s, fpr, mcc, ppv, tpr, tnr; they are calculated per target and with predictors threshold optimized by F1-Score.

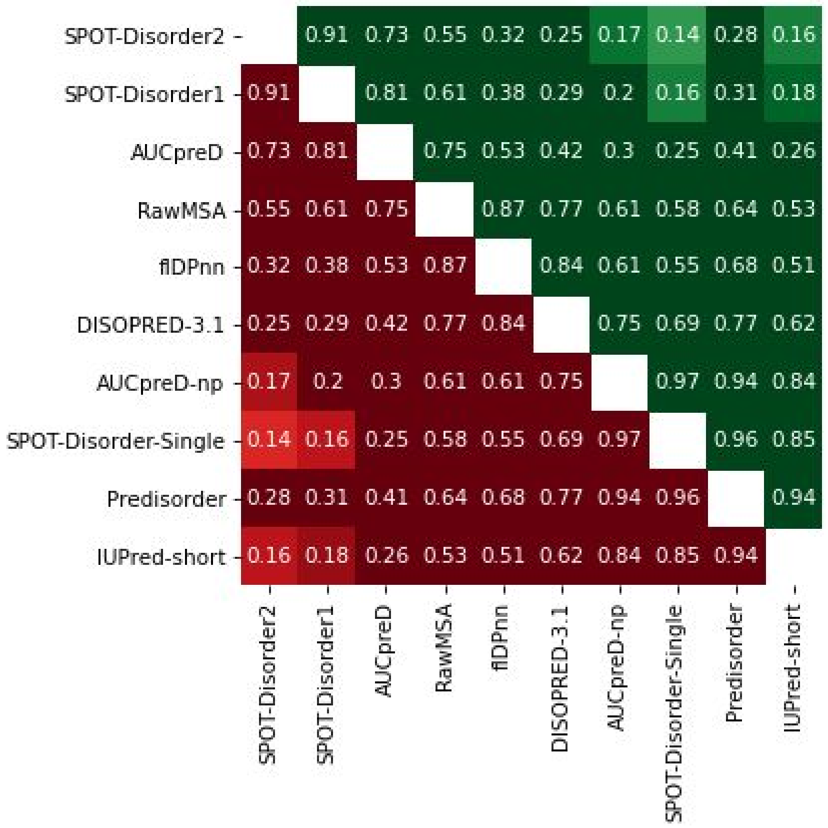


**Supplementary Figure 18**. Overall average ranking of the 10 best ranking predictors and baselines in the *DisProt* dataset. Heatmap of the T-test p-value associated to the statistical significance of the difference between ranking distribution of predictors. A ranking distribution for a predictor is the position of that predictor in its ranking for each metric. Metrics used are: bac, f1s, fpr, mcc, ppv, tpr, tnr; they are calculated with predictors threshold optimized by F1-Score.

Mammals

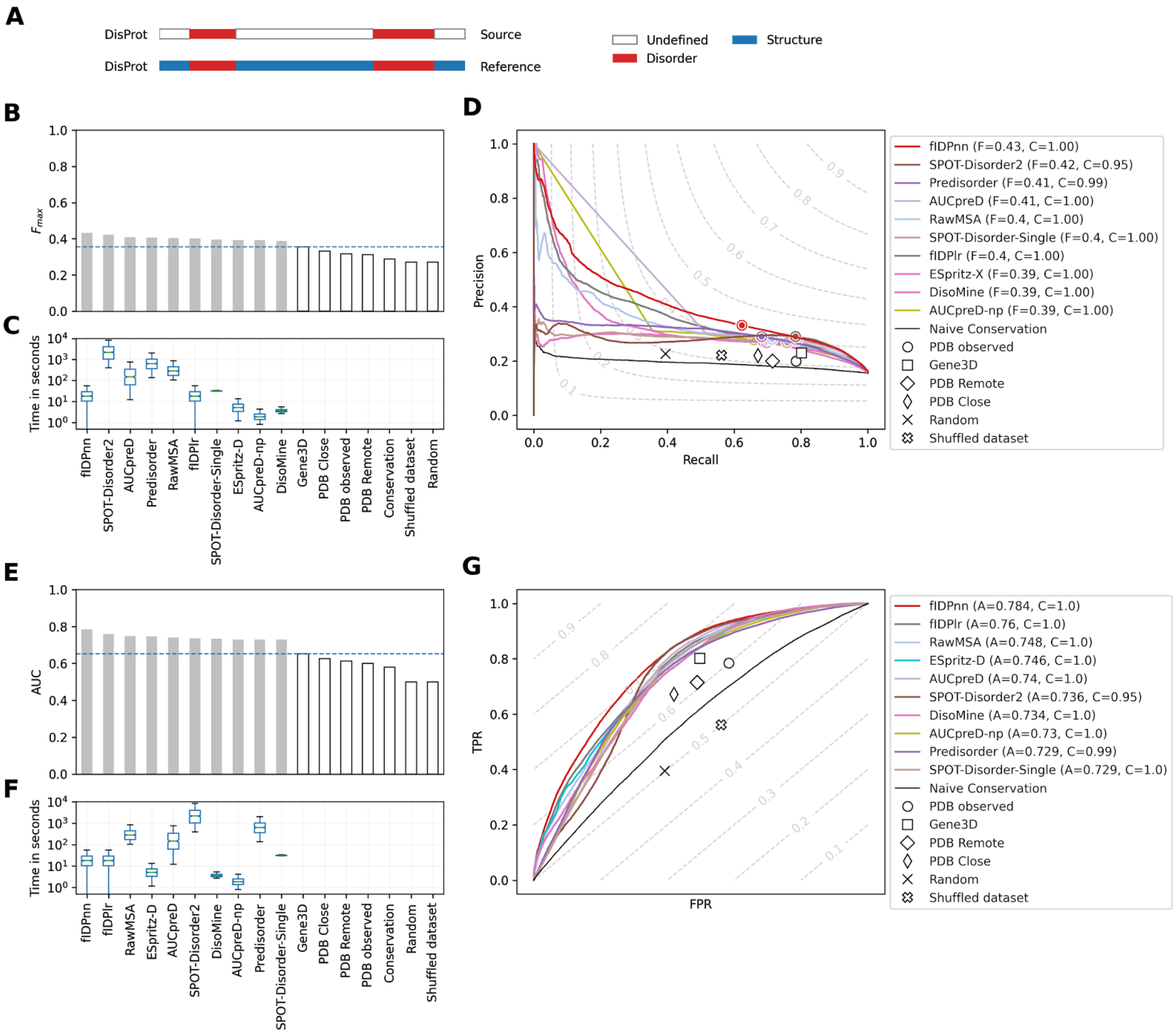


**Supplementary Figure 19.** Prediction success and CPU times for the ten top-ranking disorder predictors for mammalian proteins in the *DisProt* dataset. Reference used (*DisProt*) in the analysis and how it is obtained (panel A). Performance of predictors expressed as maximum F1-Score across all thresholds ($F_{max}$) (panel B) and AUC (panel E) for the top ten best ranking methods (light gray) and baselines (white) and the distribution of execution time per-target (panels C, F) using *DisProt* dataset. The horizontal line in panels B, E indicates the $F_{max}$ and AUC of the best baseline, respectively. Precision-Recall (panel D) and ROC curves (panel G) of ten top-ranking methods and baselines using *DisProt* dataset, with level curves of the F1-Score and Balanced accuracy, respectively. Magenta dots on panels C, F indicate that the whole distribution of execution-times is lower than 1 second.

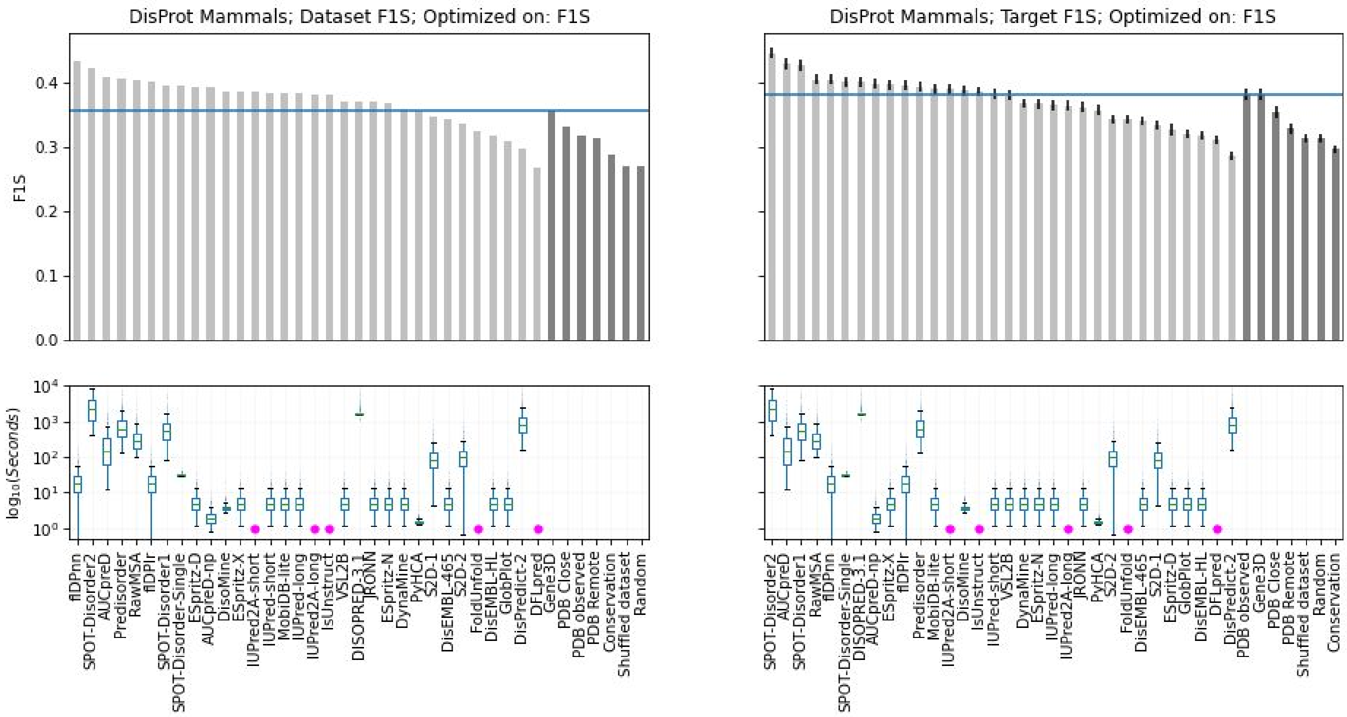


**Supplementary Figure 20**: $F_{Max}$ calculated on the whole dataset with confidence intervals as error bars (left) and averaged over proteins with Standard-Error as error bars (right). Calculated on mammalian proteins of the *DisProt* dataset.  Magenta dots indicate that the whole distribution of execution-times is lower than 1 second.

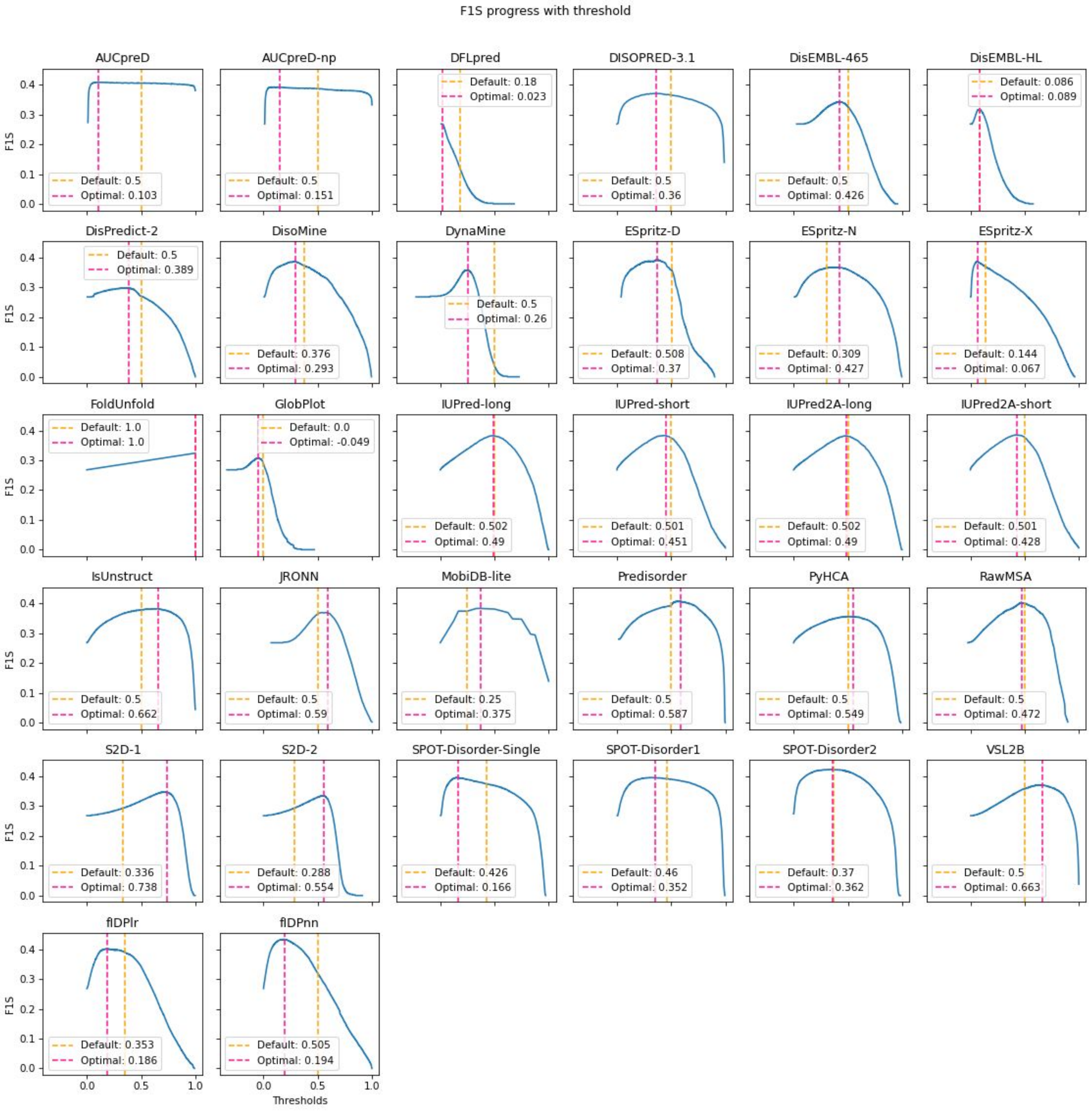


**Supplementary Figure 21**. F1-score progress (y-axis) with increasing threshold value (x-axis) for each predictor calculated on mammalian proteins on the *DisProt* dataset.

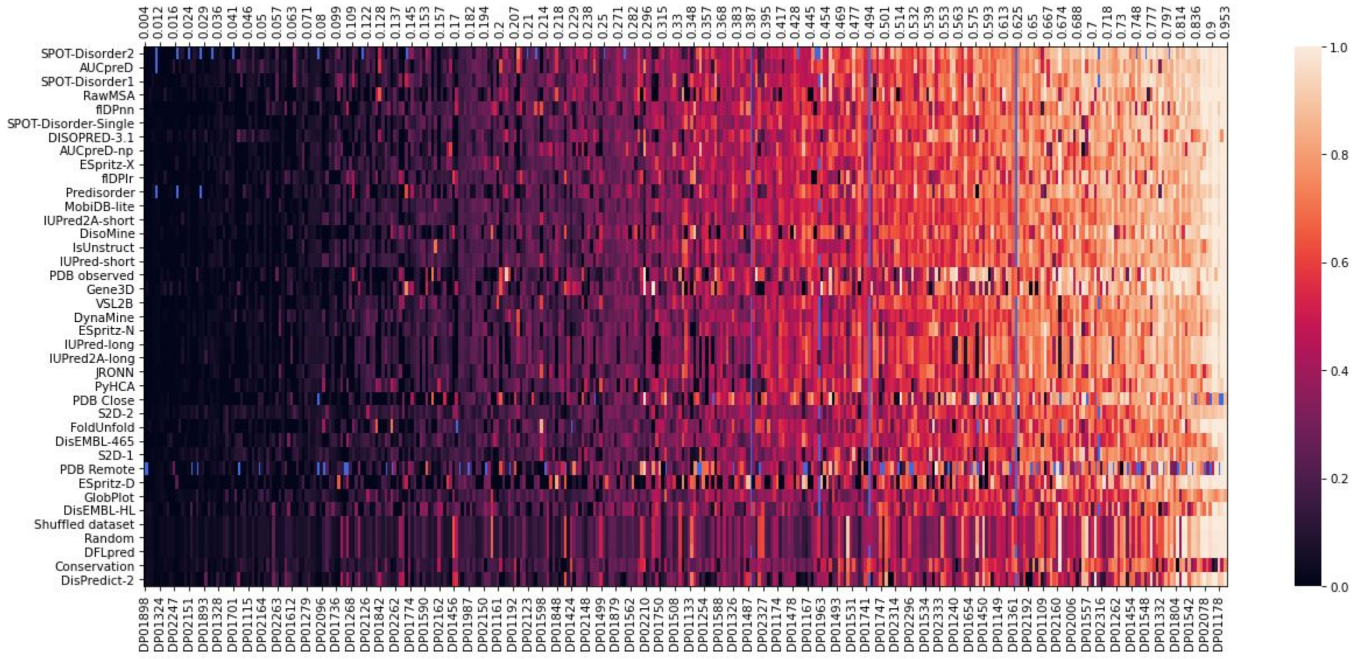


**Supplementary Figure 22**: $F_{Max}$ of each target (x-axis, bottom labels, not all labels are visible) from each predictor (y-axis). Targets are sorted by average $F_{Max}$ (x-axis, top labels). Calculated on mammalian proteins of the *DisProt* dataset. Missing values are in blue.

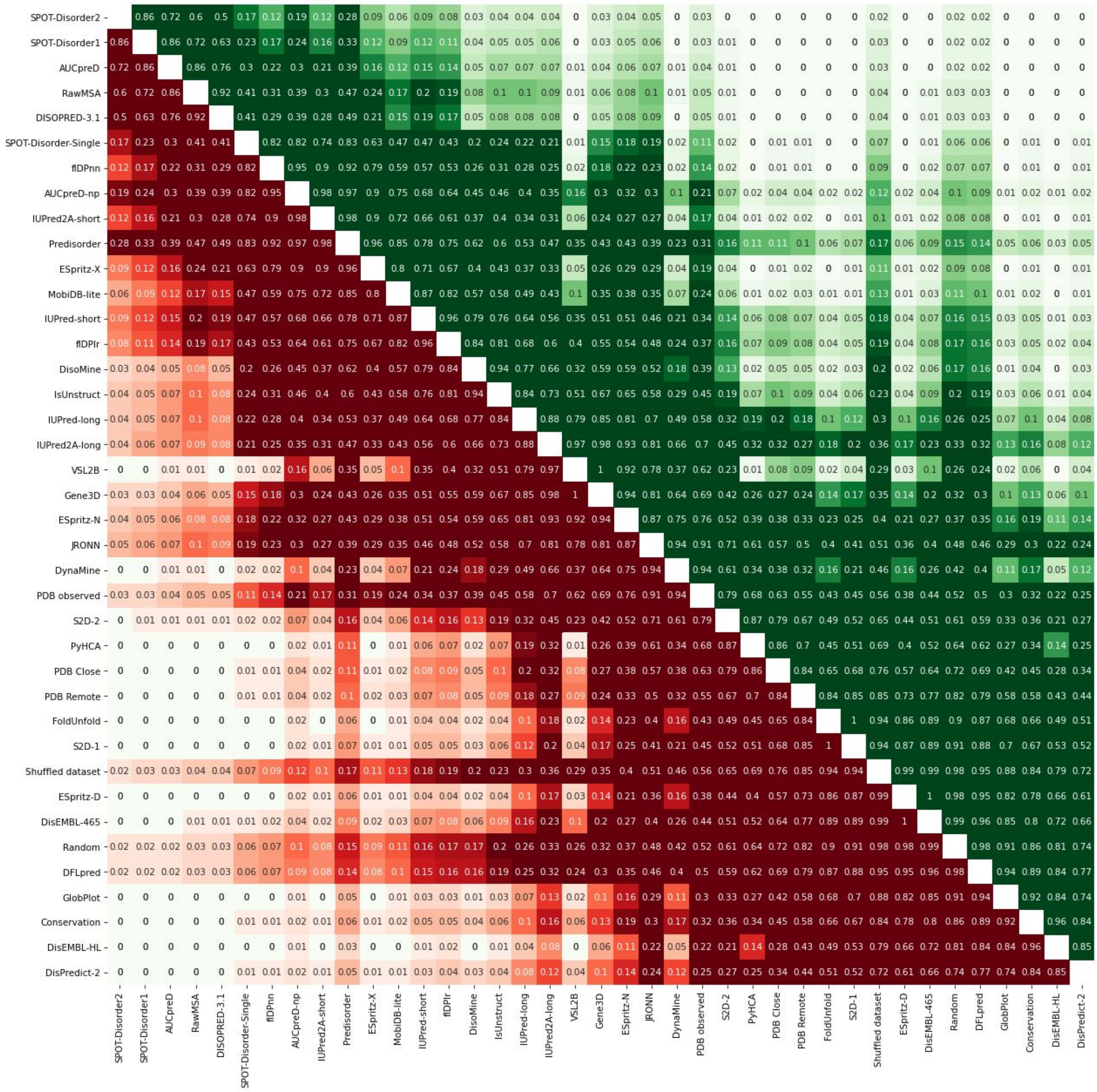

**Supplementary Figure 23**. Overall average ranking of all predictors and baselines for mammalian proteins in the *DisProt* dataset. Heatmap of the T-test p-value associated to the statistical significance of the difference between ranking distribution of predictors. A ranking distribution for a predictor is the position of that predictor in its ranking for the following metrics: 'bac', 'f1s', 'fpr', 'mcc', 'ppv', 'tpr', 'tnr'. Metrics are calculated per target and with predictors threshold optimized by F1-Score.

#### Prokaryotes

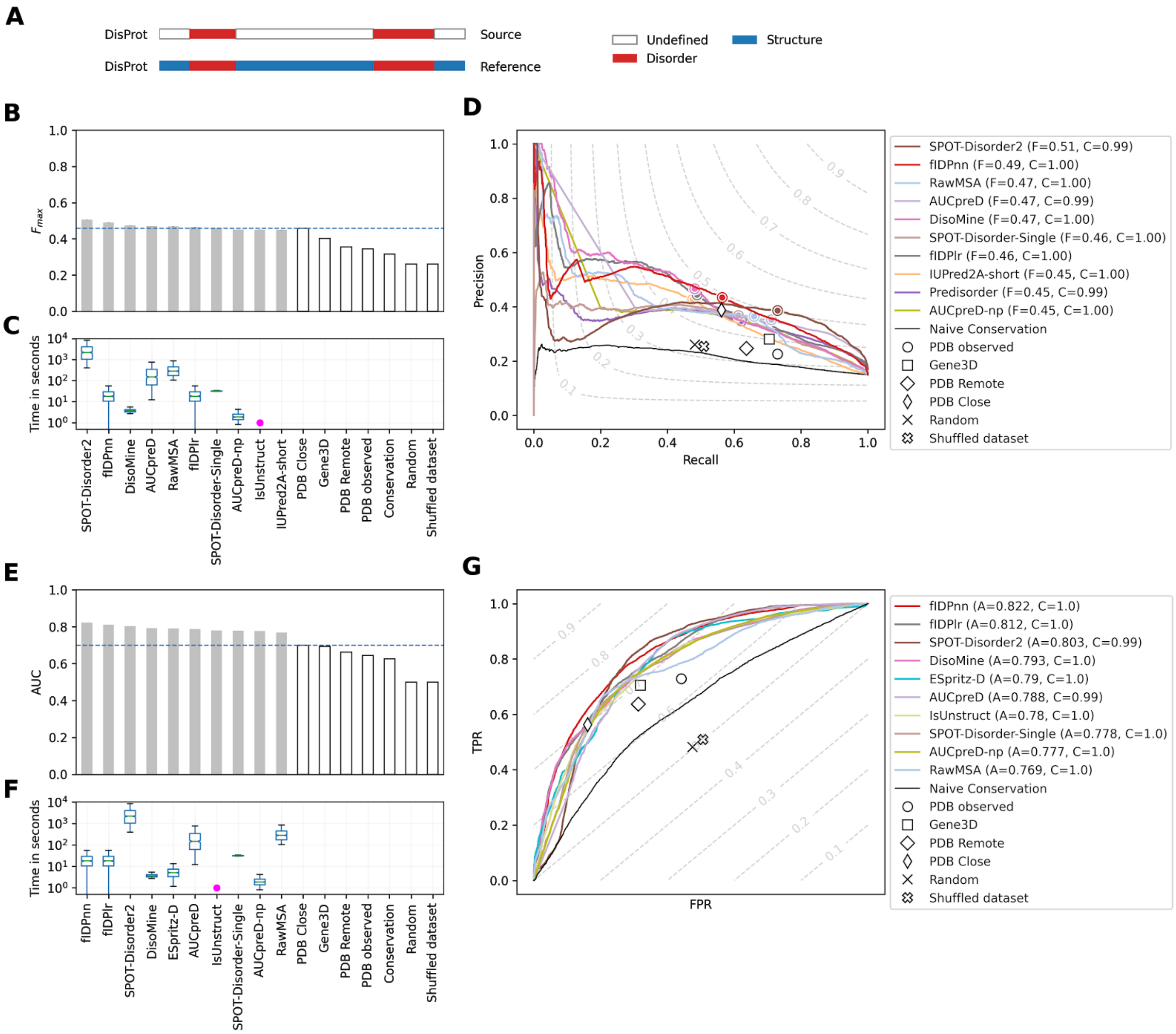


**Supplementary Figure 24.** Prediction success and CPU times for the ten top-ranking disorder predictors for prokaryotic proteins in the *DisProt* dataset. Reference used (*DisProt*) in the analysis and how it is obtained (panel A). Performance of predictors expressed as maximum F1-Score across all thresholds ($F_{max}$) (panel B) and AUC (panel E) for the top ten best ranking methods (light gray) and baselines (white) and the distribution of execution time per-target (panels C, F) using *DisProt* dataset. The horizontal line in panels B, E indicates the $F_{max}$ and AUC of the best baseline, respectively. Precision-Recall (panel D) and ROC curves (panel G) of ten top-ranking methods and baselines using *DisProt* dataset, with level curves of the F1-Score and Balanced accuracy, respectively. Magenta dots on panels C, F indicate that the whole distribution of execution-times is lower than 1 second.

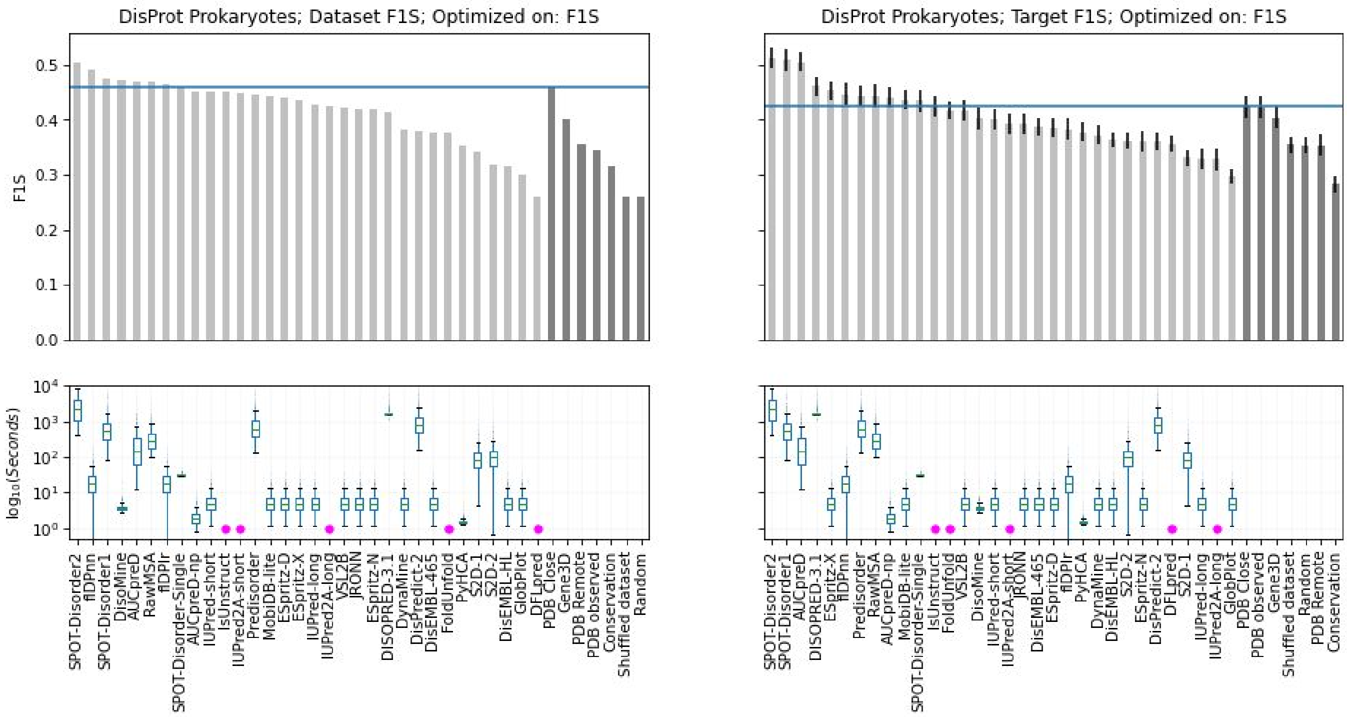


**Supplementary Figure 25**: $F_{Max}$ calculated on the whole dataset with confidence intervals as error bars (left) and averaged over proteins with Standard-Error as error bars (right). Calculated on prokaryotic proteins of the *DisProt* dataset. Magenta dots indicate that the whole distribution of execution-times is lower than 1 second.

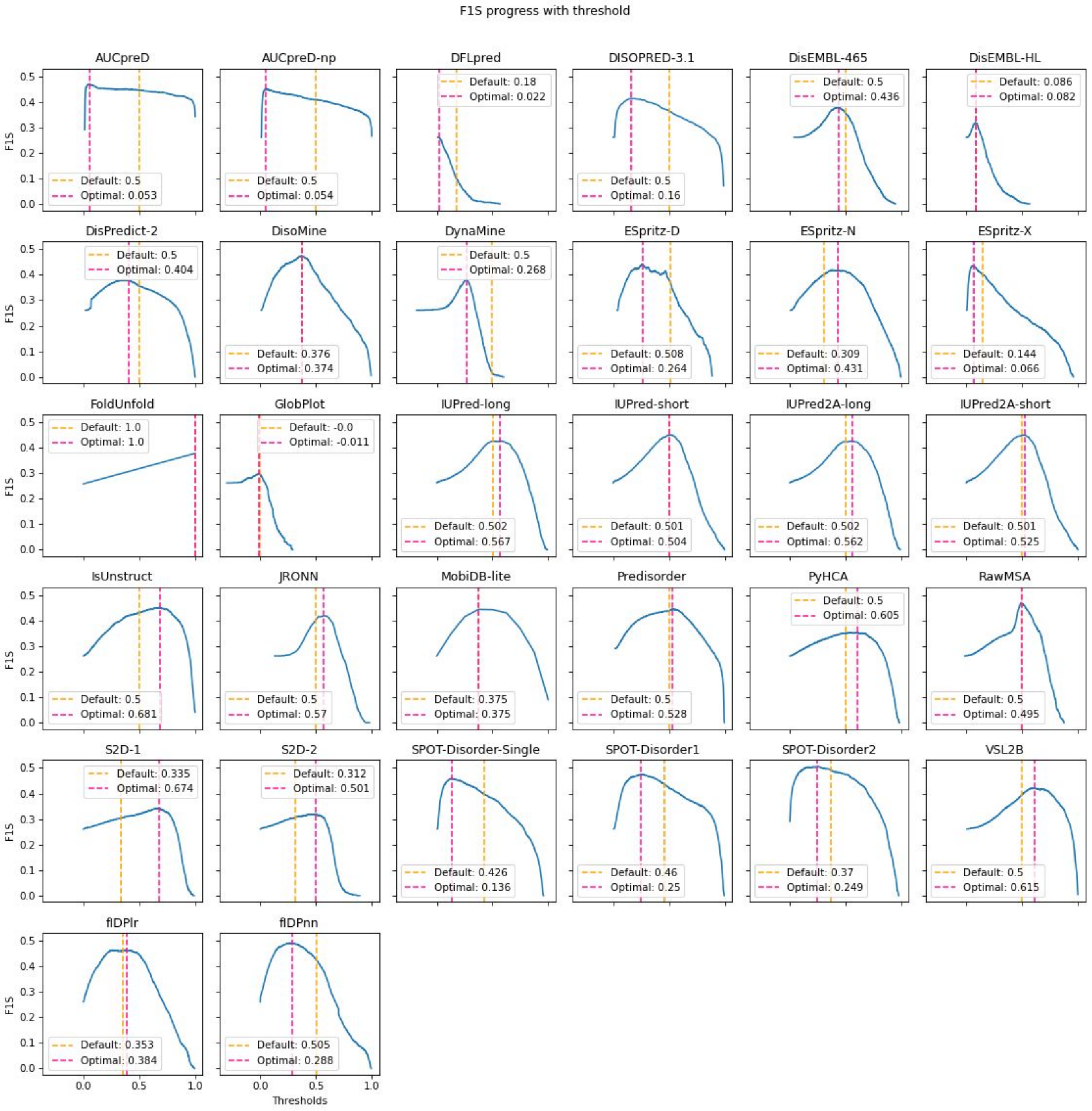


**Supplementary Figure 26**. F1-score progress (y-axis) with increasing threshold value (x-axis) for each predictor calculated on prokaryotic proteins on the *DisProt* dataset.

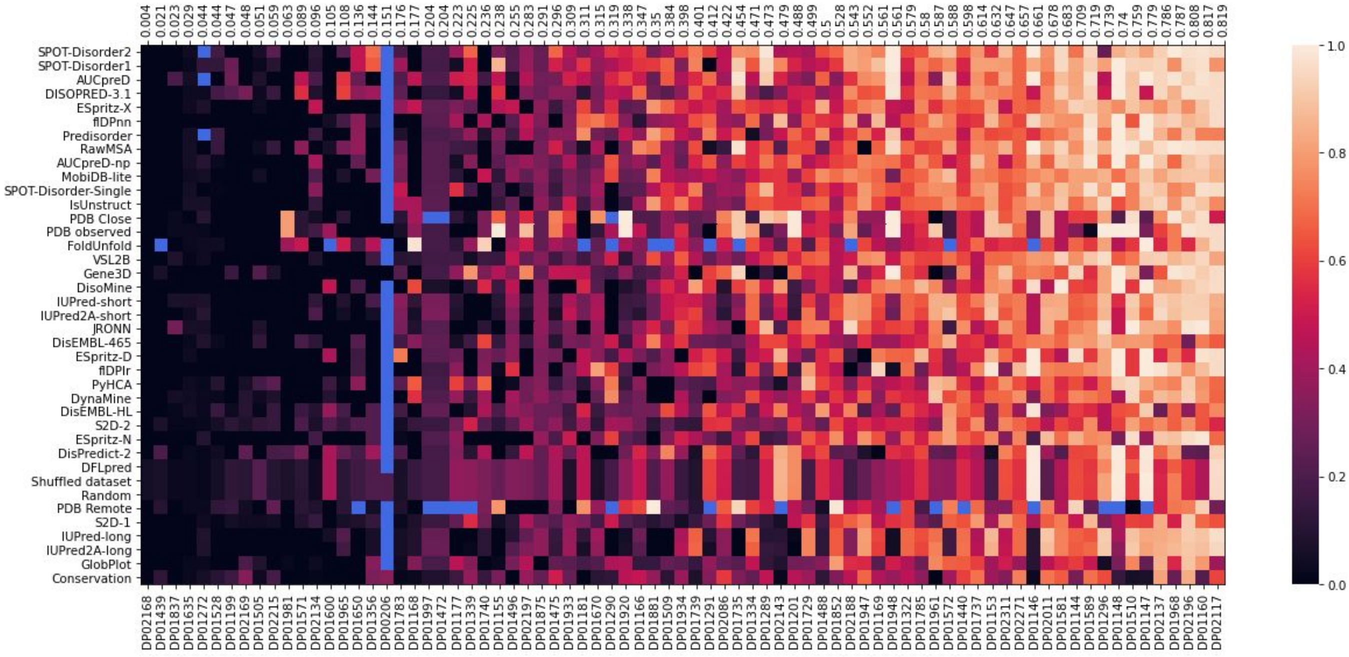


**Supplementary Figure 27**: $F_{Max}$ of each target (x-axis, bottom labels) from each predictor (y-axis). Targets are sorted by average $F_{Max}$ (x-axis, top labels). Calculated on prokaryotic proteins of the *DisProt* dataset. Missing values are in blue.

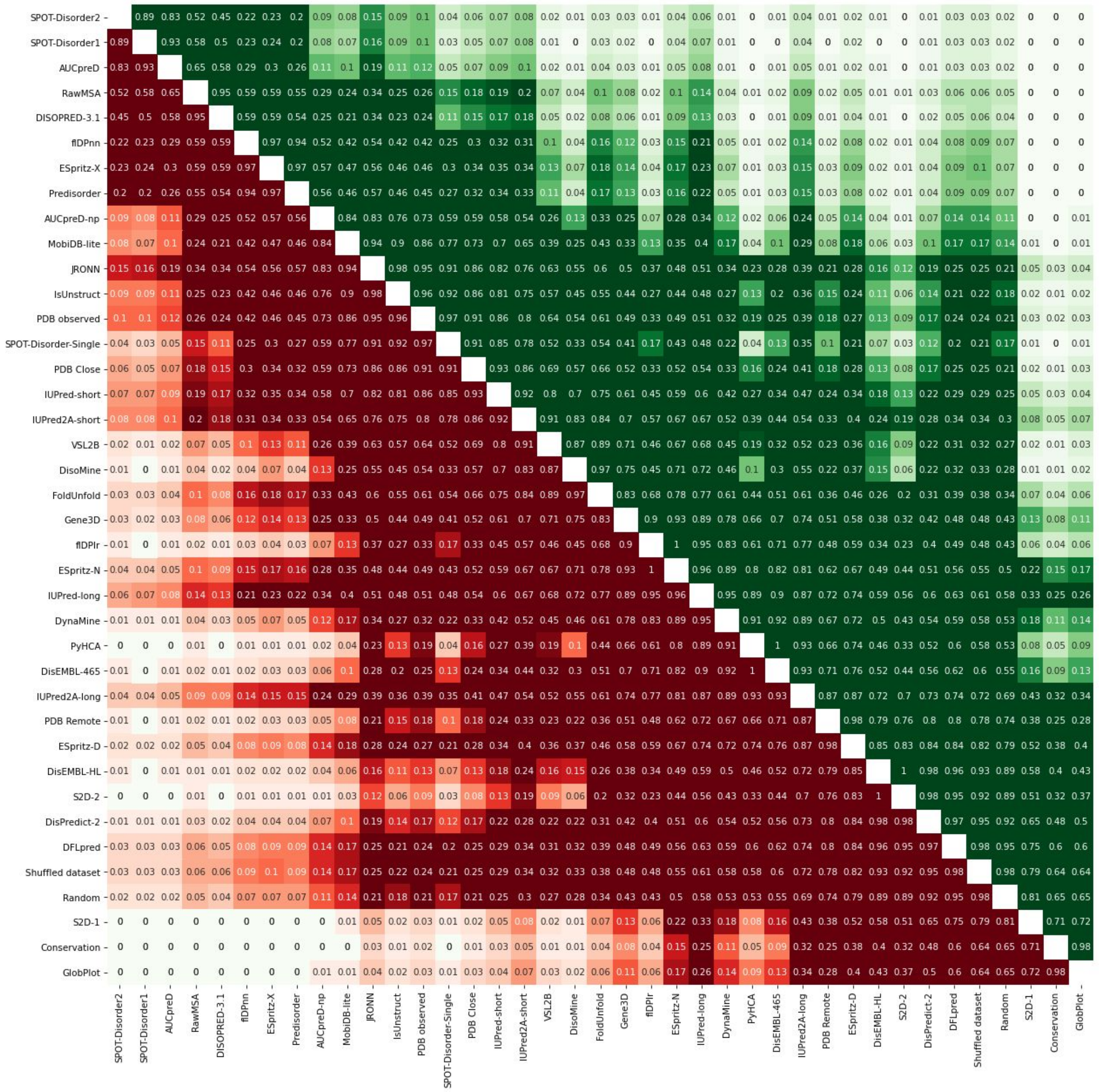

**Supplementary Figure 28**. Overall average ranking of all predictors and baselines for prokaryotic proteins in the *DisProt* dataset. Heatmap of the T-test p-value associated to the statistical significance of the difference between ranking distribution of predictors. A ranking distribution for a predictor is the position of that predictor in its ranking for the following metrics: 'bac', 'f1s', 'fpr', 'mcc', 'ppv', 'tpr', 'tnr'. Metrics are calculated per target and with predictors threshold optimized by F1-Score.

#### DisProt-PDB dataset

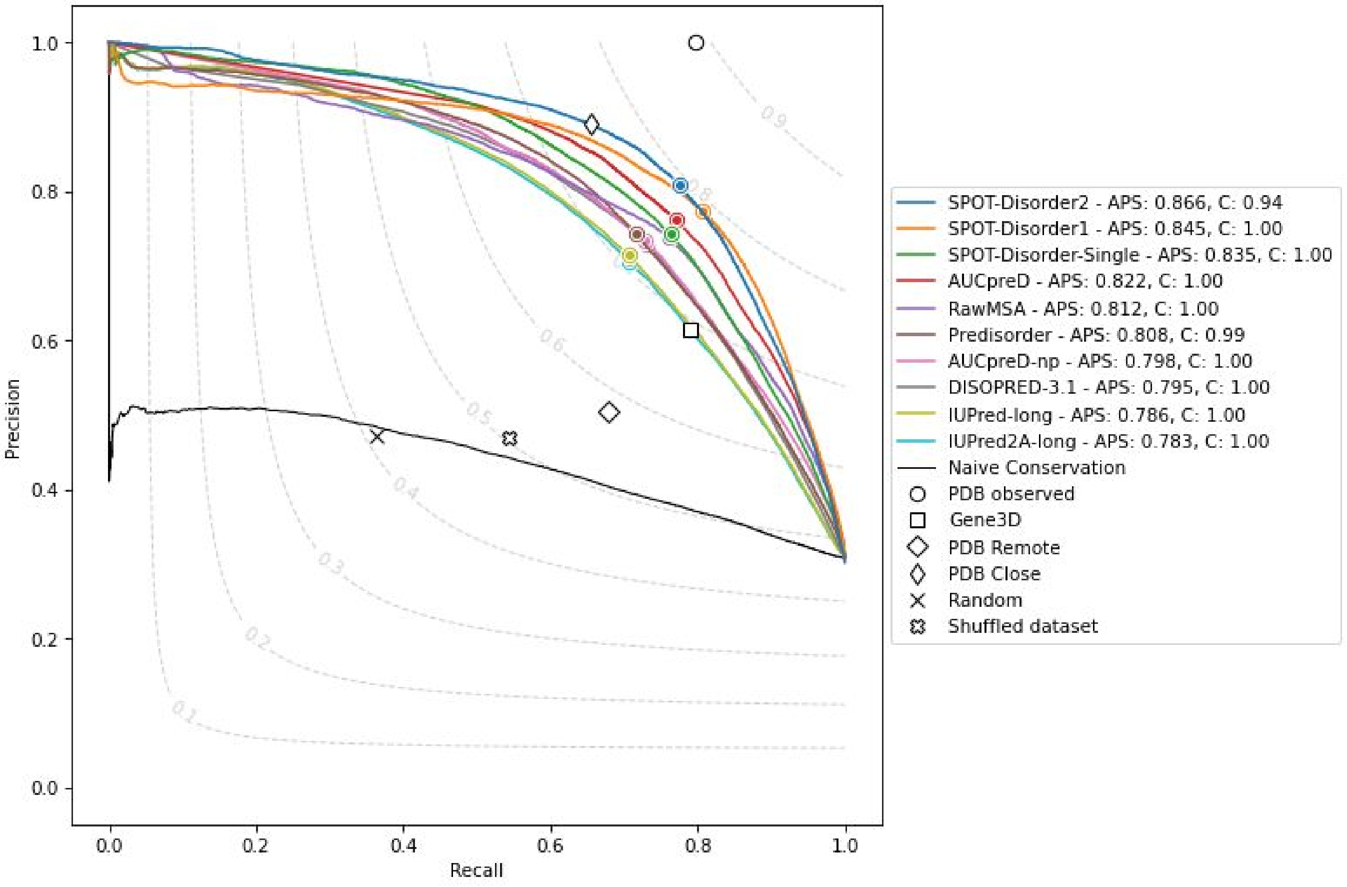


**Supplementary Figure 29**: Precision (y-axis) recall (x-axis) curves of the 10 best ranking methods. Ranking is based on their APS (average precision score) on the *DisProt-PDB* dataset.

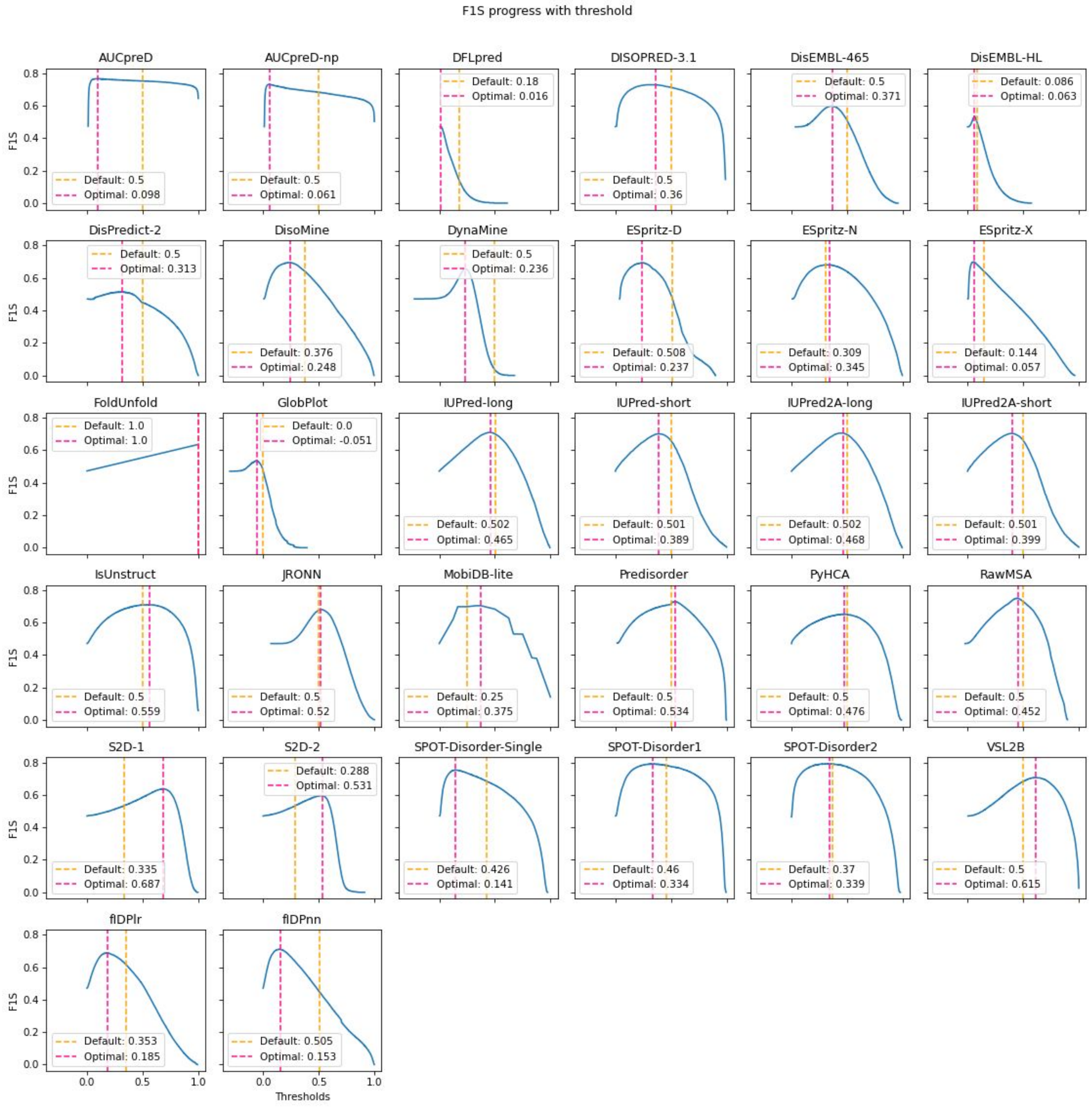


**Supplementary Figure 30**. F1-score progress (y-axis) with increasing threshold value (x-axis) for each predictor on the *DisProt-PDB* dataset.

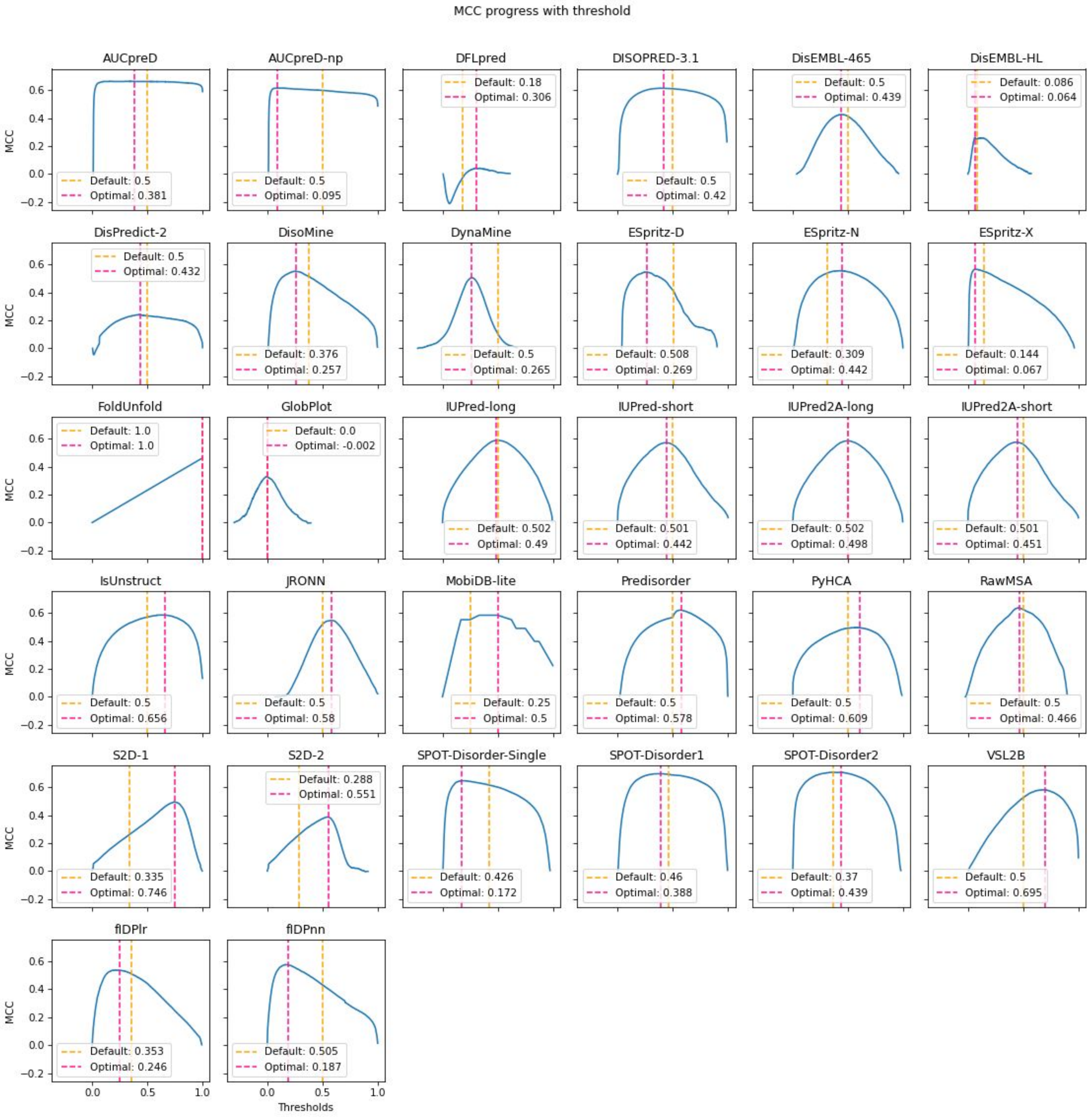


**Supplementary Figure 31**. MCC progress (y-axis) with increasing threshold value (x-axis) for each predictor on the *DisProt-PDB* dataset.

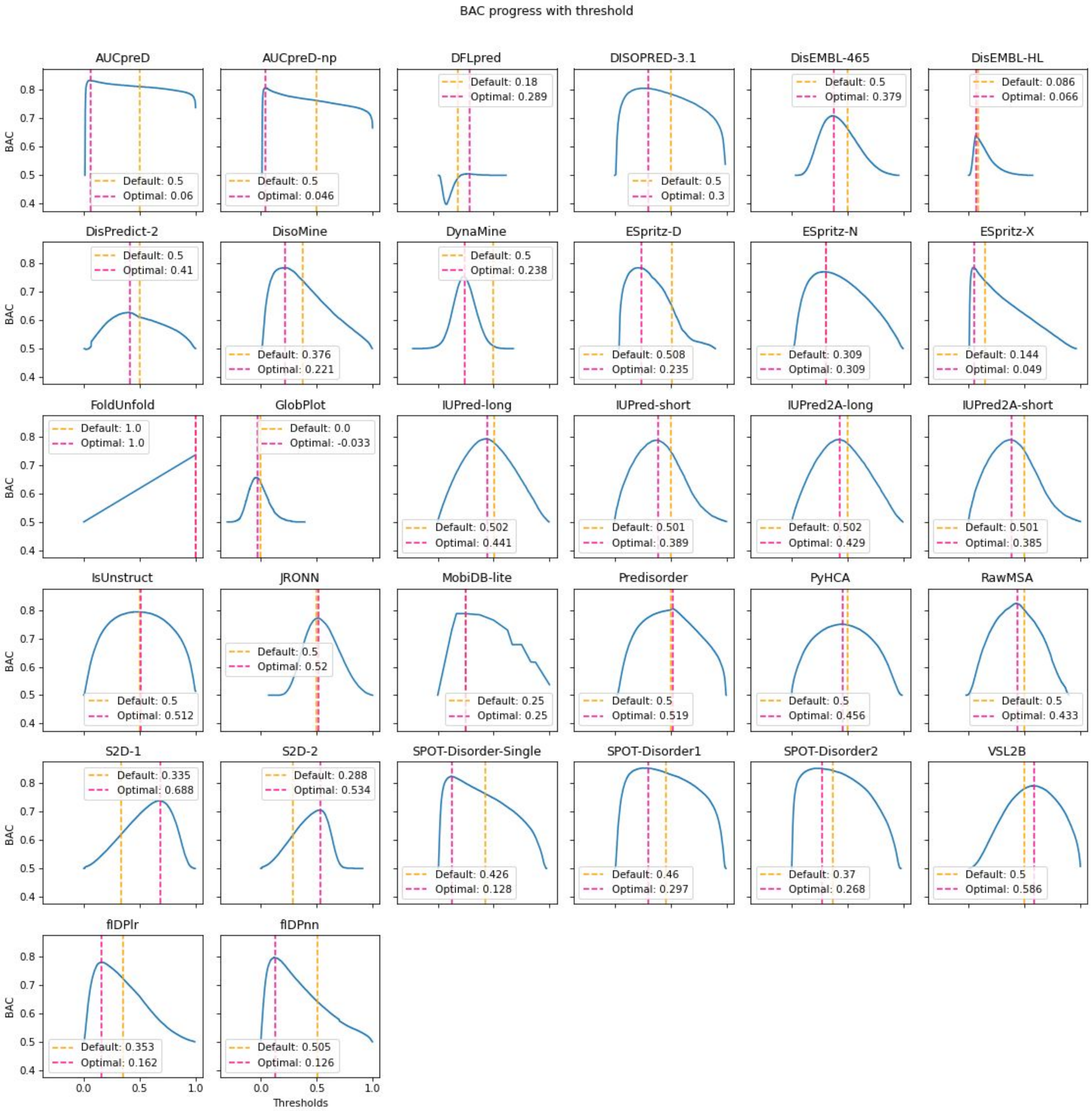


**Supplementary Figure 32**. Balanced accuracy progress (y-axis) with increasing threshold value (x-axis) for each predictor on the *DisProt-PDB* dataset.

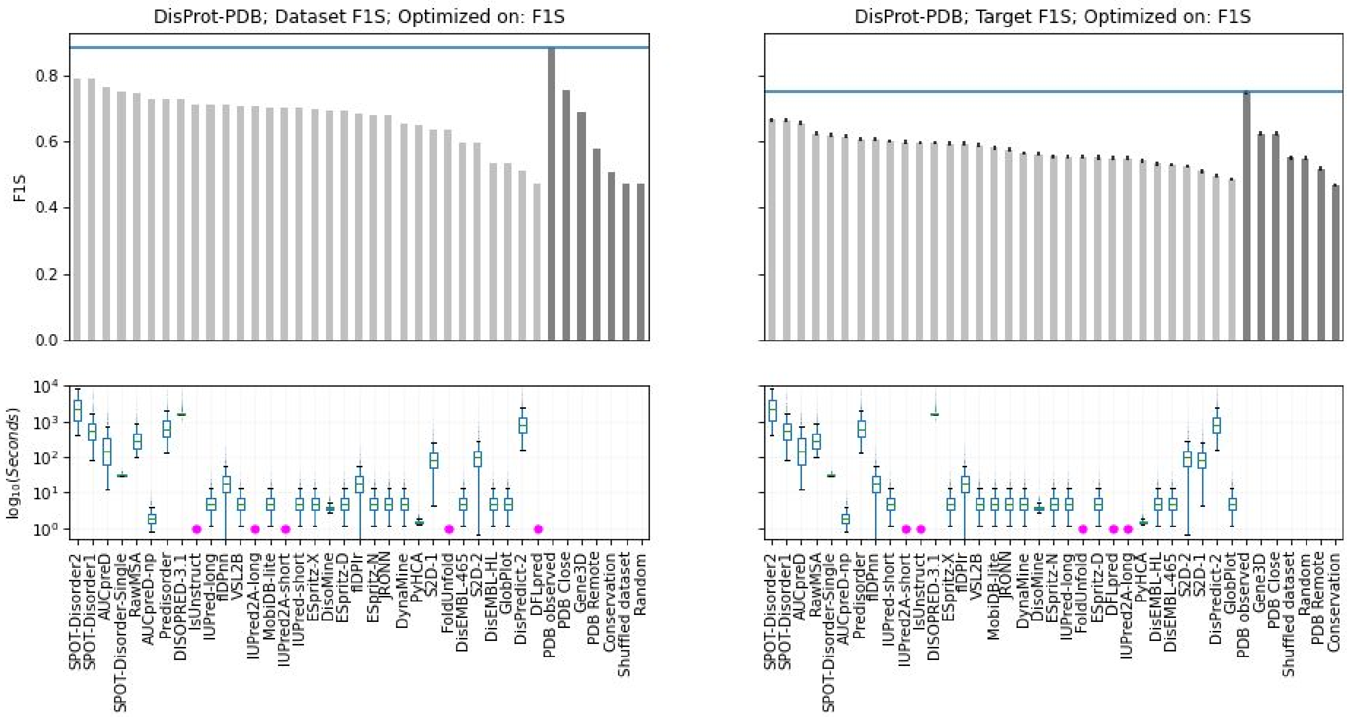


**Supplementary Figure 33**: $F_{Max}$ calculated on the whole dataset with confidence intervals as error bars (left) and averaged over proteins with Standard-Error as error bars (right). Calculated on *DisProt-PDB* dataset. Magenta dots indicate that the whole distribution of execution-times is lower than 1 second.

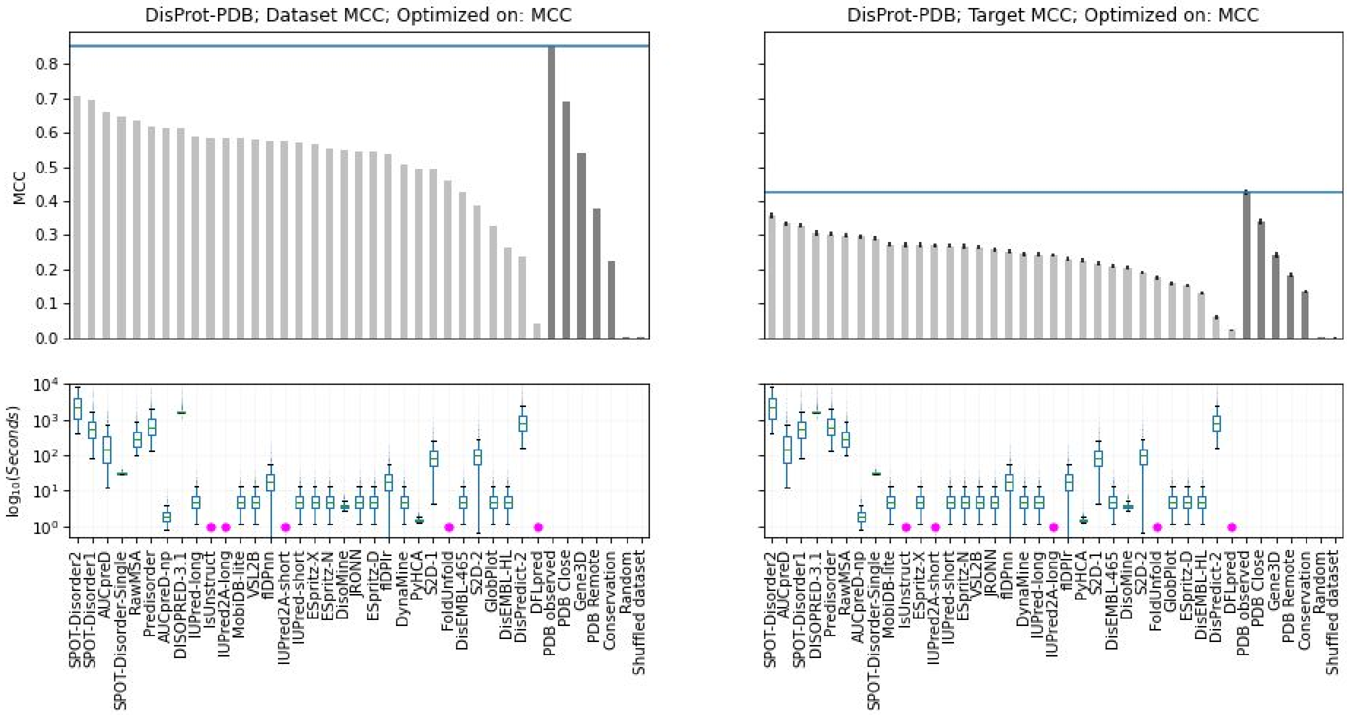


**Supplementary Figure 34**: MCC calculated on the whole dataset with confidence intervals as error bars (left) and averaged over proteins with Standard-Error as error bars (right). Calculated on *DisProt-PDB* dataset. Predictors threshold is optimized on MCC. Magenta dots indicate that the whole distribution of execution-times is lower than 1 second.

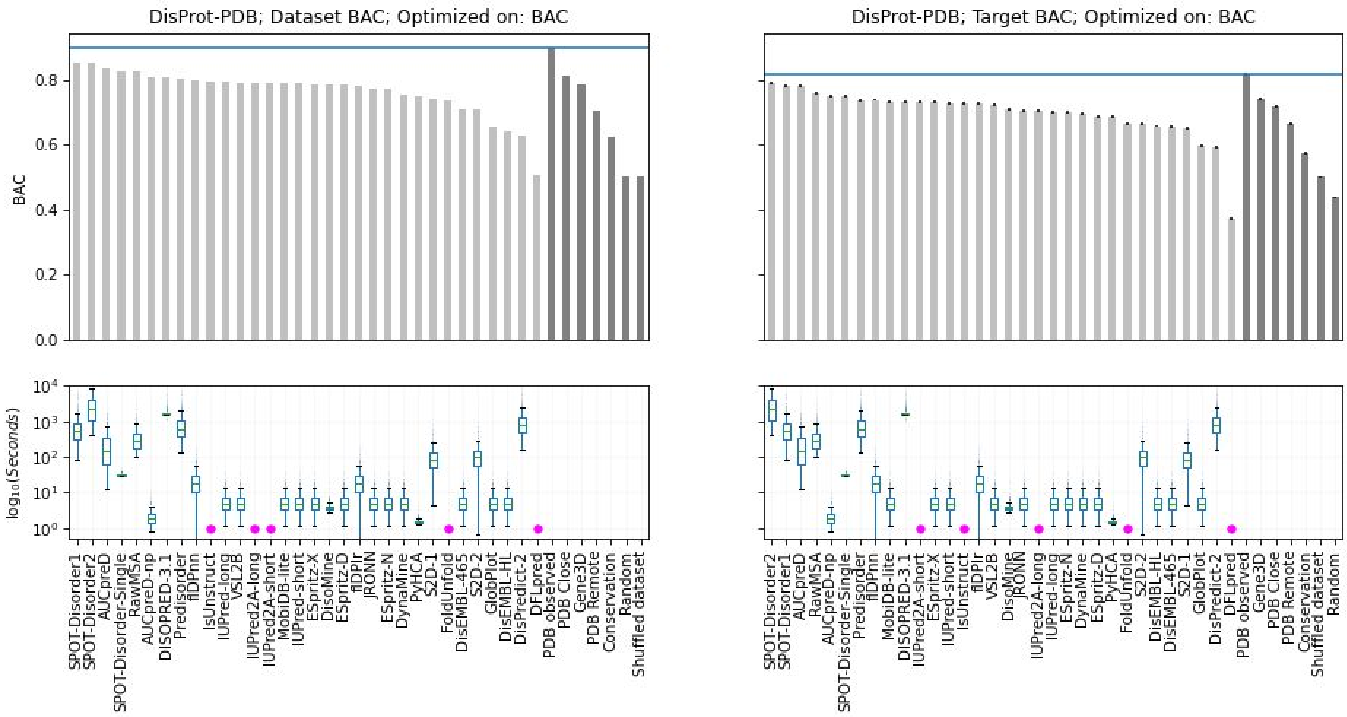


**Supplementary Figure 35**: Balanced accuracy calculated on the whole dataset with confidence intervals as error bars (left) and averaged over proteins with Standard-Error as error bar (right). Calculated on *DisProt-PDB* dataset. Predictors threshold is optimized on Balanced accuracy. Magenta dots indicate that the whole distribution of execution-times is lower than 1 second.

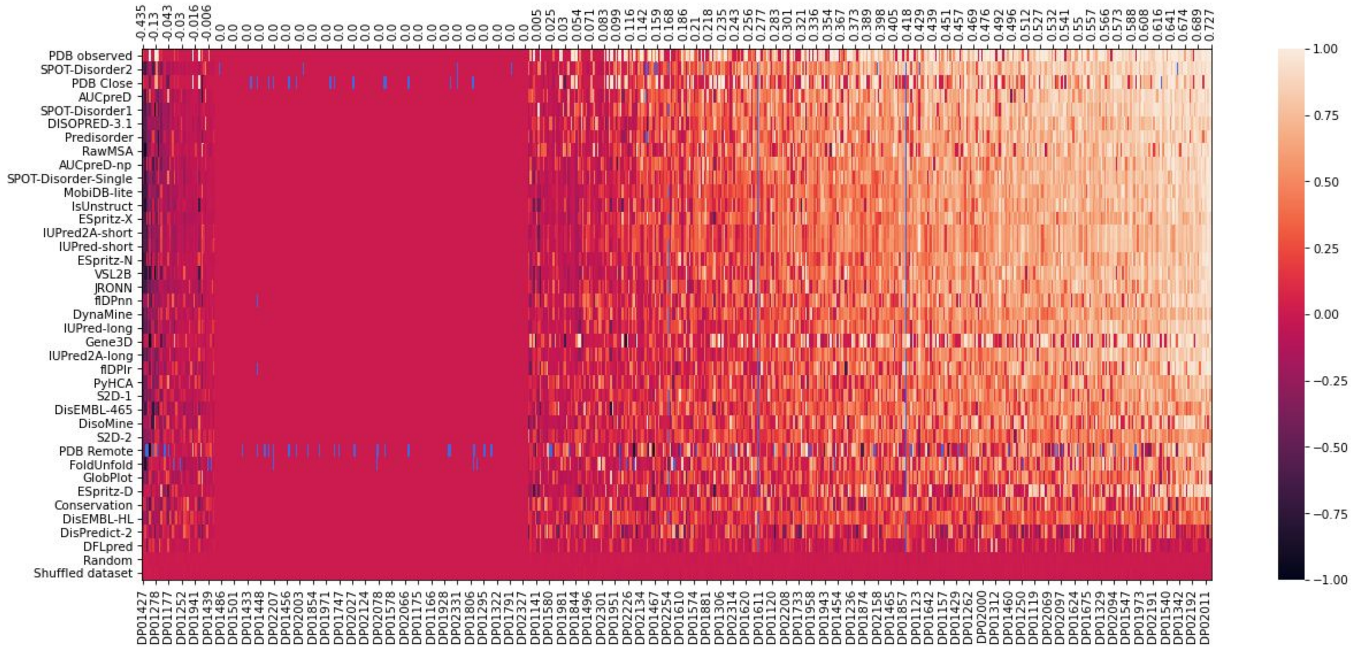


**Supplementary Figure 36**: MCC of each target (x-axis, bottom labels, not all labels are visible) from each predictor (y-axis). Targets are sorted by average MCC (x-axis, top labels). Calculated on *DisProt-PDB* dataset. Predictors threshold is optimized on MCC. Missing values are in blue.

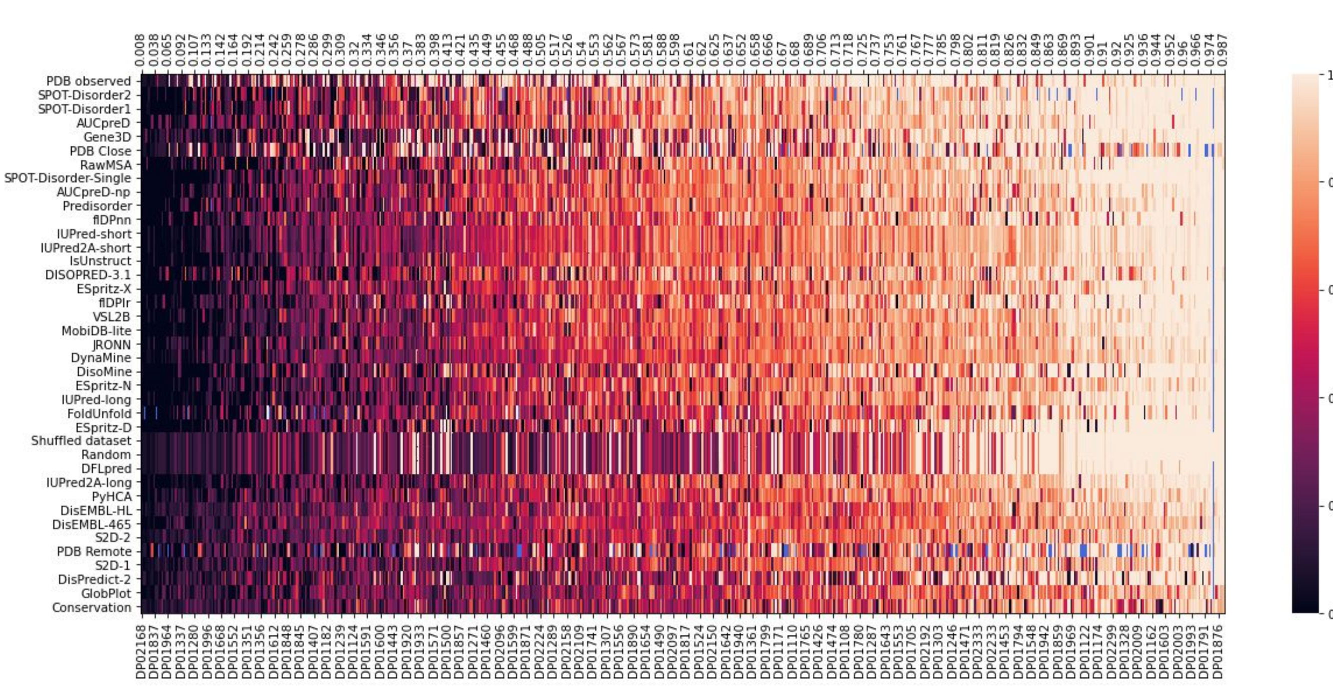


**Supplementary Figure 37**: $F_{Max}$ of each target (x-axis, bottom labels, not all labels are visible) from each predictor (y-axis). Targets are sorted by average $F_{Max}$ (x-axis, top labels). Calculated on *DisProt-PDB* dataset. Missing values are in blue.

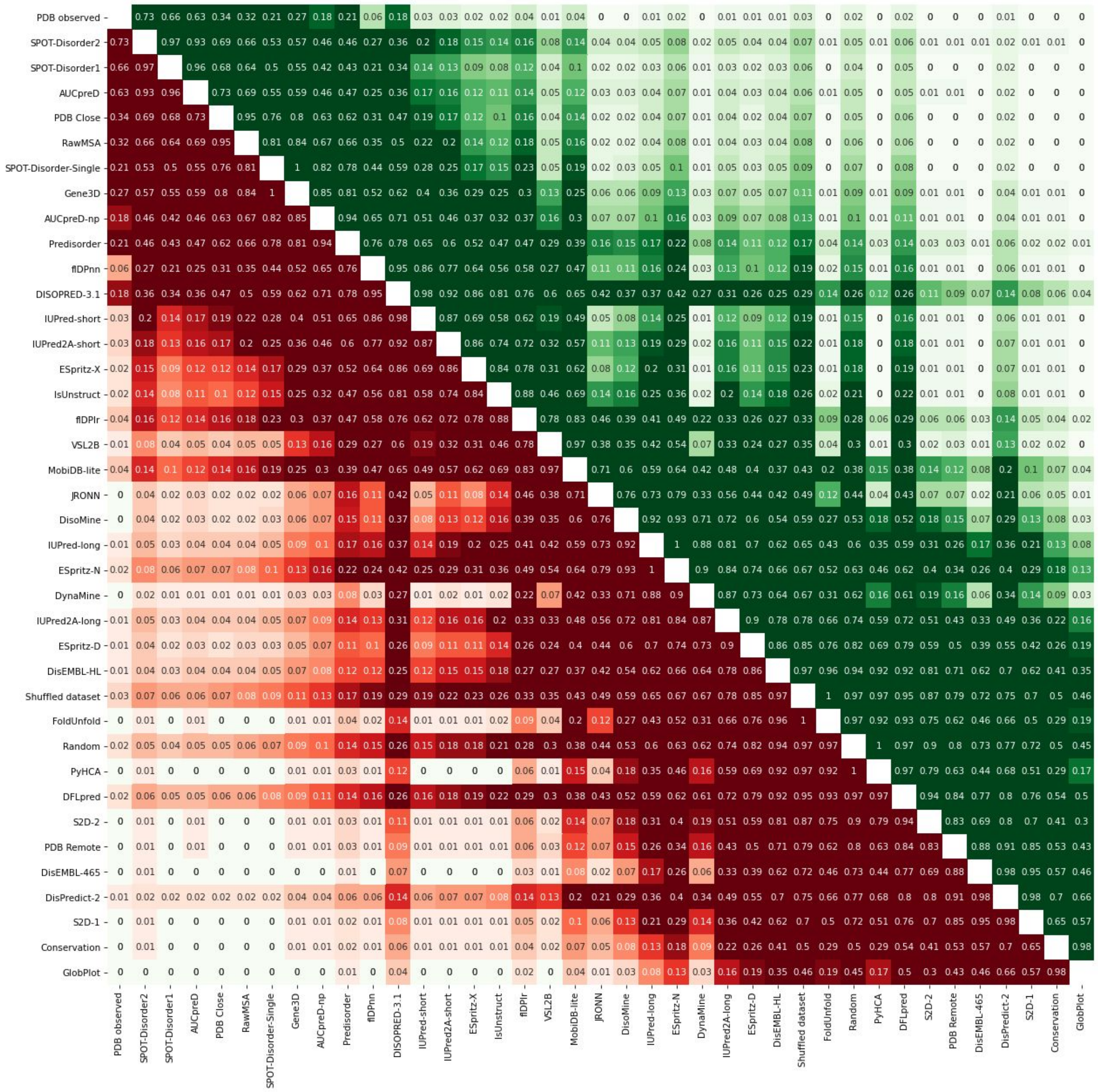


**Supplementary Figure 38**. Overall average ranking of all predictors and baselines in the *DisProt-PDB* dataset. Heatmap of the T-test p-value associated to the statistical significance of the difference between ranking distribution of predictors. A ranking distribution for a predictor is the position of that predictor in its ranking for each metric. Metrics used are: bac, f1s, fpr, mcc, ppv, tpr, tnr; they are calculated with predictors threshold optimized by F1-Score.

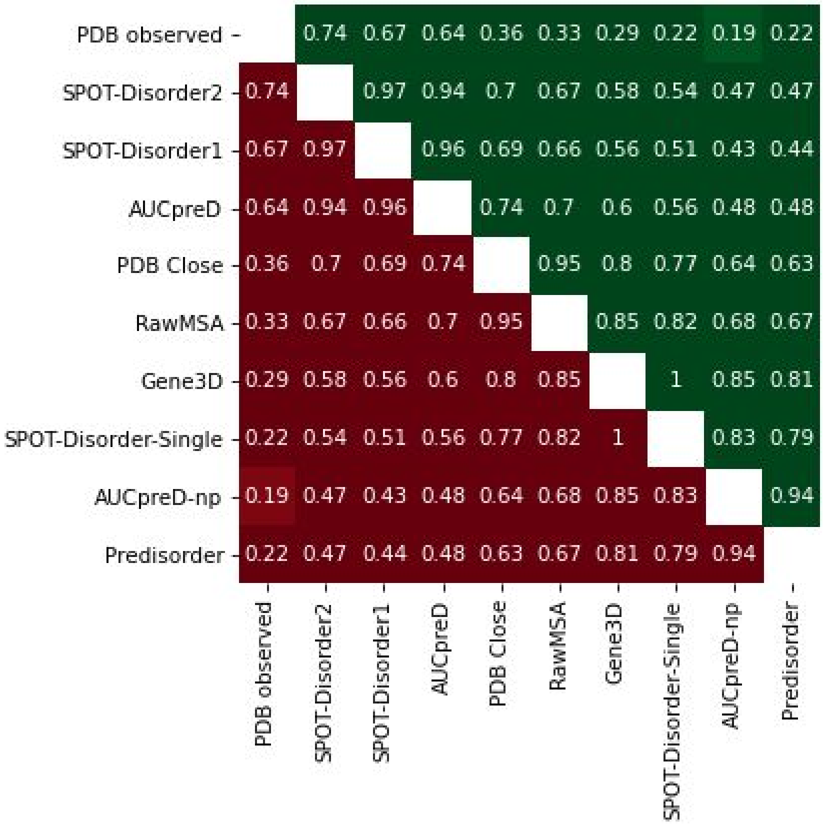


**Supplementary Figure 39**. Overall average ranking of the 10 best ranking predictors and baselines in the *DisProt-PDB* dataset. Heatmap of the T-test p-value associated to the statistical significance of the difference between ranking distribution of predictors. A ranking distribution for a predictor is the position of that predictor in its ranking for each metric. Metrics used are: bac, f1s, fpr, mcc, ppv, tpr, tnr; they are calculated with predictors threshold optimized by F1-Score.

Mammals

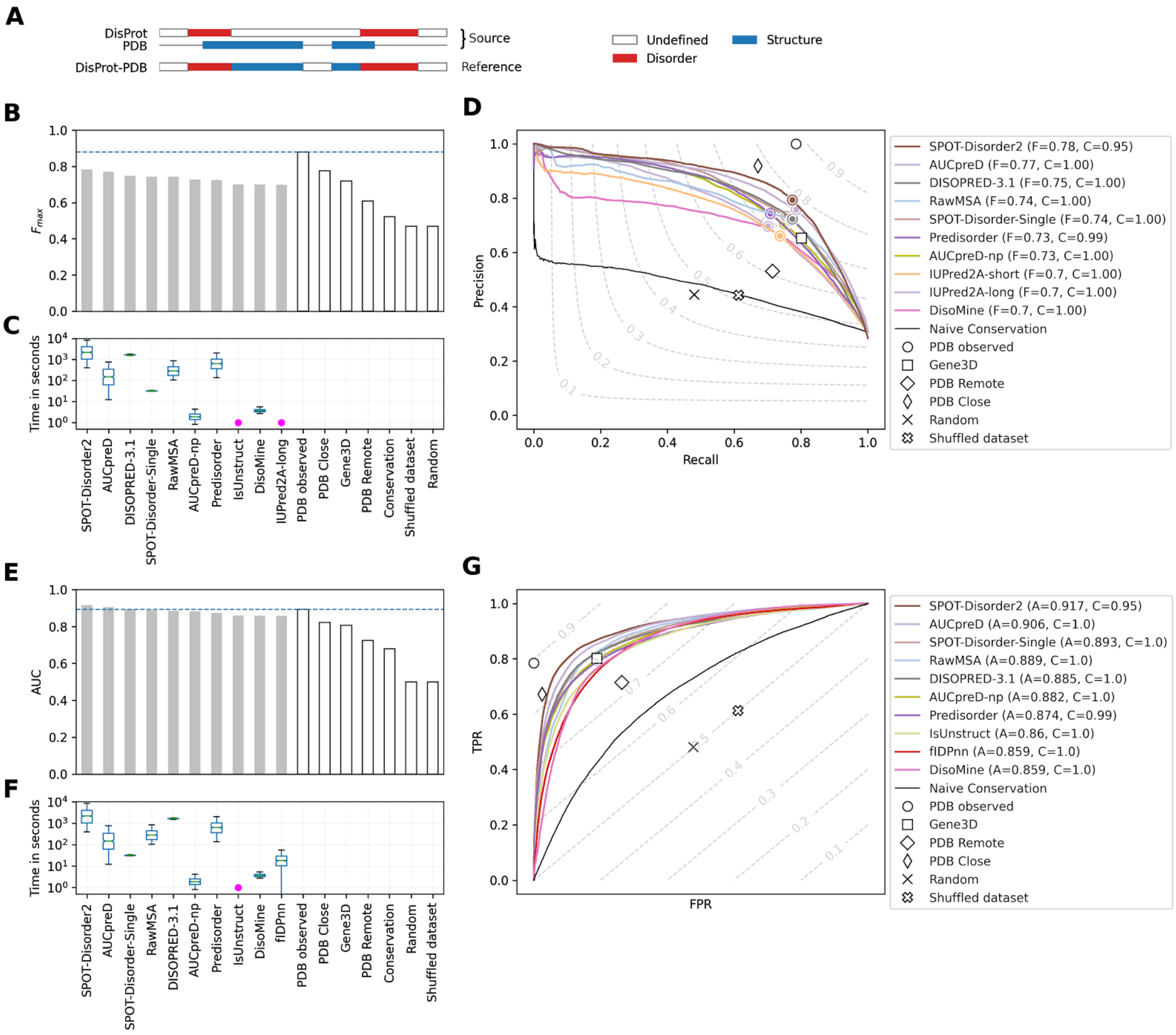


**Supplementary Figure 40.** Prediction success and CPU times for the ten top-ranking disorder predictors for mammalian proteins in the *DisProt-PDB* dataset. Reference used (*DisProt-PDB*) in the analysis and how it is obtained (panel A). Performance of predictors expressed as maximum F1-Score across all thresholds ($F_{max}$) (panel B) and AUC (panel E) for the top ten best ranking methods (light gray) and baselines (white) and the distribution of execution time per-target (panels C, F) using *DisProt-PDB* dataset. The horizontal line in panels B, E indicates the $F_{max}$ and AUC of the best baseline, respectively. Precision-Recall (panel D) and ROC curves (panel G) of ten top-ranking methods and baselines using *DisProt-PDB* dataset, with level curves of the F1-Score and Balanced accuracy, respectively. Magenta dots on panels C, F indicate that the whole distribution of execution-times is lower than 1 second.

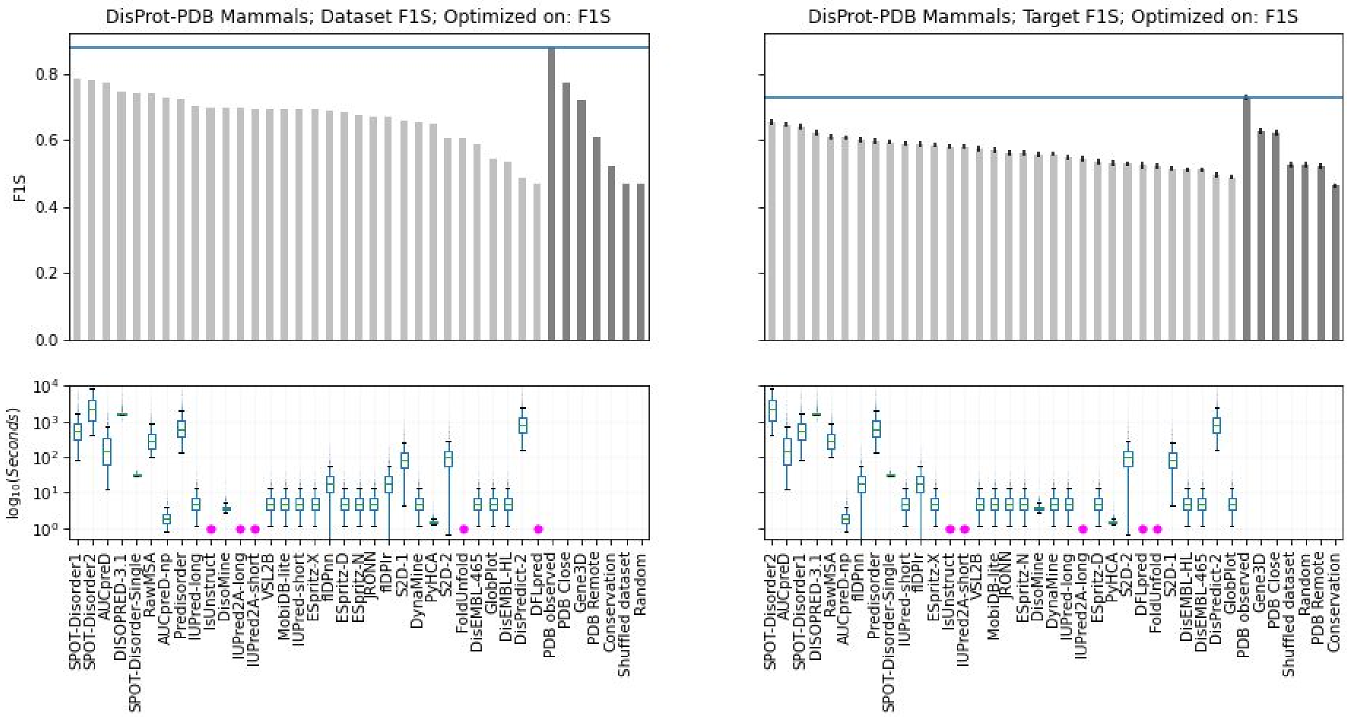


**Supplementary Figure 41**: $F_{Max}$ calculated on the whole dataset with confidence intervals as error bars (left) and averaged over proteins with Standard-Error as error bars (right). Calculated on mammalian proteins of the *DisProt-PDB* dataset.  Magenta dots indicate that the whole distribution of execution-times is lower than 1 second.

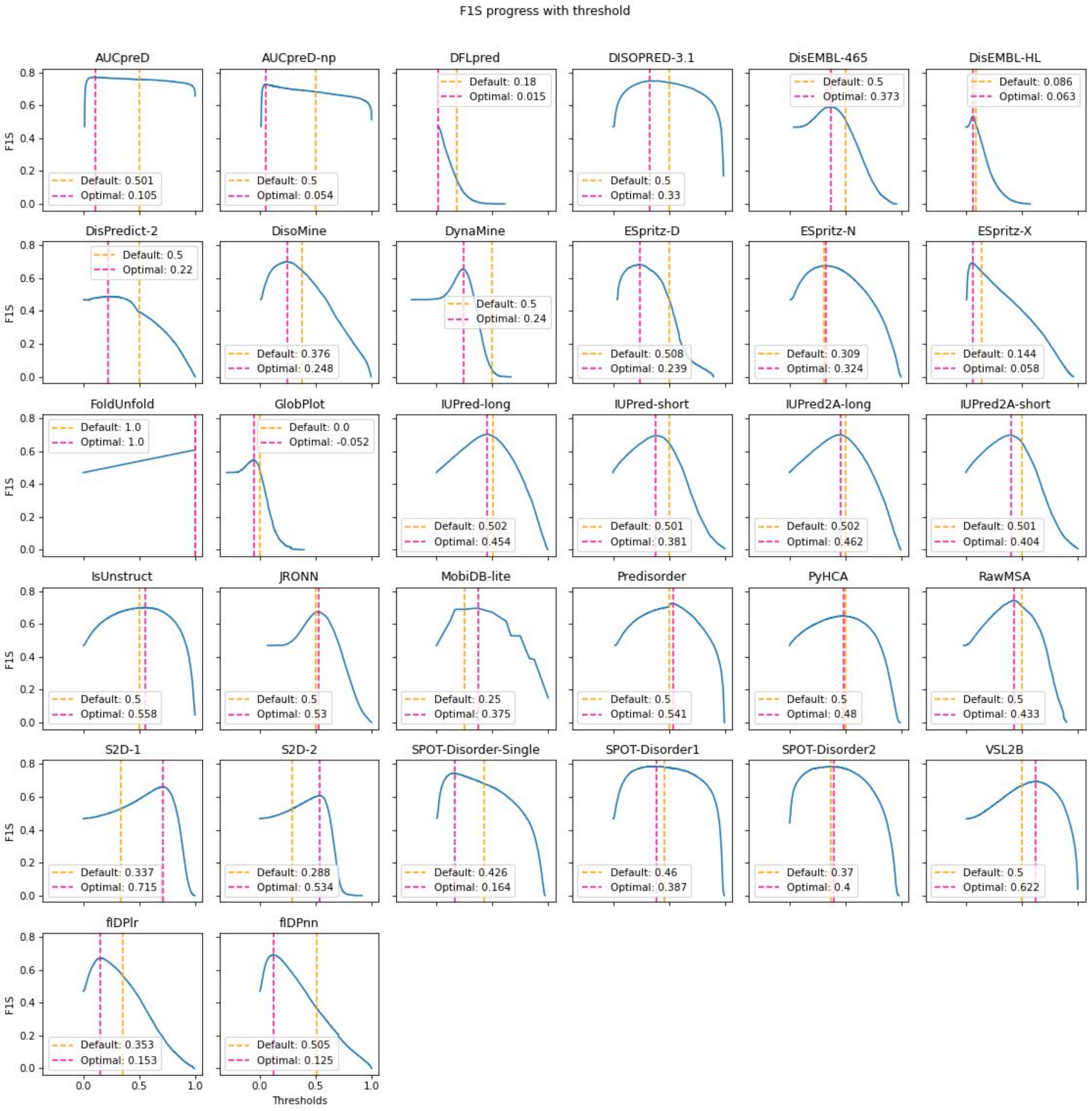


**Supplementary Figure 42**. F1-score progress (y-axis) with increasing threshold value (x-axis) for each predictor calculated on mammalian proteins on the *DisProt-PDB* dataset.

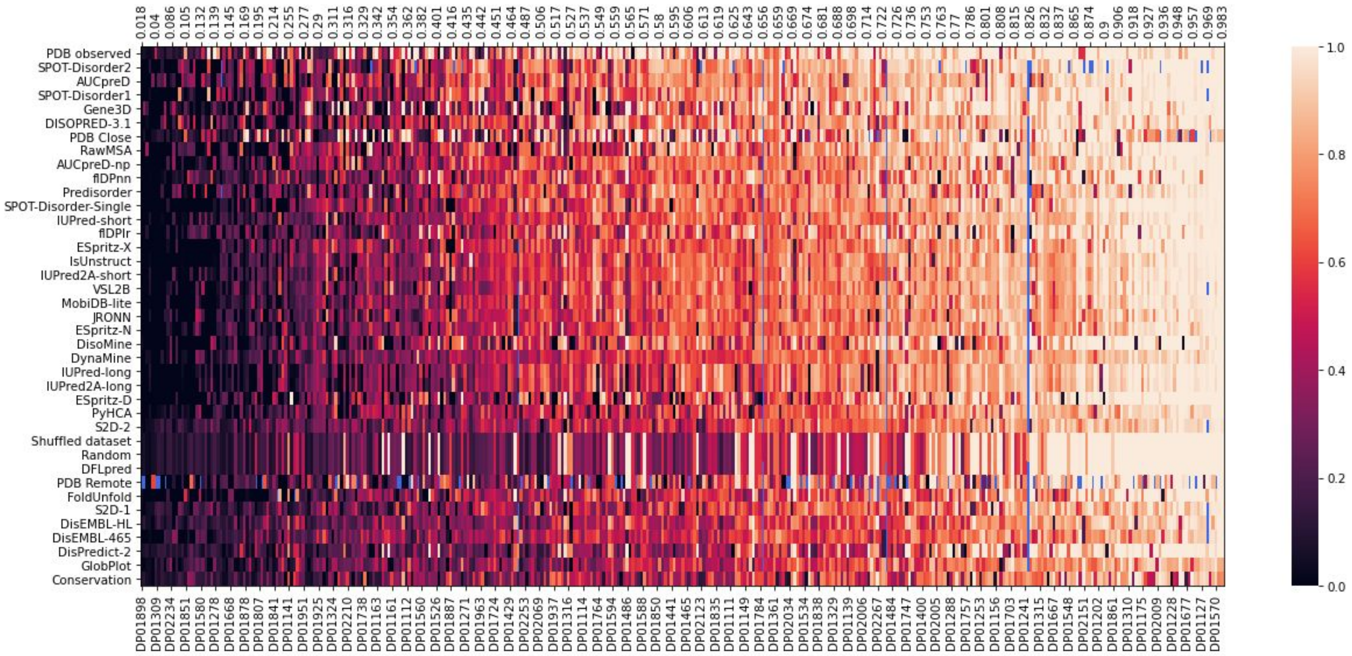


**Supplementary Figure 43**: $F_{Max}$ of each target (x-axis, bottom labels) from each predictor (y-axis). Targets are sorted by average $F_{Max}$ (x-axis, top labels). Calculated on mammalian proteins of the *DisProt-PDB* dataset. Missing values are in blue.

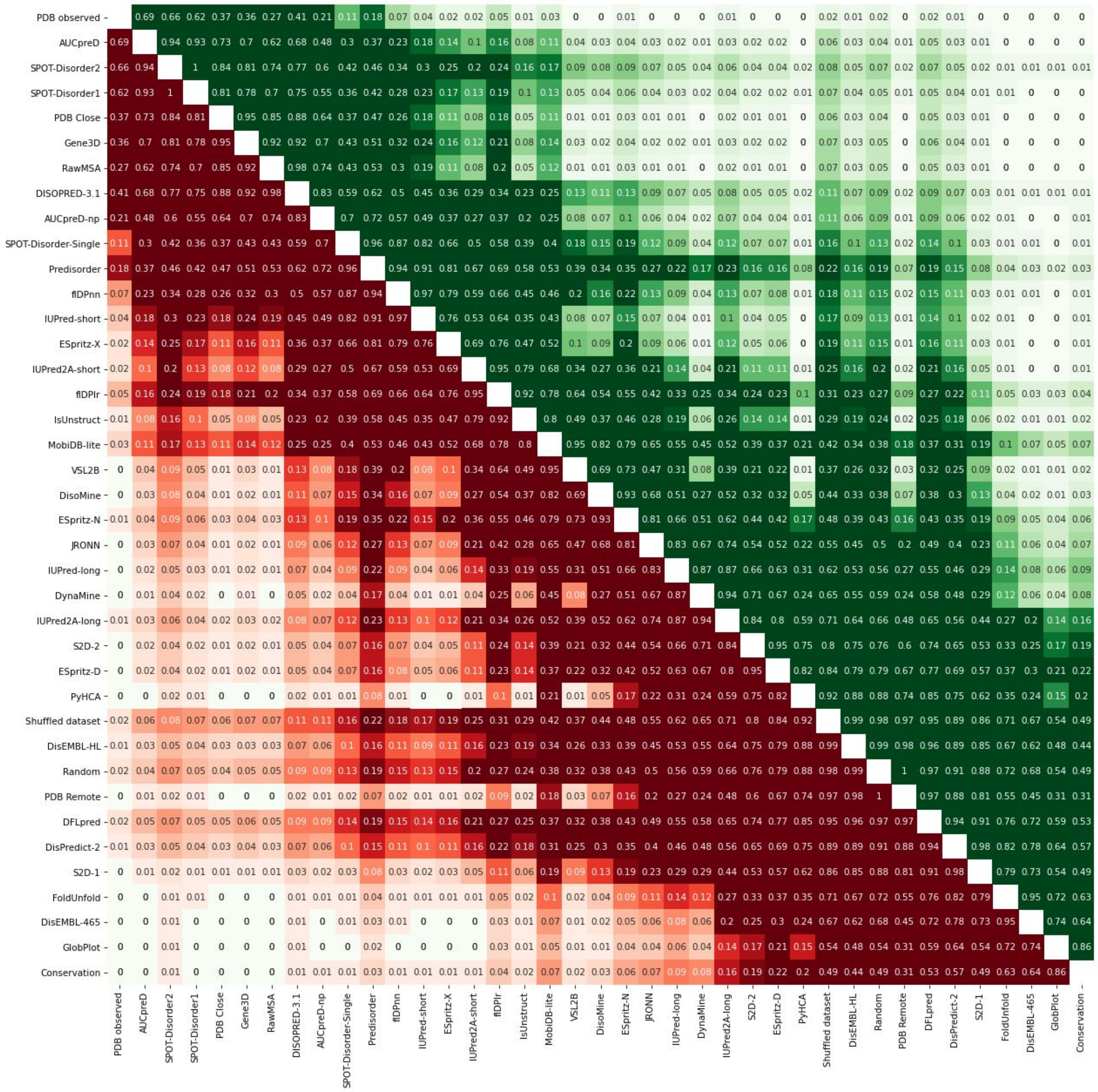

**Supplementary Figure 44**. Overall average ranking of all predictors and baselines for mammalian proteins in the *DisProt-PDB* dataset. Heatmap of the T-test p-value associated to the statistical significance of the difference between ranking distribution of predictors. A ranking distribution for a predictor is the position of that predictor in its ranking for the following metrics: 'bac', 'f1s', 'fpr', 'mcc', 'ppv', 'tpr', 'tnr'. Metrics are calculated per target and with predictors threshold optimized by F1-Score.

Prokaryotes

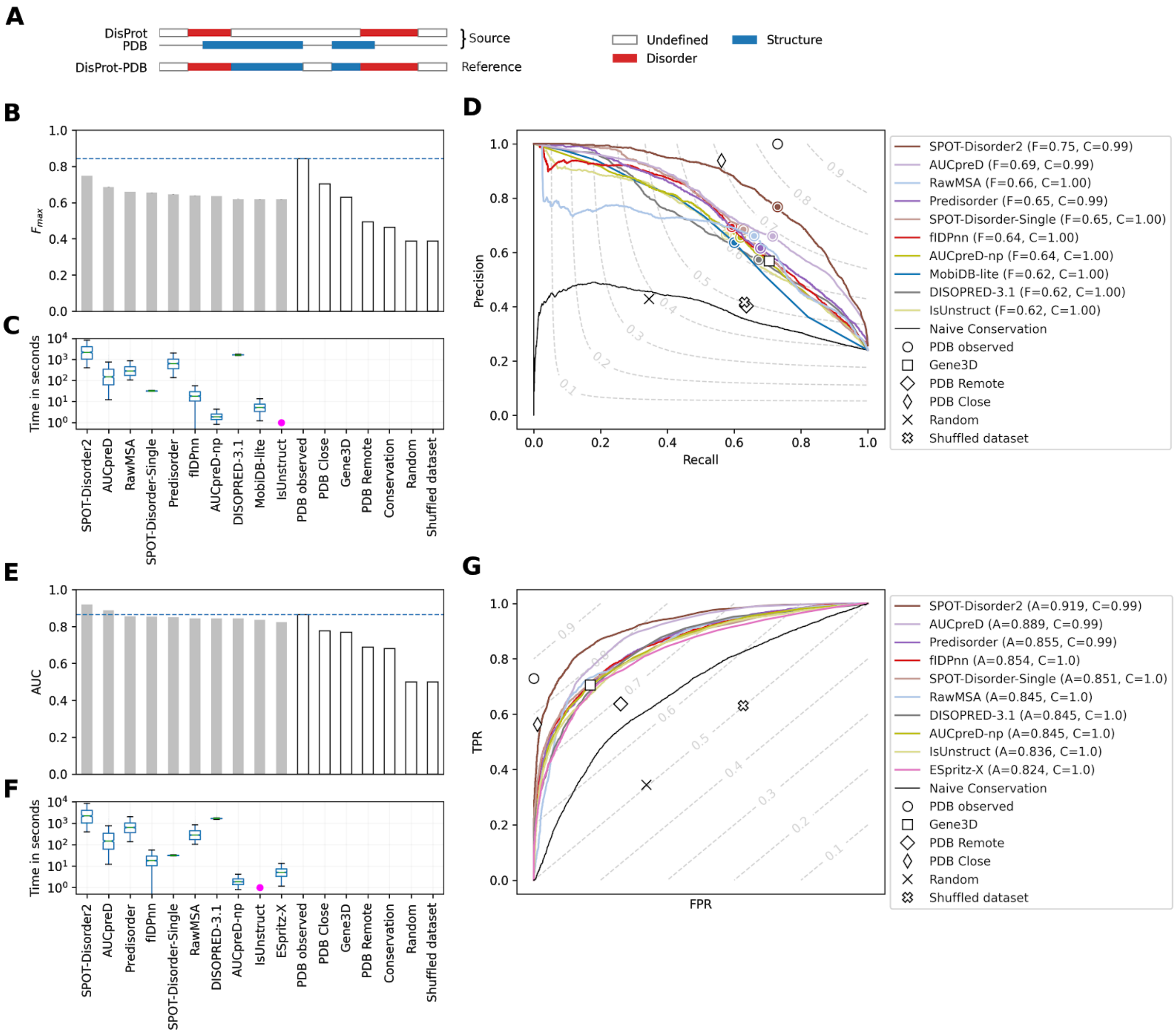


**Supplementary Figure 45.** Prediction success and CPU times for the ten top-ranking disorder predictors for prokaryotic proteins in the *DisProt-PDB* dataset. Reference used (*DisProt-PDB*) in the analysis and how it is obtained (panel A). Performance of predictors expressed as maximum F1-Score across all thresholds ($F_{max}$) (panel B) and AUC (panel E) for the top ten best ranking methods (light gray) and baselines (white) and the distribution of execution time per-target (panels C, F) using *DisProt-PDB* dataset. The horizontal line in panels B, E indicates the $F_{max}$ and AUC of the best baseline, respectively. Precision-Recall (panel D) and ROC curves (panel G) of ten top-ranking methods and baselines using *DisProt-PDB* dataset, with level curves of the F1-Score and Balanced accuracy, respectively. Magenta dots on panels C, F indicate that the whole distribution of execution-times is lower than 1 second.

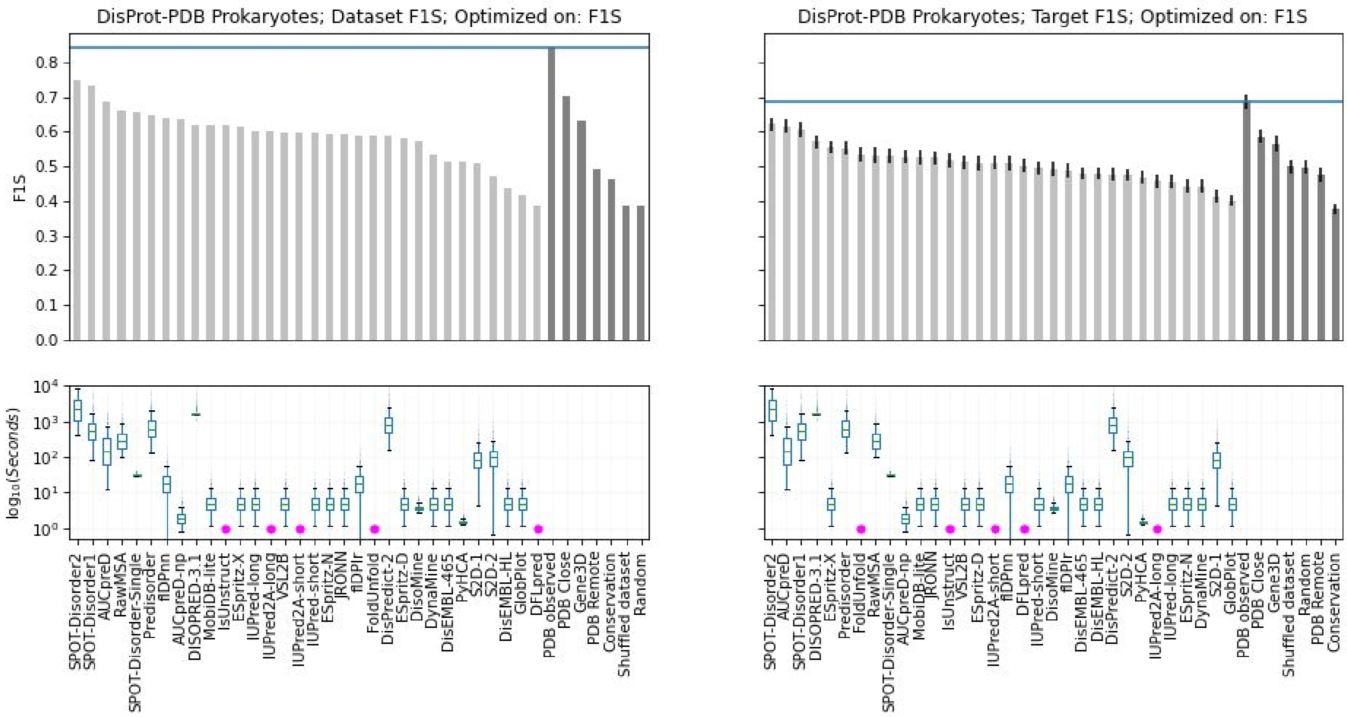


**Supplementary Figure 46**: $F_{Max}$ calculated on the whole dataset with confidence intervals as error bars (left) and averaged over proteins with Standard-Error as error bars (right). Calculated on prokaryotic proteins of the *DisProt-PDB* dataset. Magenta dots indicate that the whole distribution of execution-times is lower than 1 second.

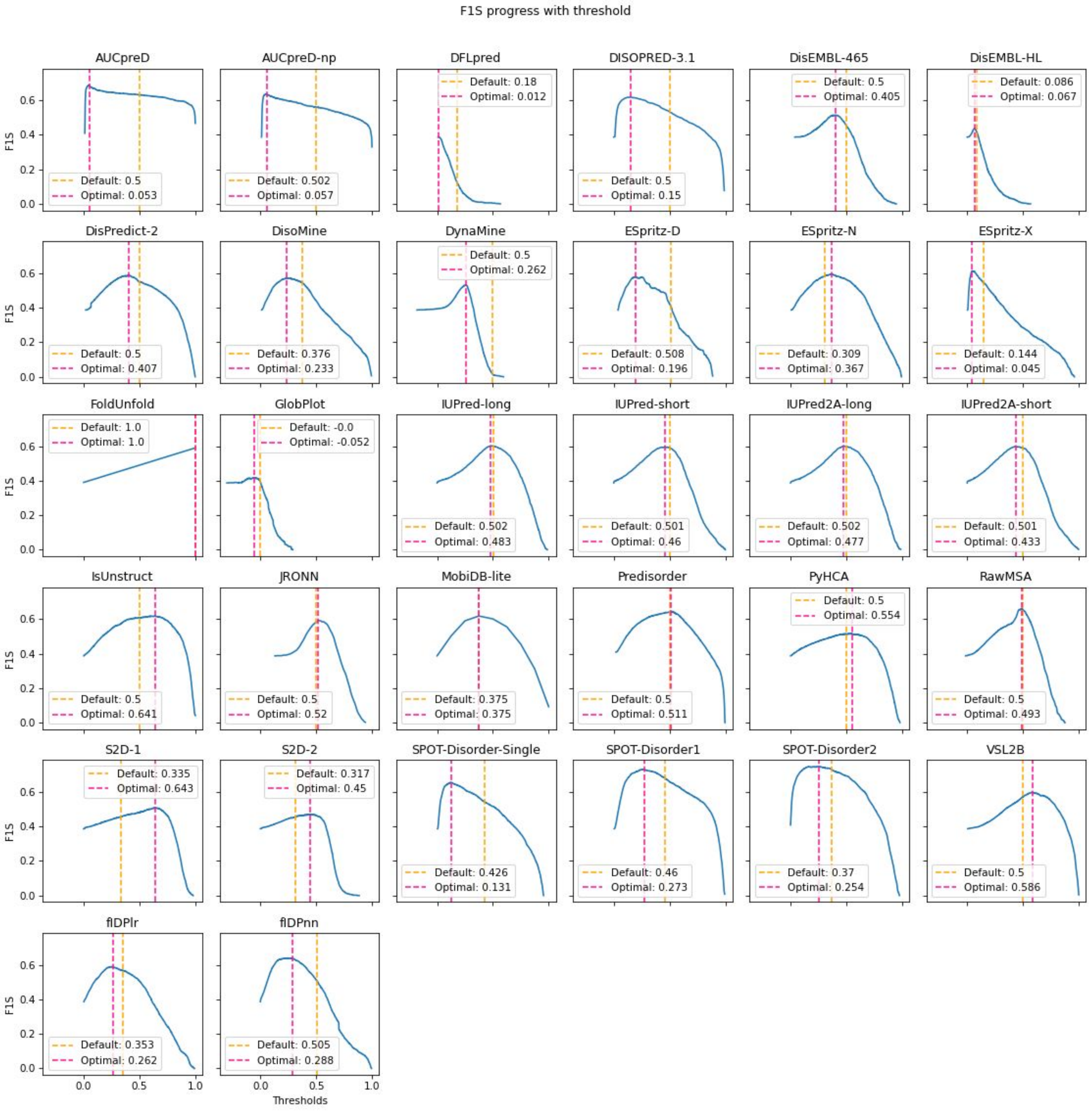


**Supplementary Figure 47**. F1-score progress (y-axis) with increasing threshold value (x-axis) for each predictor calculated on prokaryotic proteins on the *DisProt-PDB* dataset.

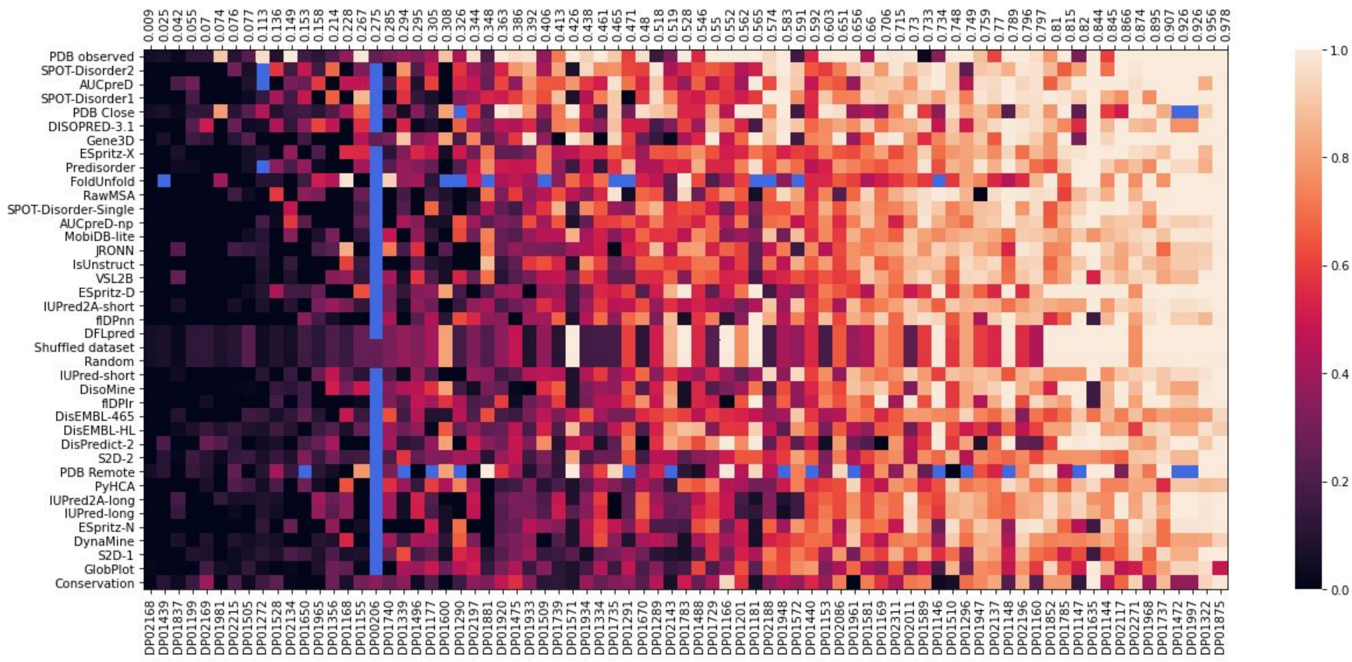

**Supplementary Figure 48**: $F_{Max}$ of each target (x-axis, bottom labels) from each predictor (y-axis). Targets are sorted by average $F_{Max}$ (x-axis, top labels). Calculated on prokaryotic proteins of the *DisProt-PDB* dataset. Missing values are in blue.

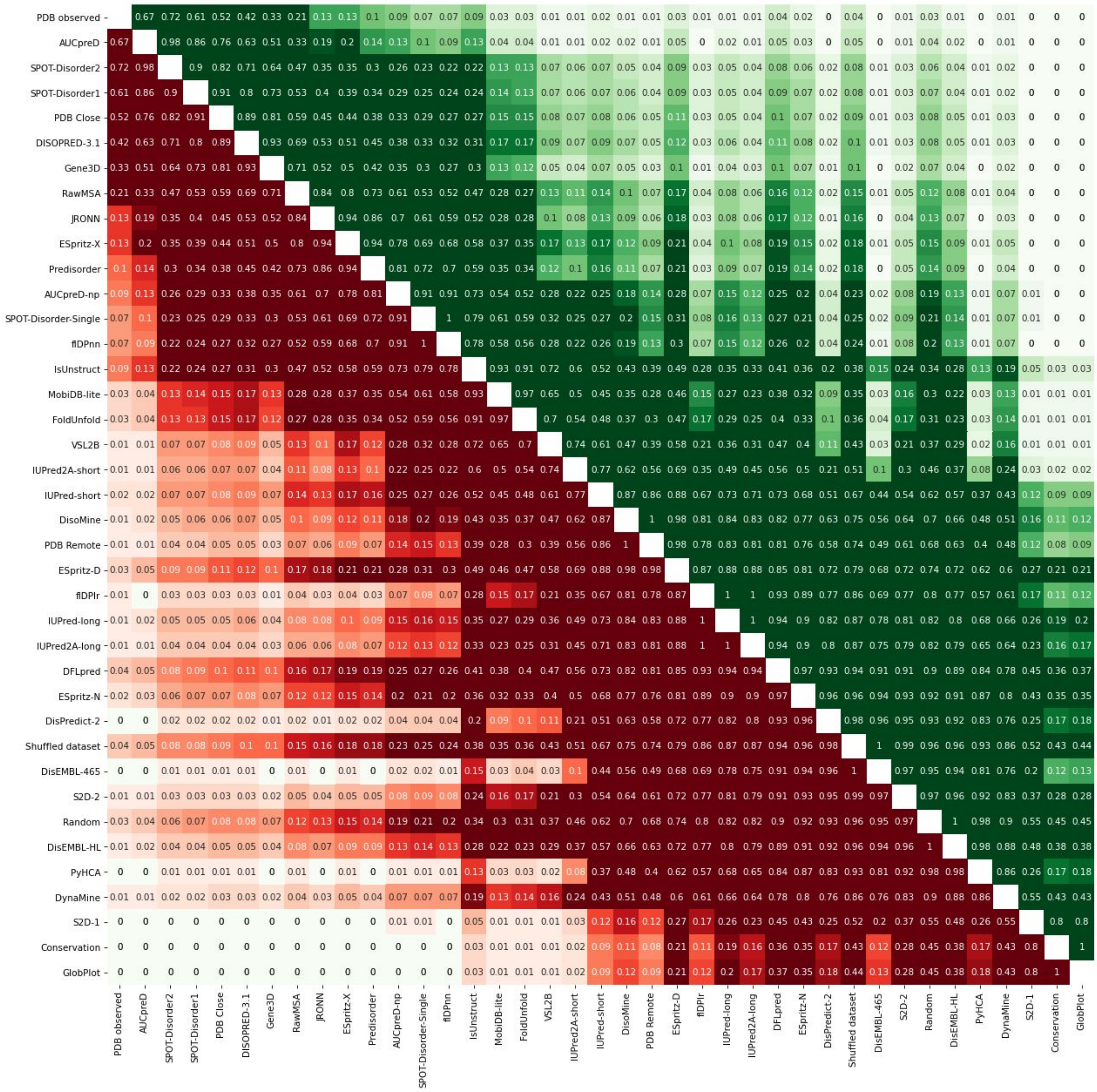

**Supplementary Figure 49**. Overall average ranking of all predictors and baselines for prokaryotic proteins in the *DisProt-PDB* dataset. Heatmap of the T-test p-value associated to the statistical significance of the difference between ranking distribution of predictors. A ranking distribution for a predictor is the position of that predictor in its ranking for the following metrics: 'bac', 'f1s', 'fpr', 'mcc', 'ppv', 'tpr', 'tnr'. Metrics are calculated per target and with predictors threshold optimized by F1-Score.

### Binding

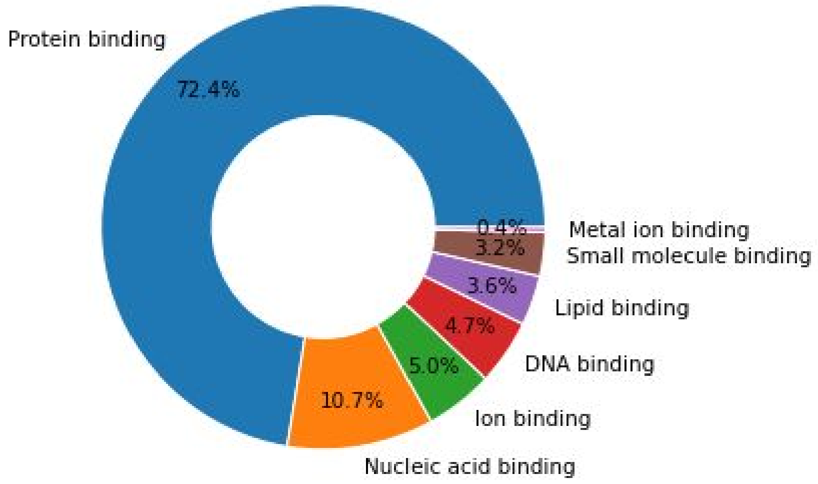


**Supplementary Figure 50**. Representation of the type of bindings among the binding regions in the binding datasets.

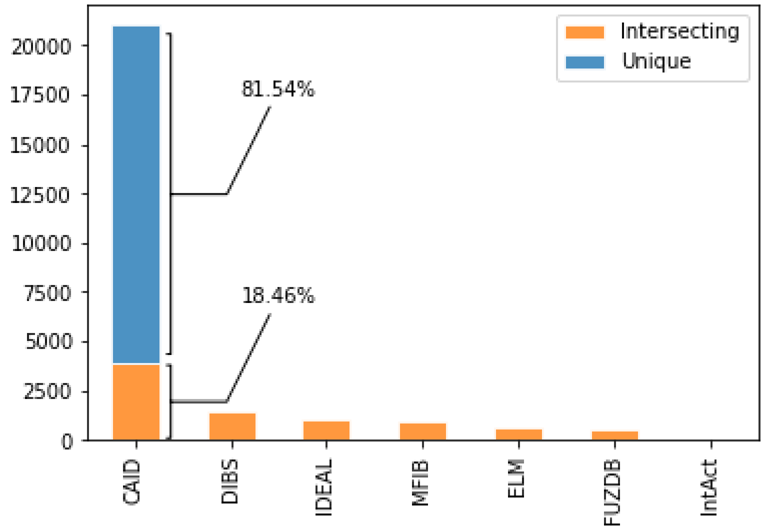


**Supplementary Figure 51**. Number and fraction of residues annotated with binding in CAID dataset that are Unique to CAID dataset (blue) or present in other datasets (orange).

#### DisProt-Binding dataset

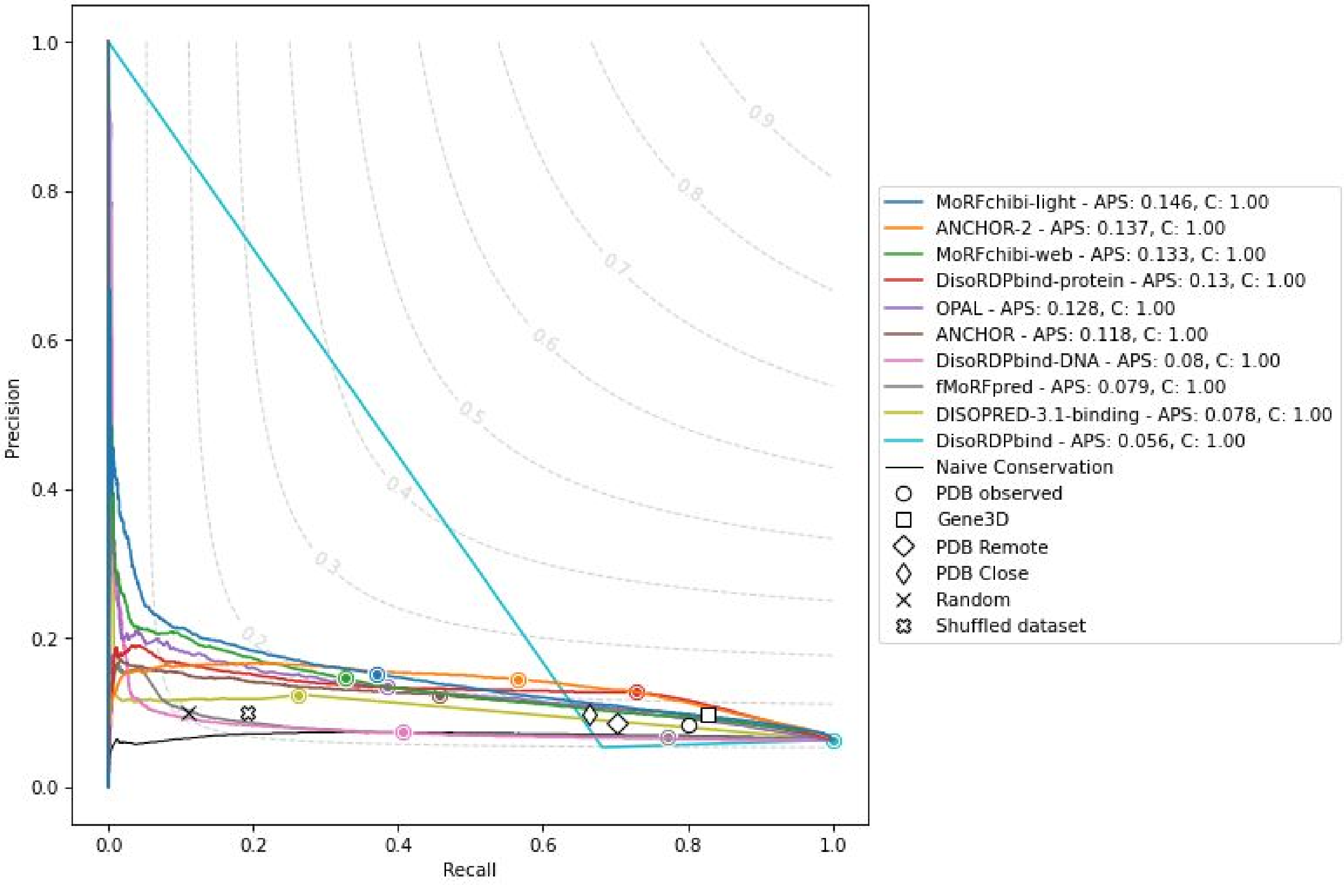


**Supplementary Figure 52**: Precision (y-axis) recall (x-axis) curves of the 10 best ranking methods. Ranking is based on their APS (average precision score) in the *DisProt-Binding* dataset.

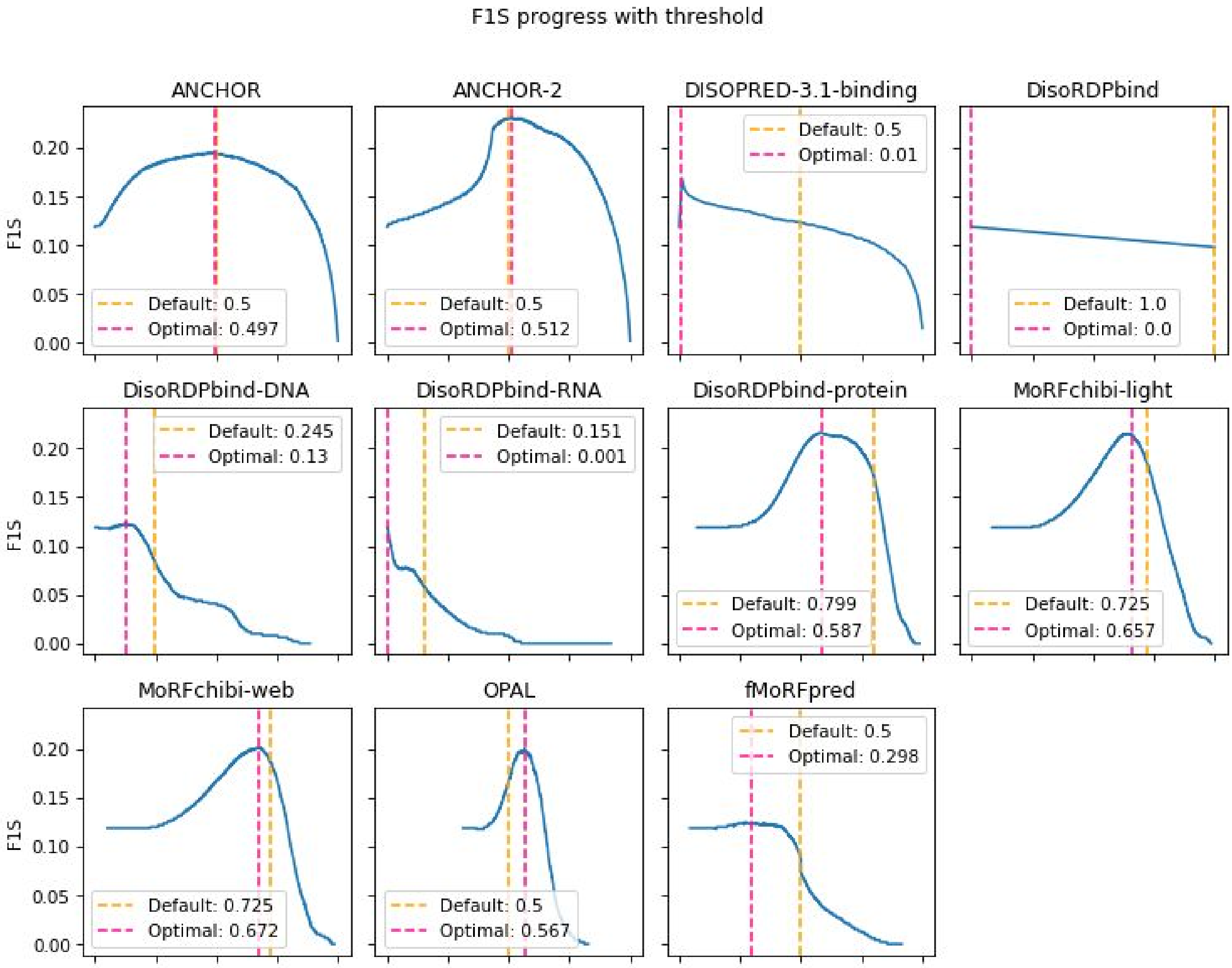


**Supplementary Figure 53**. F1-score progress (y-axis) with increasing threshold value (x-axis) for each predictor in the *DisProt-Binding* dataset.

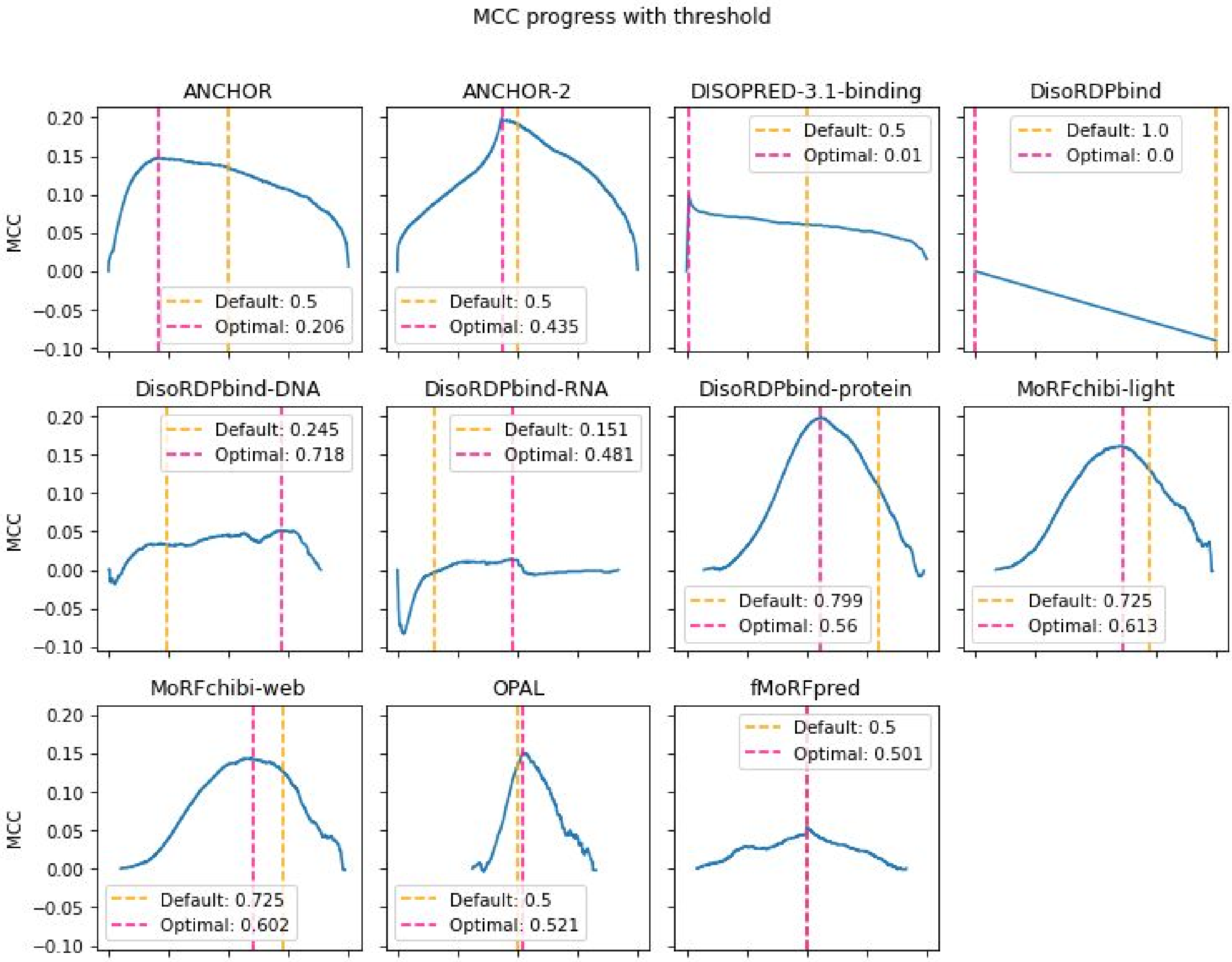


**Supplementary Figure 54**. MCC progress (y-axis) with increasing threshold value (x-axis) for each predictor in the *DisProt-binding* dataset.

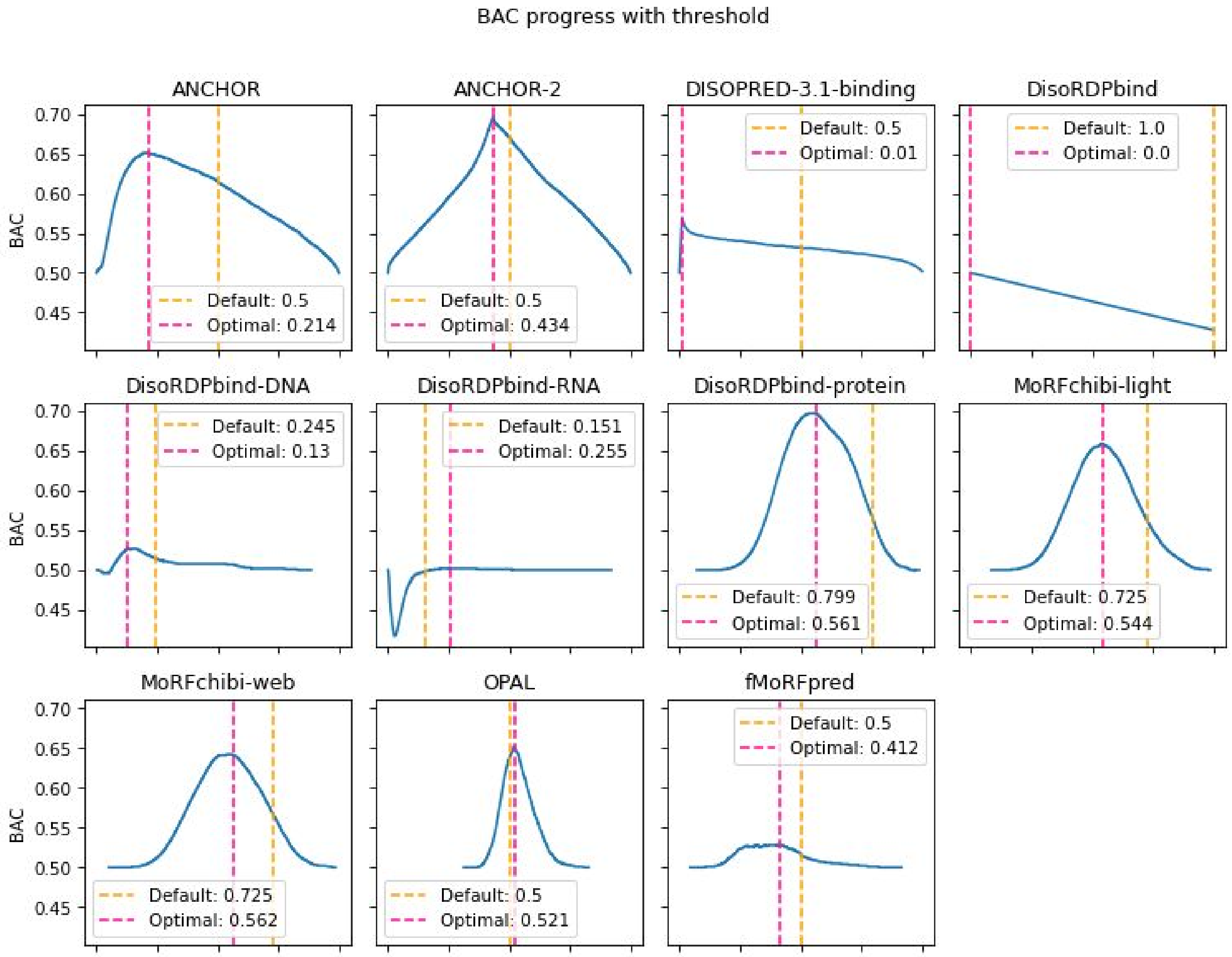


**Supplementary Figure 55**. Balanced accuracy progress (y-axis) with increasing threshold value (x-axis) for each predictor in the *DisProt-binding* dataset.

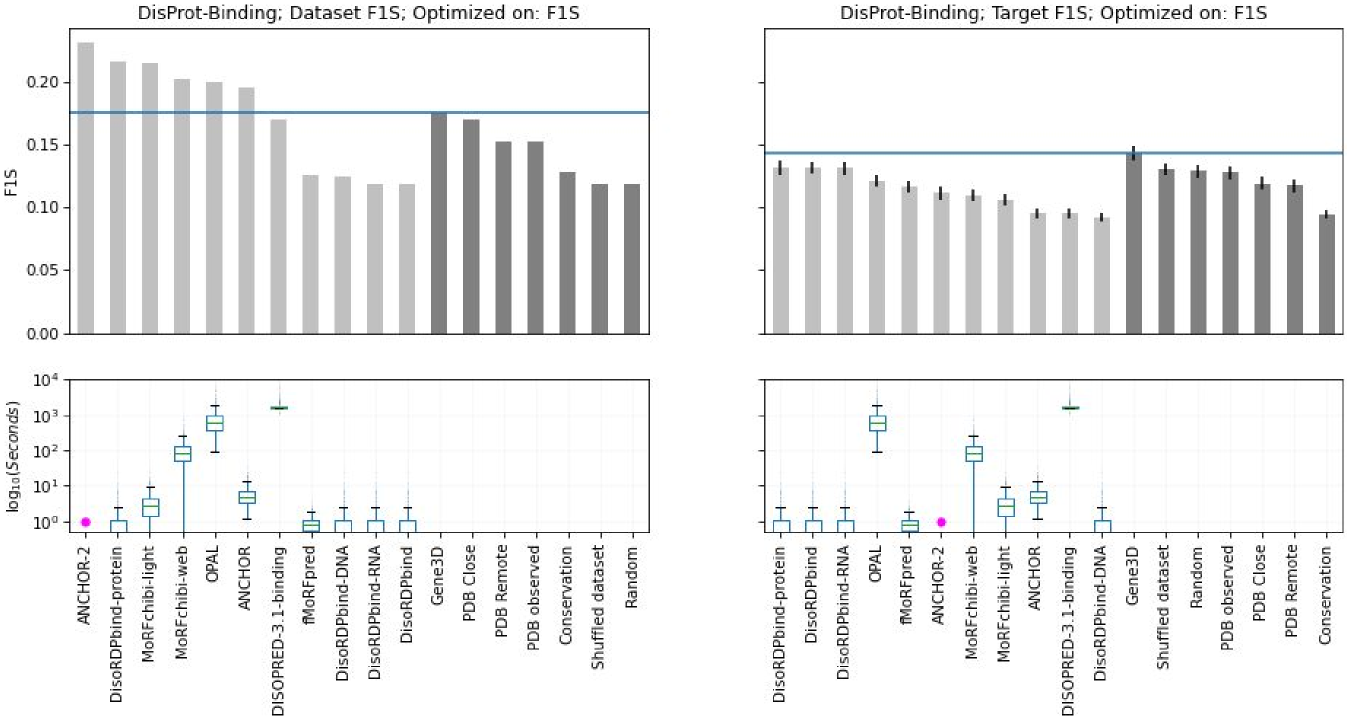


**Supplementary Figure 56**: $F_{Max}$ calculated on the whole dataset with confidence intervals as error bars (left) and averaged over proteins with Standard-Error as error bars (right). Calculated on *DisProt-Binding* dataset. Magenta dots indicate that the whole distribution of execution-times is lower than 1 second.

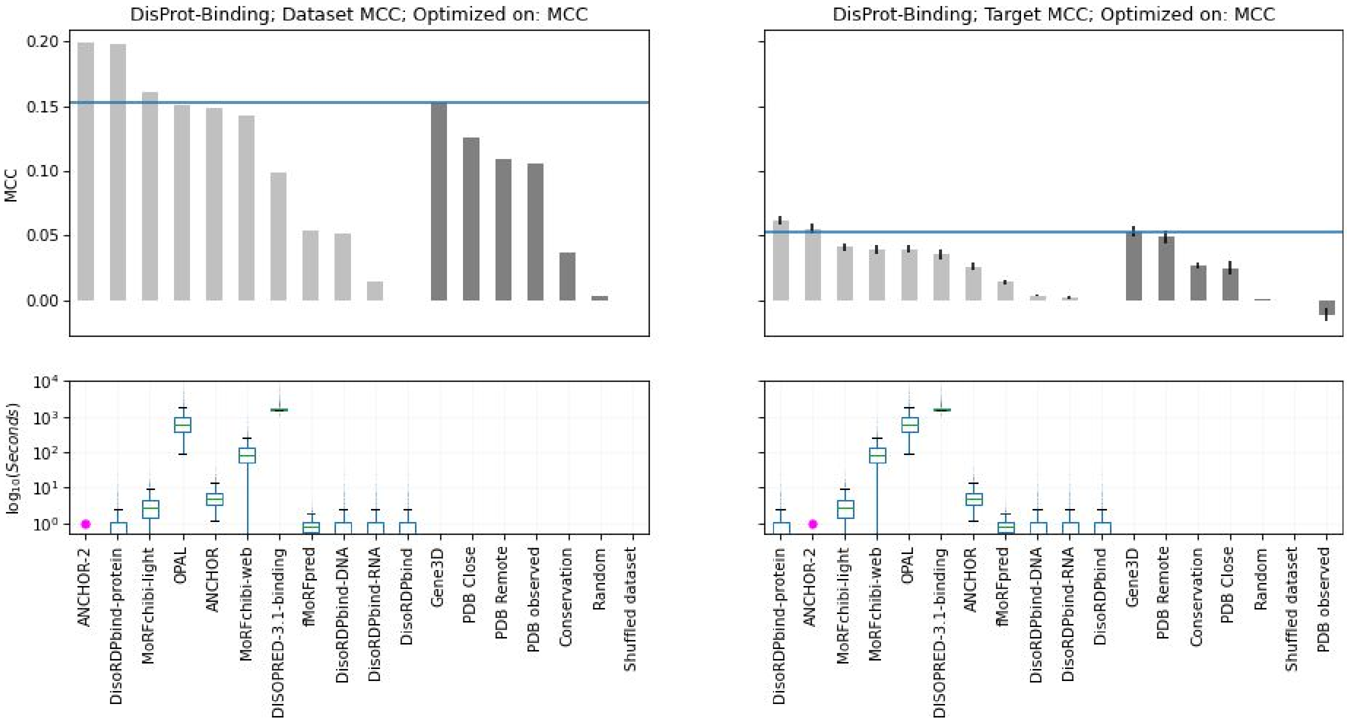


**Supplementary Figure 57**: MCC calculated on the whole dataset with confidence intervals as error bars (left) and averaged over proteins with Standard-Error as error bars (right). Calculated on *DisProt-Binding* dataset. Predictors threshold is optimized on MCC. Magenta dots indicate that the whole distribution of execution-times is lower than 1 second.

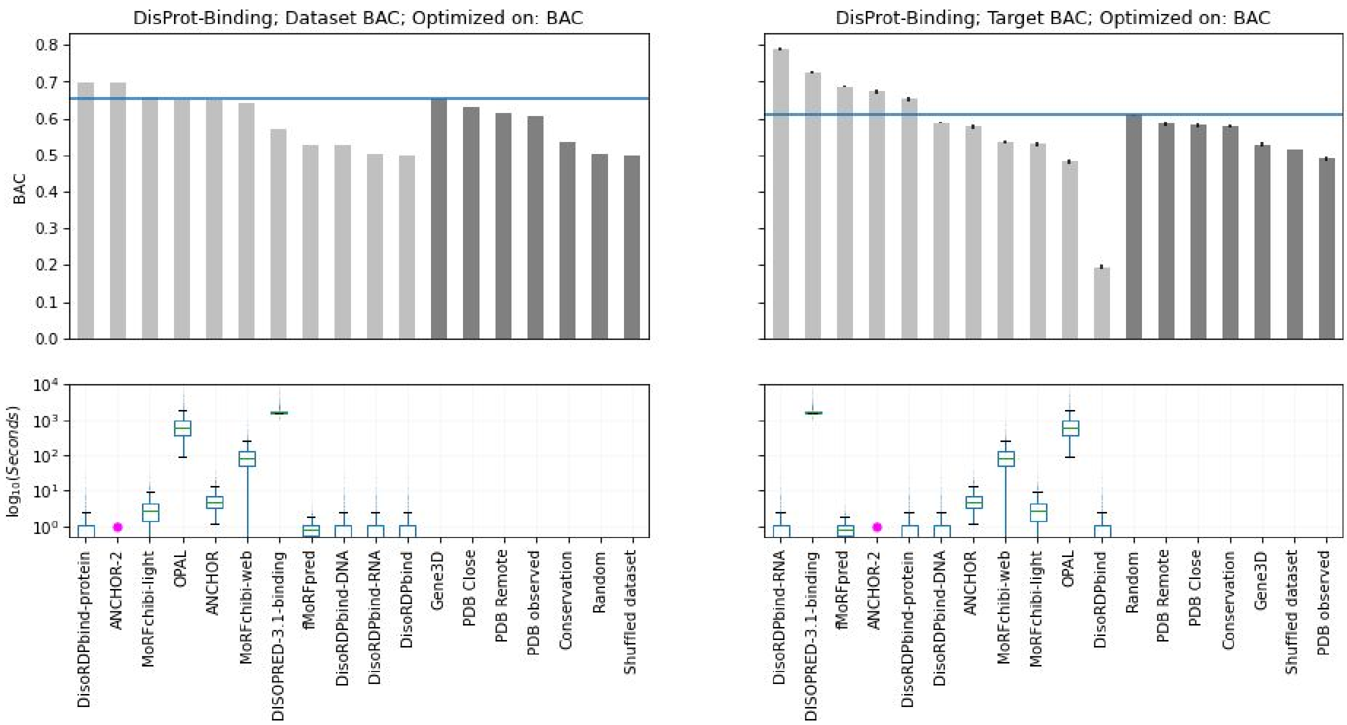


**Supplementary Figure 58**: Balanced accuracy calculated on the whole dataset with confidence intervals as error bars (left) and averaged over proteins with Standard-Error as error bars (right). Calculated on *DisProt-Binding* dataset. Predictors threshold is optimized on Balanced accuracy. Magenta dots indicate that the whole distribution of execution-times is lower than 1 second.

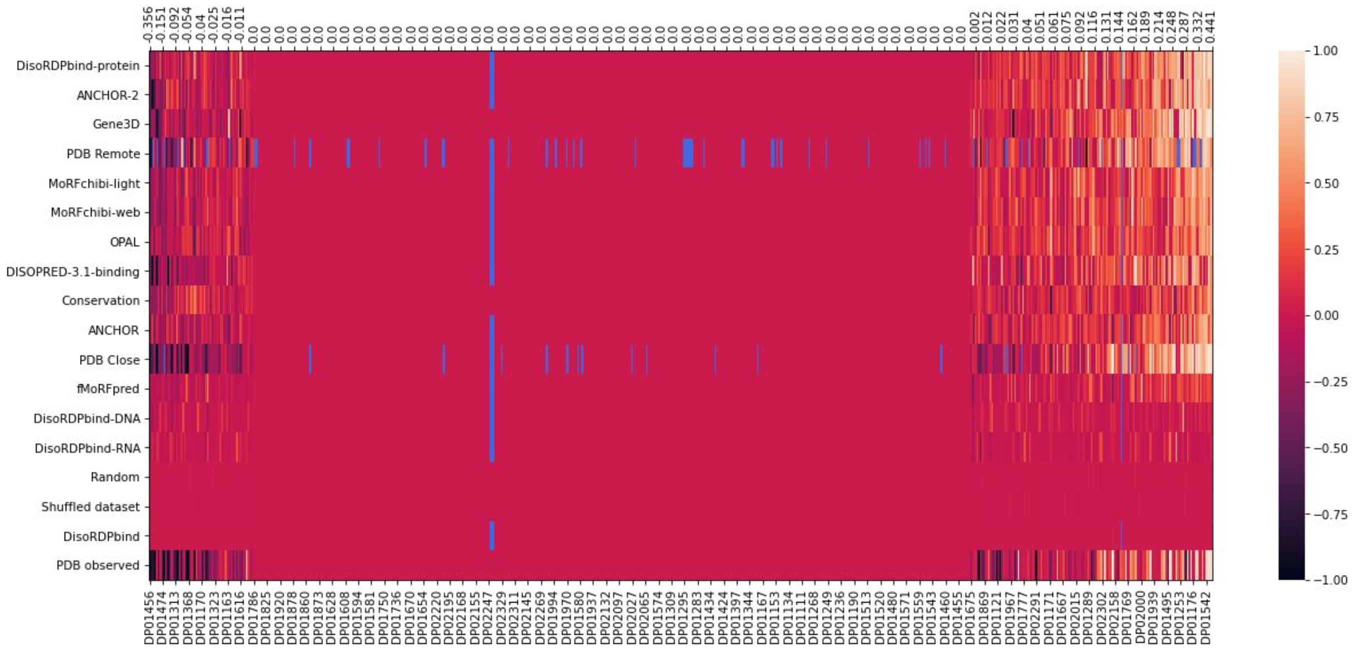


**Supplementary Figure 59**: MCC of each target (x-axis, bottom labels, not all labels are visible) from each predictor (y-axis). Targets are sorted by average MCC (x-axis, top labels). Calculated on *DisProt-Binding*. Predictors threshold is optimized on MCC. Missing values are in blue.

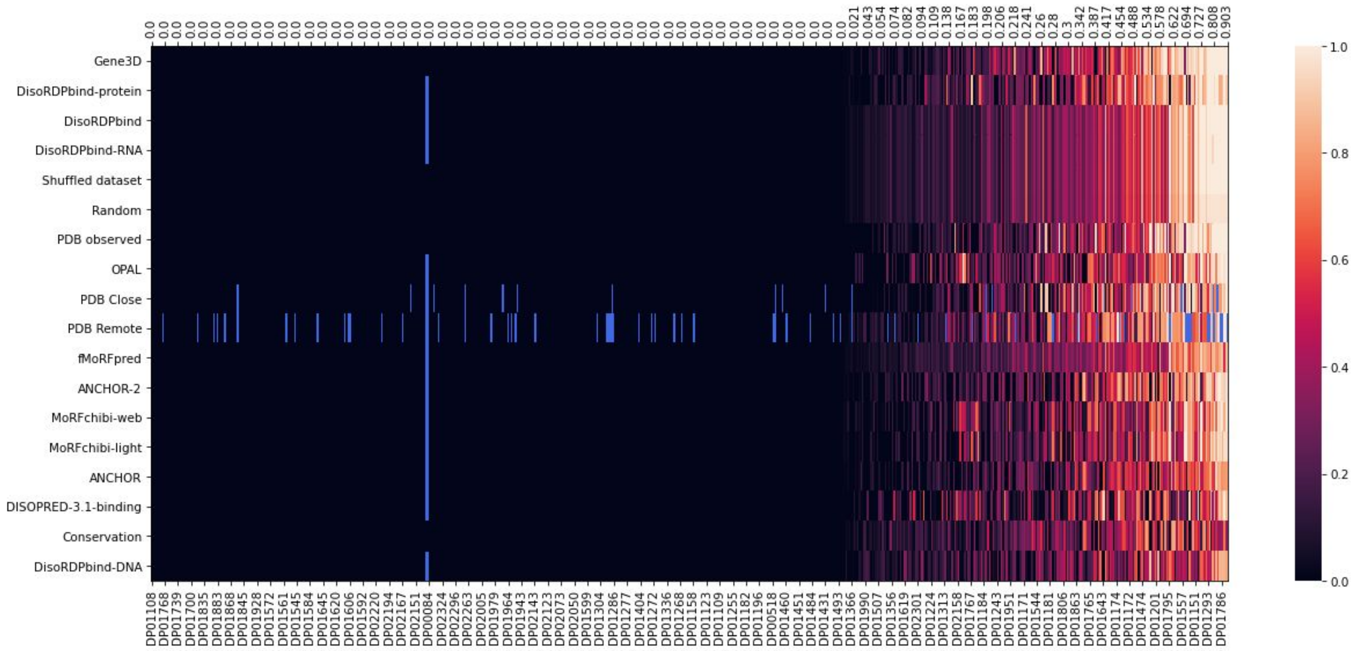


**Supplementary Figure 60**: $F_{Max}$ of each target (x-axis, bottom labels, not all labels are visible) from each predictor (y-axis). Targets are sorted by average $F_{Max}$ (x-axis, top labels). Calculated on *DisProt-Binding* dataset. Missing values are in blue.

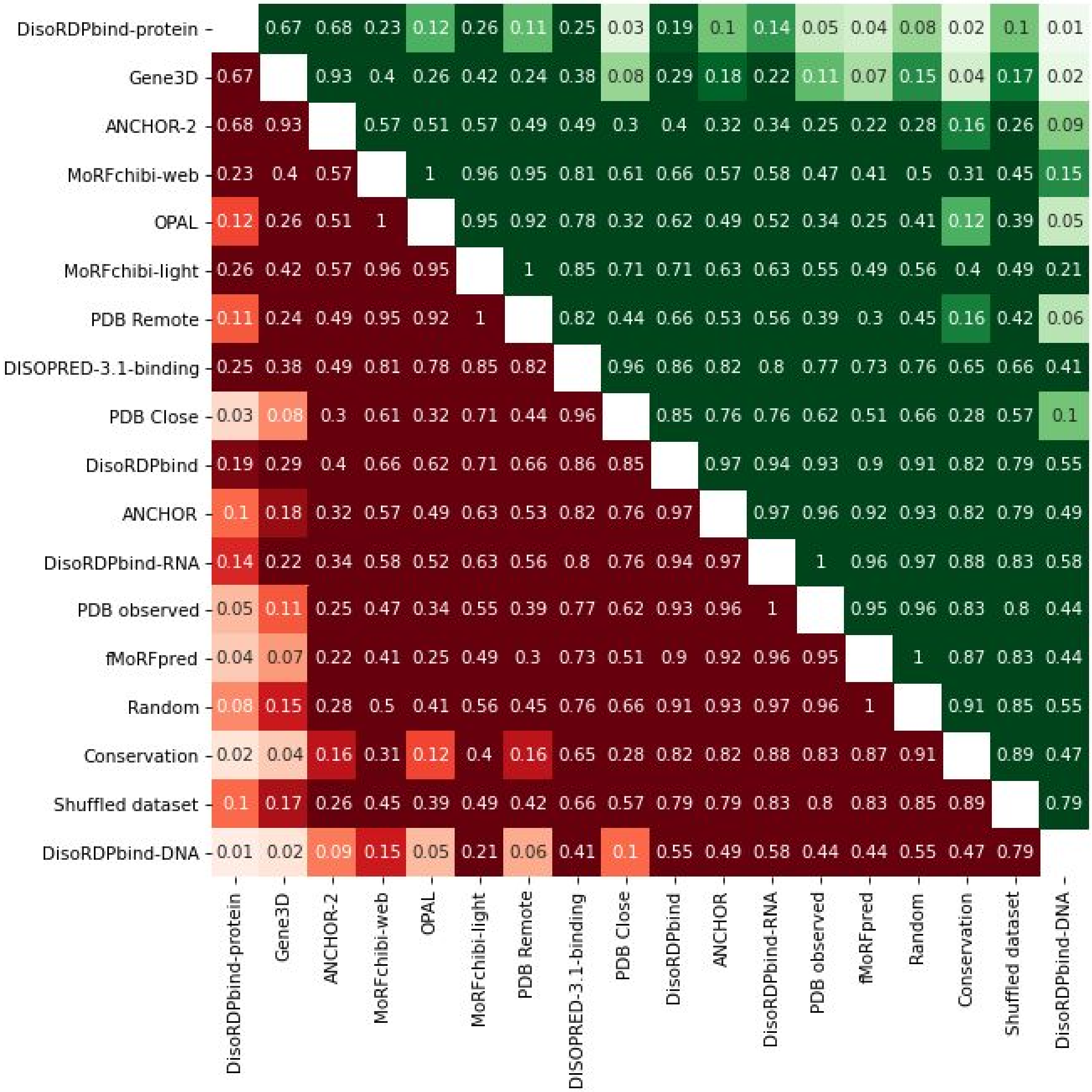


**Supplementary Figure 61**. Overall average ranking of all predictors and baselines. Heatmap of the T-test p-value associated to the statistical significance of the difference between ranking distribution of predictors. A ranking distribution for a predictor is the position of that predictor in its ranking for each metric. Metrics used are: bac, f1s, fpr, mcc, ppv, tpr, tnr; they are calculated with predictors threshold optimized by F1-Score.

Mammals

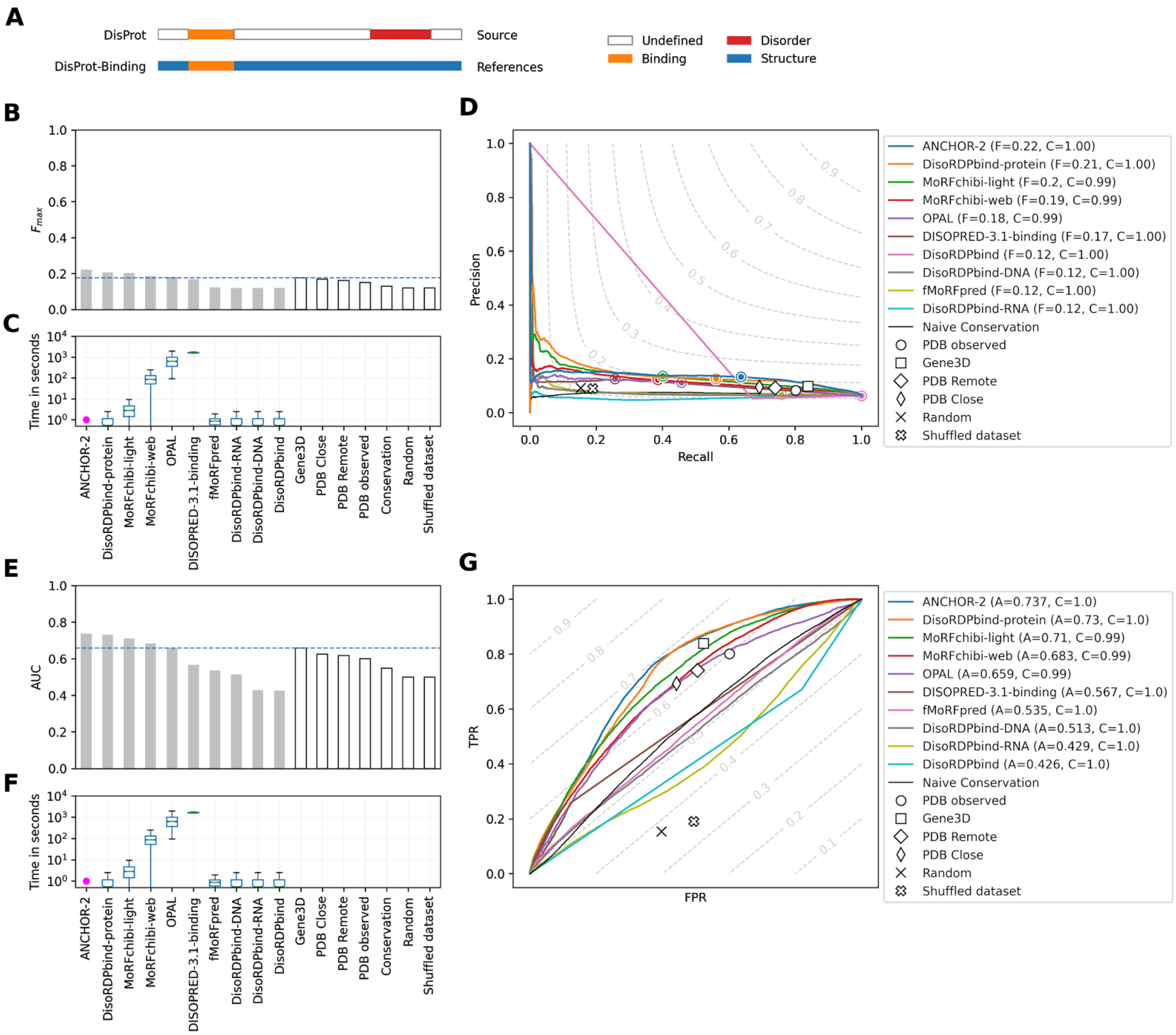


**Supplementary Figure 62.** Prediction success and CPU times for the ten top-ranking disorder predictors for mammalian proteins in the *DisProt-Binding* dataset. Reference used (*DisProt-Binding*) in the analysis and how it is obtained (panel A). Performance of predictors expressed as maximum F1-Score across all thresholds ($F_{max}$) (panel B) and AUC (panel E) for the top ten best ranking methods (light gray) and baselines (white) and the distribution of execution time per-target (panels C, F) using *DisProt-Binding* dataset. The horizontal line in panels B, E indicates the $F_{max}$ and AUC of the best baseline, respectively. Precision-Recall (panel D) and ROC curves (panel G) of ten top-ranking methods and baselines using *DisProt-Binding* dataset, with level curves of the F1-Score and Balanced accuracy, respectively. Magenta dots on panels C, F indicate that the whole distribution of execution-times is lower than 1 second.

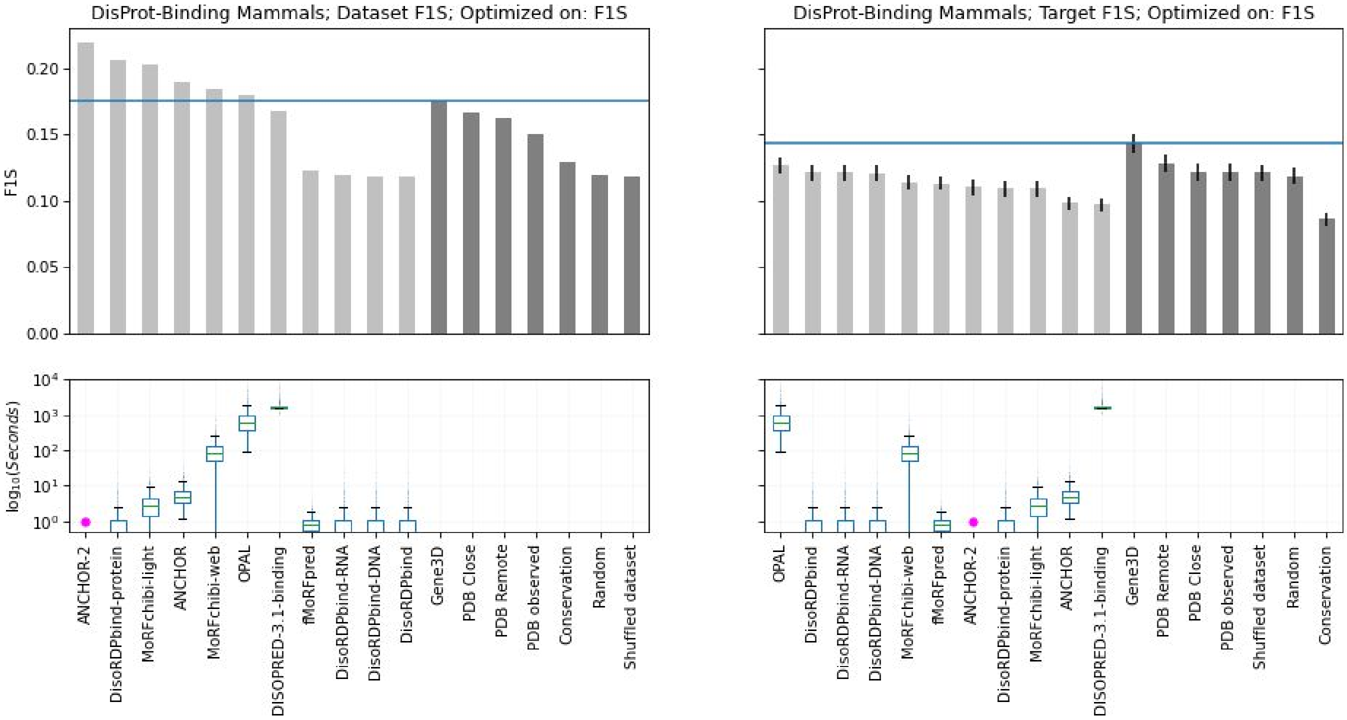


**Supplementary Figure 63**: $F_{Max}$ calculated on the whole dataset with confidence intervals as error bars (left) and averaged over proteins with Standard-Error as error bars (right). Calculated on mammalian proteins of the *DisProt-Binding* dataset. Magenta dots indicate that the whole distribution of execution-times is lower than 1 second.

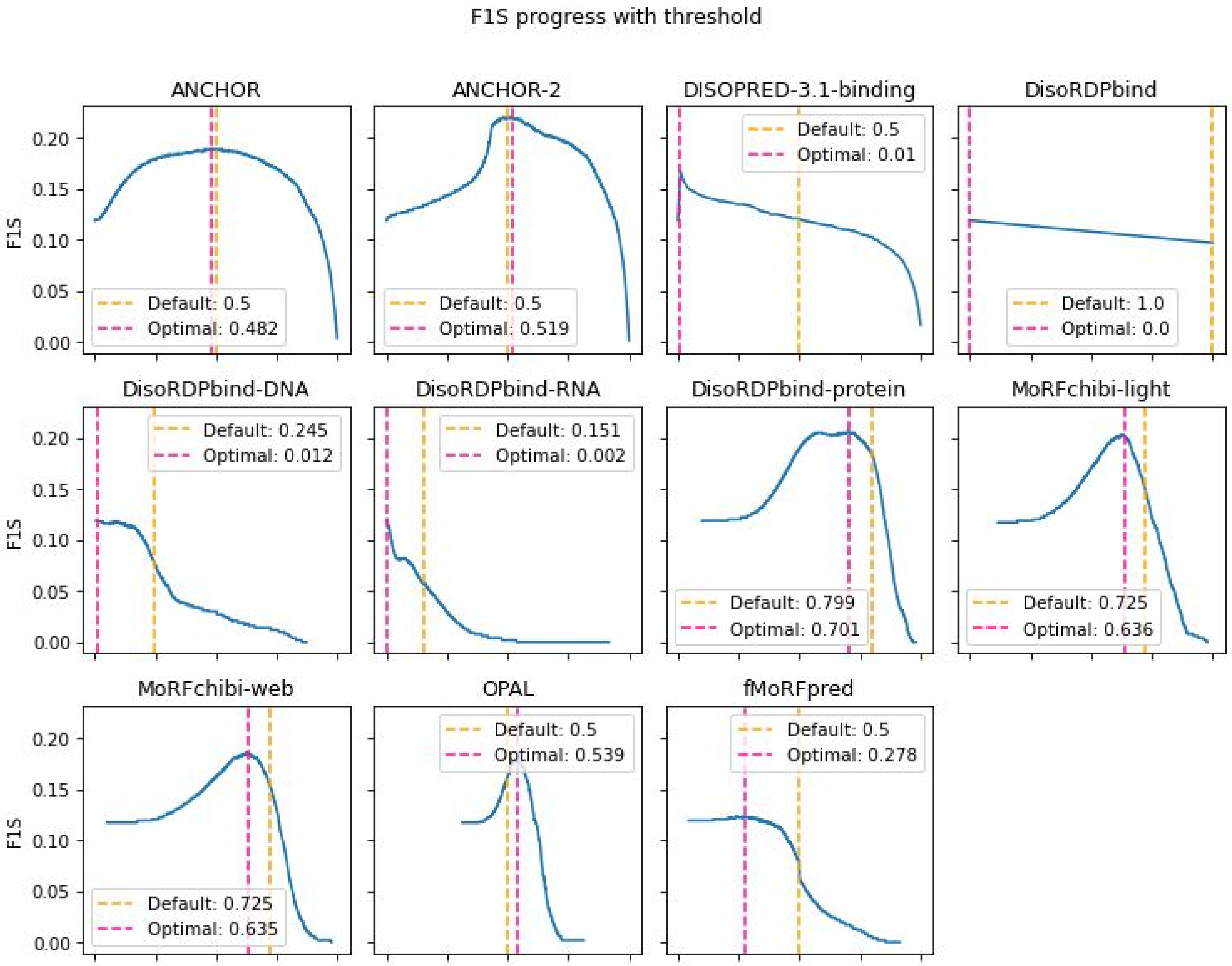


**Supplementary Figure 64**. F1-score progress (y-axis) with increasing threshold value (x-axis) for each predictor calculated on mammalian proteins on the *DisProt-Binding* dataset.

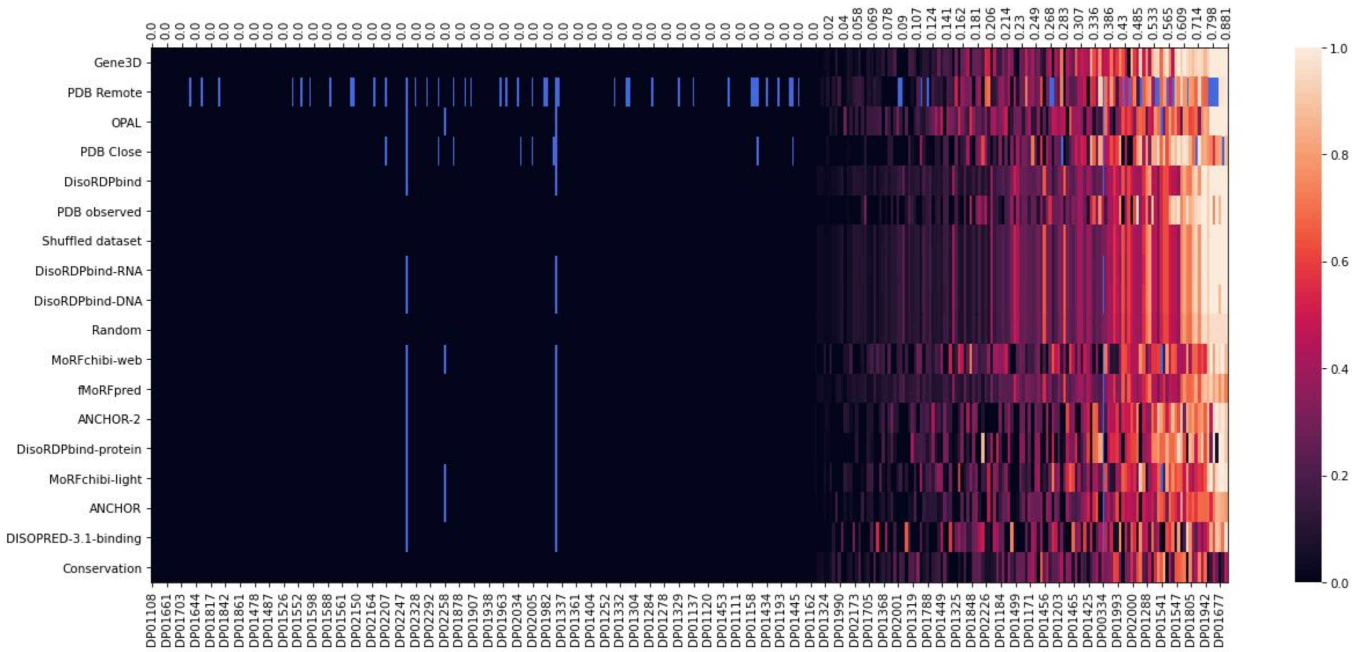

**Supplementary Figure 65**: $F_{Max}$ of each target (x-axis, bottom labels, not all labels are visible) from each predictor (y-axis). Targets are sorted by average $F_{Max}$ (x-axis, top labels). Calculated on mammalian proteins of the *DisProt-Binding* dataset. Missing values are in blue.

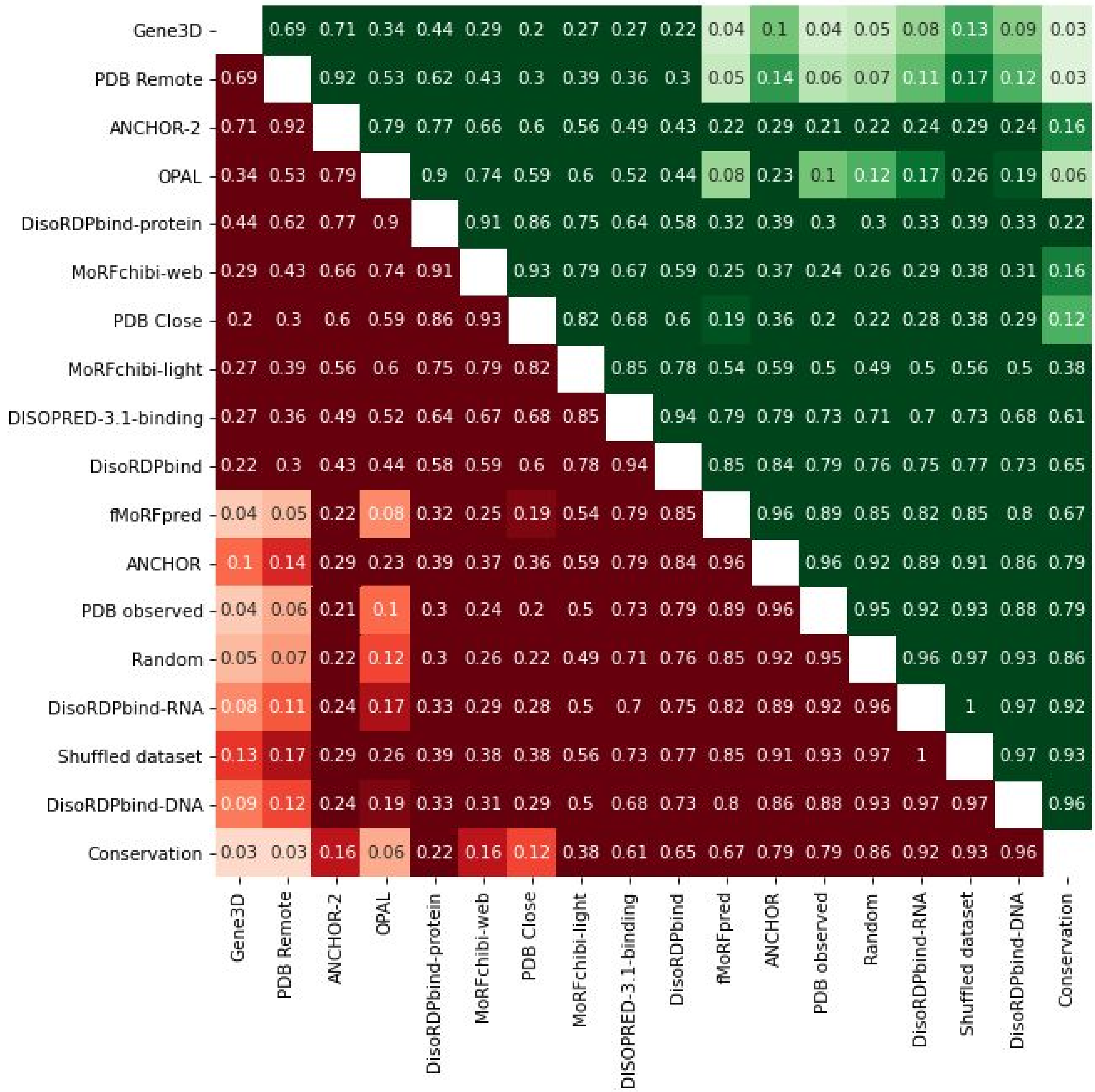


**Supplementary Figure 66**. Overall average ranking of all predictors and baselines for mammalian proteins in the *DisProt-Binding* dataset. Heatmap of the T-test p-value associated to the statistical significance of the difference between ranking distribution of predictors. A ranking distribution for a predictor is the position of that predictor in its ranking for the following metrics: 'bac', 'f1s', 'fpr', 'mcc', 'ppv', 'tpr', 'tnr'. Metrics are calculated per target and with predictors threshold optimized by F1-Score.

#### Prokaryotes

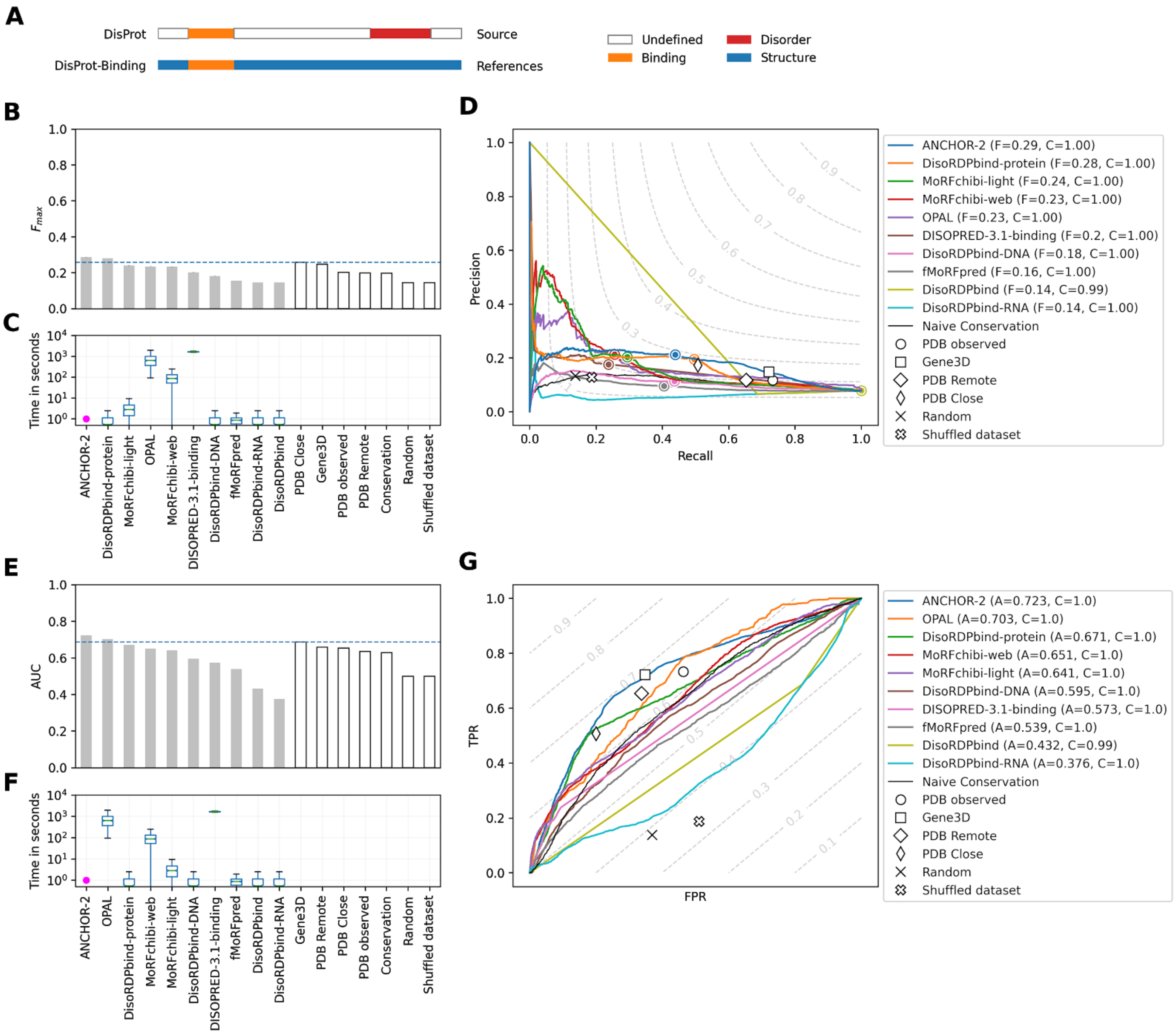


**Supplementary Figure 67.** Prediction success and CPU times for the ten top-ranking disorder predictors for prokaryotic proteins in the *DisProt-Binding* dataset. Reference used (*DisProt-Binding*) in the analysis and how it is obtained (panel A). Performance of predictors expressed as maximum F1-Score across all thresholds ($F_{max}$) (panel B) and AUC (panel E) for the top ten best ranking methods (light gray) and baselines (white) and the distribution of execution time per-target (panels C, F) using *DisProt-Binding* dataset. The horizontal line in panels B, E indicates the $F_{max}$ and AUC of the best baseline, respectively. Precision-Recall (panel D) and ROC curves (panel G) of ten top-ranking methods and baselines using *DisProt-Binding* dataset, with level curves of the F1-Score and Balanced accuracy, respectively. Magenta dots on panels C, F indicate that the whole distribution of execution-times is lower than 1 second.

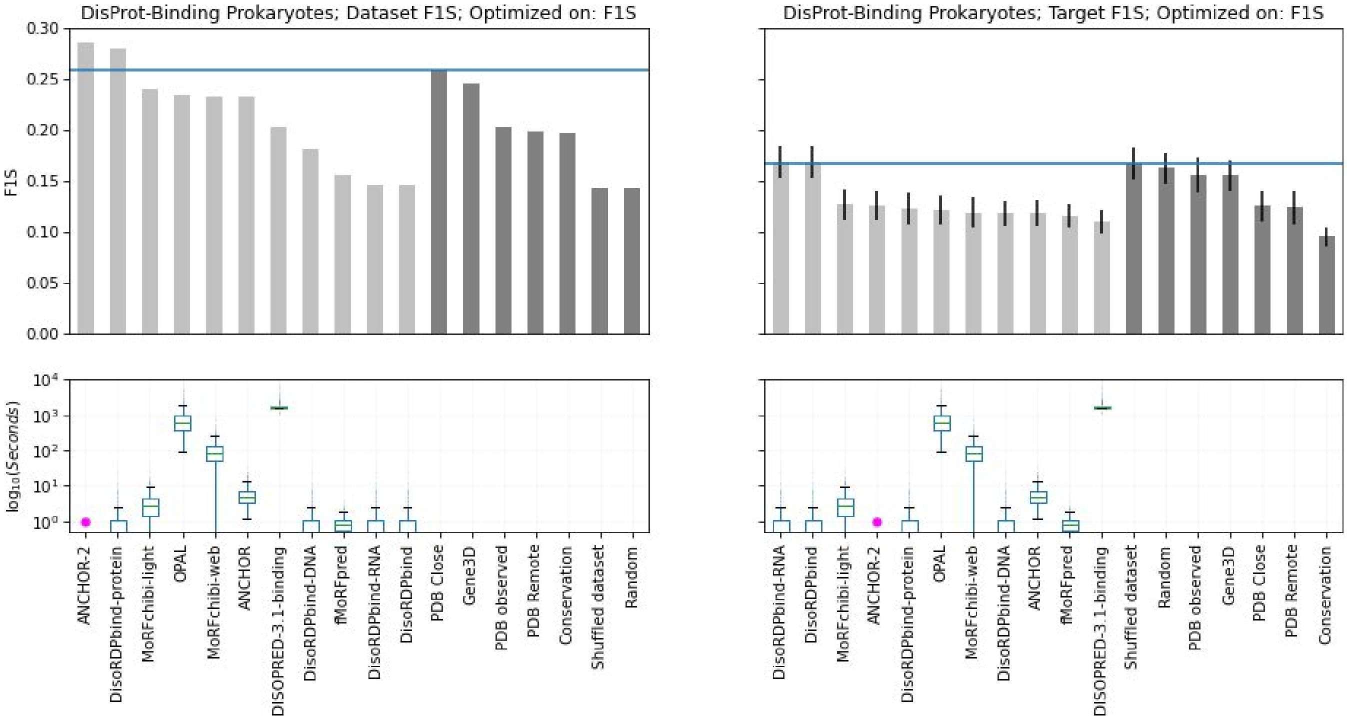


**Supplementary Figure 68**: $F_{Max}$ calculated on the whole dataset with confidence intervals as error bars (left) and averaged over proteins with Standard-Error as error bars (right). Calculated on prokaryotic proteins of the *DisProt-Binding* dataset. Magenta dots indicate that the whole distribution of execution-times is lower than 1 second.

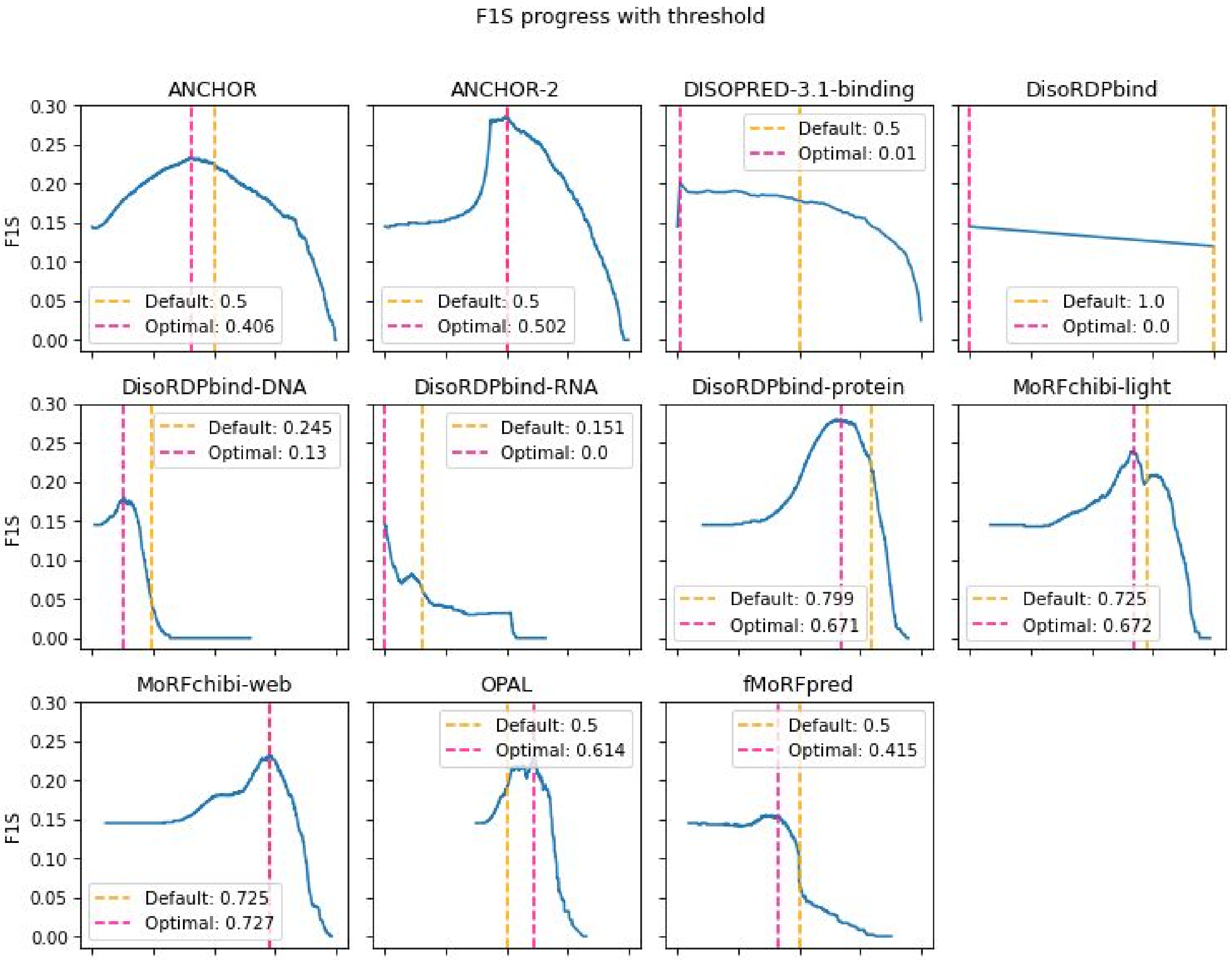


**Supplementary Figure 69**. F1-score progress (y-axis) with increasing threshold value (x-axis) for each predictor calculated on prokaryotic proteins on the *DisProt-Binding* dataset.

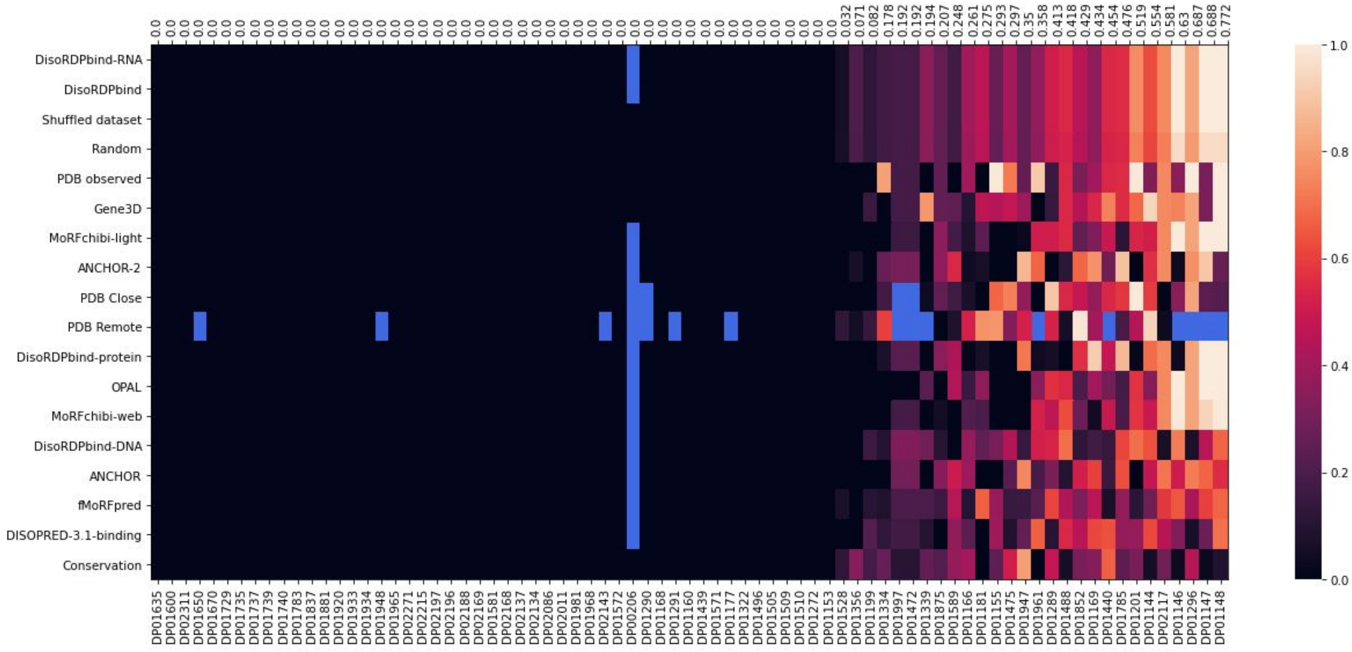

**Supplementary Figure 70**: $F_{Max}$ of each target (x-axis, bottom labels) from each predictor (y-axis). Targets are sorted by average $F_{Max}$ (x-axis, top labels). Calculated on prokaryotic proteins of the *DisProt-Binding* dataset. Missing values are in blue.

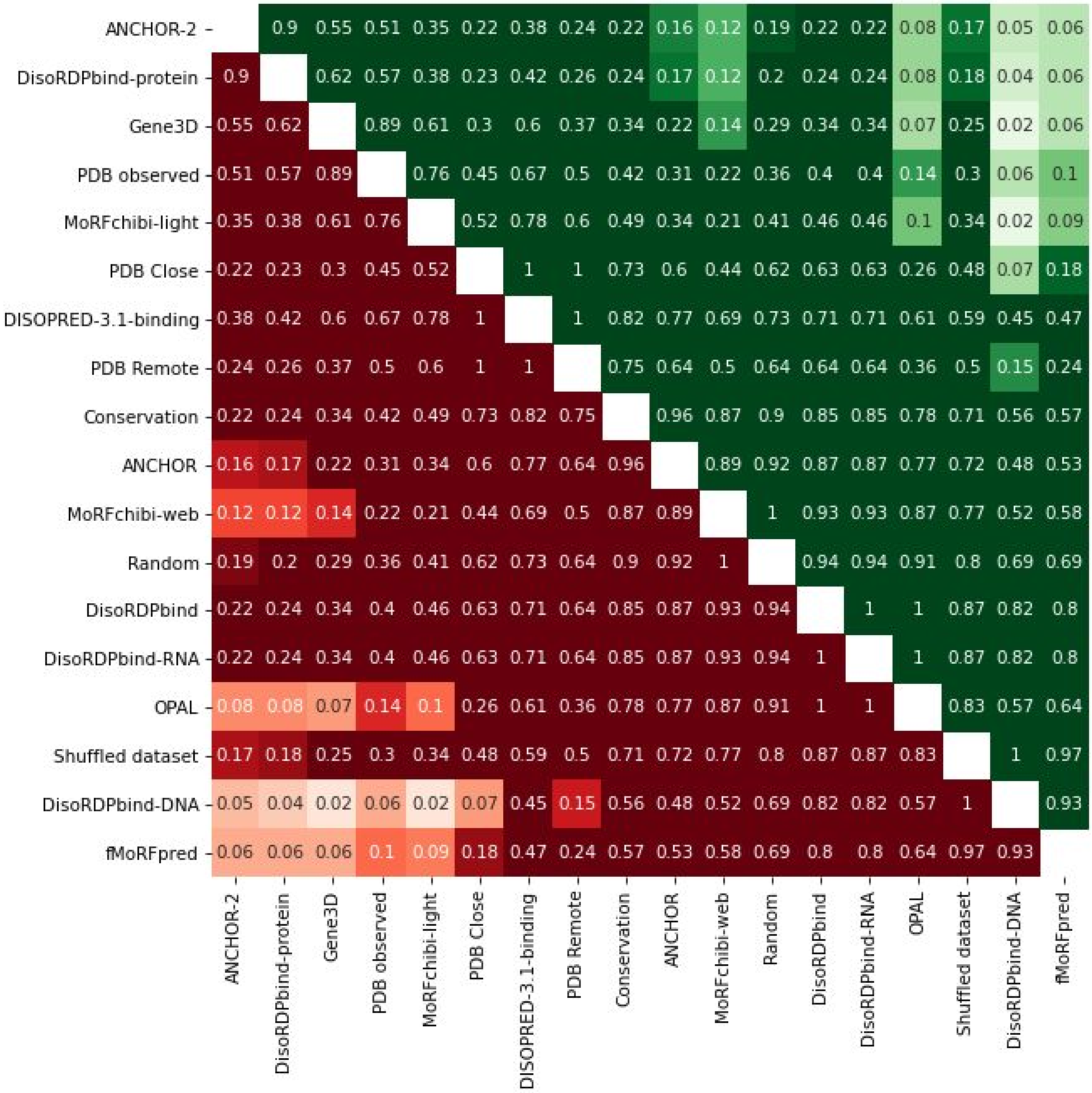


**Supplementary Figure 71**. Overall average ranking of all predictors and baselines for prokaryotic proteins in the *DisProt-Binding* dataset. Heatmap of the T-test p-value associated to the statistical significance of the difference between ranking distribution of predictors. A ranking distribution for a predictor is the position of that predictor in its ranking for the following metrics: 'bac', 'f1s', 'fpr', 'mcc', 'ppv', 'tpr', 'tnr'. Metrics are calculated per target and with predictors threshold optimized by F1-Score.
